# A dual barcoding approach to bacterial strain nomenclature: Genomic taxonomy of *Klebsiella pneumoniae* strains

**DOI:** 10.1101/2021.07.26.453808

**Authors:** Melanie Hennart, Julien Guglielmini, Martin C.J. Maiden, Keith A. Jolley, Alexis Criscuolo, Sylvain Brisse

## Abstract

Sublineages within microbial species can differ widely in their ecology and pathogenicity, and their precise definition is important in basic research and industrial or public health applications. Whereas the classification and naming of prokaryotes is unified at the species level and higher taxonomic ranks, universally accepted definitions of sublineages within species are largely missing, which introduces confusion in population biology and epidemiological surveillance.

Here we propose a broadly applicable genomic classification and nomenclature approach for bacterial strains, using the prominent public health threat *Klebsiella pneumoniae* as a model. Based on a 629-gene core genome multilocus sequence typing (cgMLST) scheme, we devised a dual barcoding system that combines multilevel single linkage (MLSL) clustering and life identification numbers (LIN). Phylogenetic and clustering analyses of >7,000 genome sequences captured population structure discontinuities, which were used to guide the definition of 10 infra-specific genetic dissimilarity thresholds. The widely used 7-gene multilocus sequence typing (MLST) nomenclature was mapped onto MLSL sublineages (threshold: 190 allelic mismatches) and clonal group (threshold: 43) identifiers for backwards nomenclature compatibility. The taxonomy is publicly accessible through a community-curated platform (https://bigsdb.pasteur.fr/klebsiella), which also enables external users’ genomic sequences identification.

The proposed strain taxonomy combines two phylogenetically informative barcodes systems that provide full stability (LIN codes) and nomenclatural continuity with previous nomenclature (MLSL). This species-specific dual barcoding strategy for the genomic taxonomy of microbial strains is broadly applicable and should contribute to unify global and cross-sector collaborative knowledge on the emergence and microevolution of bacterial pathogens.

## Introduction

Taxonomy is a foundation of biology that entails the classification, nomenclature, and identification of biological objects (Cowan, 1965). Although the Linnaean taxonomy was initially devised for plant and animal organisms, it has been extended to include prokaryotes, which are organized into taxonomic ranks down to the level of species (Sneath, 1992). Because prokaryotes reproduce predominantly by binary fission and clonal reproduction, sublineages within microbial species can diversify as independently evolving lineages that persist over long periods of time (Selander and Levin, 1980). In addition, the broad microbial species definition and horizontal gene transfer of accessory genes underlie extensive strain heterogeneity of phenotypes with ecological, medical or industrial relevance (Hacker and Kaper, 2000; Lan and Reeves, 2001; Feil, 2004; Konstantinidis and Tiedje, 2005), which is mostly overlooked by current prokaryotic taxonomy. The subspecies taxonomic rank is rarely used and the large number of distinguishable sublineages in most bacterial species renders the applicability of Latin binomial identifiers impractical.

Most attempts to develop and maintain microbial strains taxonomies aimed at facilitating epidemiological surveillance and outbreak detection (Maiden et al., 1998; van Belkum et al., 2007; Maiden et al., 2013). Although local epidemiology can rely on vernacular type designations, the benefits of unified nomenclatures of sublineages for large-scale epidemiology and population biology were recognized early (Struelens et al., 1998). By far the most successful taxonomic system of microbial strains is the multilocus sequence typing (MLST) approach (Maiden et al., 1998; Achtman et al., 2012). This highly reproducible and portable nomenclature system has been extensively used for studies of population biology and public health surveillance, to the extent that dedicated MLST nomenclatural systems exist for all important pathogenic bacterial species (Jolley et al., 2018). Core genome MLST (cgMLST) extends the advantages of the MLST approach at the genomic scale (Jolley and Maiden, 2010; Maiden et al., 2013) and provides strain discrimination at much finer scales.

Using cgMLST, strain classification can be achieved by grouping allelic profiles by similarity, and several clustering thresholds can be used simultaneously, leading to a succession of group identifiers (‘barcodes’) that provide relatedness information at increasing levels of phylogenetic depth (Maiden et al., 2013; Moura et al., 2016). This approach was recently formalized as hierarchical clustering (Zhou et al., 2021) or the ‘single nucleotide polymorphism (SNP) address’ (Dallman et al., 2018). Unfortunately, the single linkage clustering approach suffers from instability, due to the possible fusion of preexisting groups as additional genomes are introduced.

An alternative approach, the Life Identification Number (LIN) codes, was proposed by Vinatzer and colleagues (Marakeby et al., 2014; Weisberg et al., 2015; Vinatzer et al., 2016, 2017; Tian et al., 2020): a multi-position numerical code is assigned to each genome based on its similarity with the closest genome already encoded. An attractive property of this approach is that LIN codes are definitive, *i.e*. not affected by subsequent additions of genomes, as they are attributed to individual genomic sequences rather than to groups. However, in the current implementation of LIN codes the similarity between genomes is estimated using Average Nucleotide Identity (ANI), which may be imprecise for nearly identical strains.

Here, we present a strain classification, naming and identification system for bacterial strains, which is based on cgMLST and combines the MLSL and LIN code approaches. We took as a model the *Klebsiella pneumoniae* species complex, a genetically and ecologically highly diverse bacterial group that causes a wide range of infections in humans and animals (Brisse et al., 2006; Wyres et al., 2020a). Given its extensive diversity and fast evolutionary dynamics, *K. pneumoniae* is a challenging model for the development of a genomic taxonomy of strains, but the rapid emergence and global dissemination of multidrug resistance in *K. pneumoniae*, sometimes combined with high virulence (Bialek-Davenet et al., 2014; Wyres et al., 2020b) have created a pressing need for an efficient *K. pneumoniae* strain definition and tracking system.

## Results

### Genome-based phylogenetic structure of the K. pneumoniae species complex and 7-gene MLST classification

The deep phylogenetic structure of the *K. pneumoniae* (Kp) species complex (**Figure 1**) reflects the previously recognized seven major phylogroups, Kp1 to Kp7 (Brisse and Verhoef, 2001; Fevre et al., 2005; Blin et al., 2017; Long et al., 2017; Rodrigues et al., 2019; Wyres et al., 2020a). The most represented phylogroup (91.7%; n=6,476) is Kp1, *i.e*., *K. pneumoniae sensu stricto* (**Table 1**), and its phylogenetic structure (**Figure 2**) revealed a multitude of sublineages. There were multiple closely-related isolates within some sublineages, most prominently within a sublineage comprising genomes with 7-gene MLST identifiers ST258, ST11, ST512 (**Figure 2**), which represented more than a third (33.4%) of the Kp1 dataset. The abundance of this sublineage and a few others, such as ST23, reflected the clinical microbiology focus on multidrug resistant or hypervirulent isolates (Bowers et al., 2015; Struve et al., 2015; Lam et al., 2018; Wyres et al., 2020b). The phylogenetic structure within other *K. pneumoniae* phylogroups also revealed a multitude of distinct sublineages but no predominant ones, and medically important lineages in these phylogroups are yet to be recognized.

**Figure 1.**
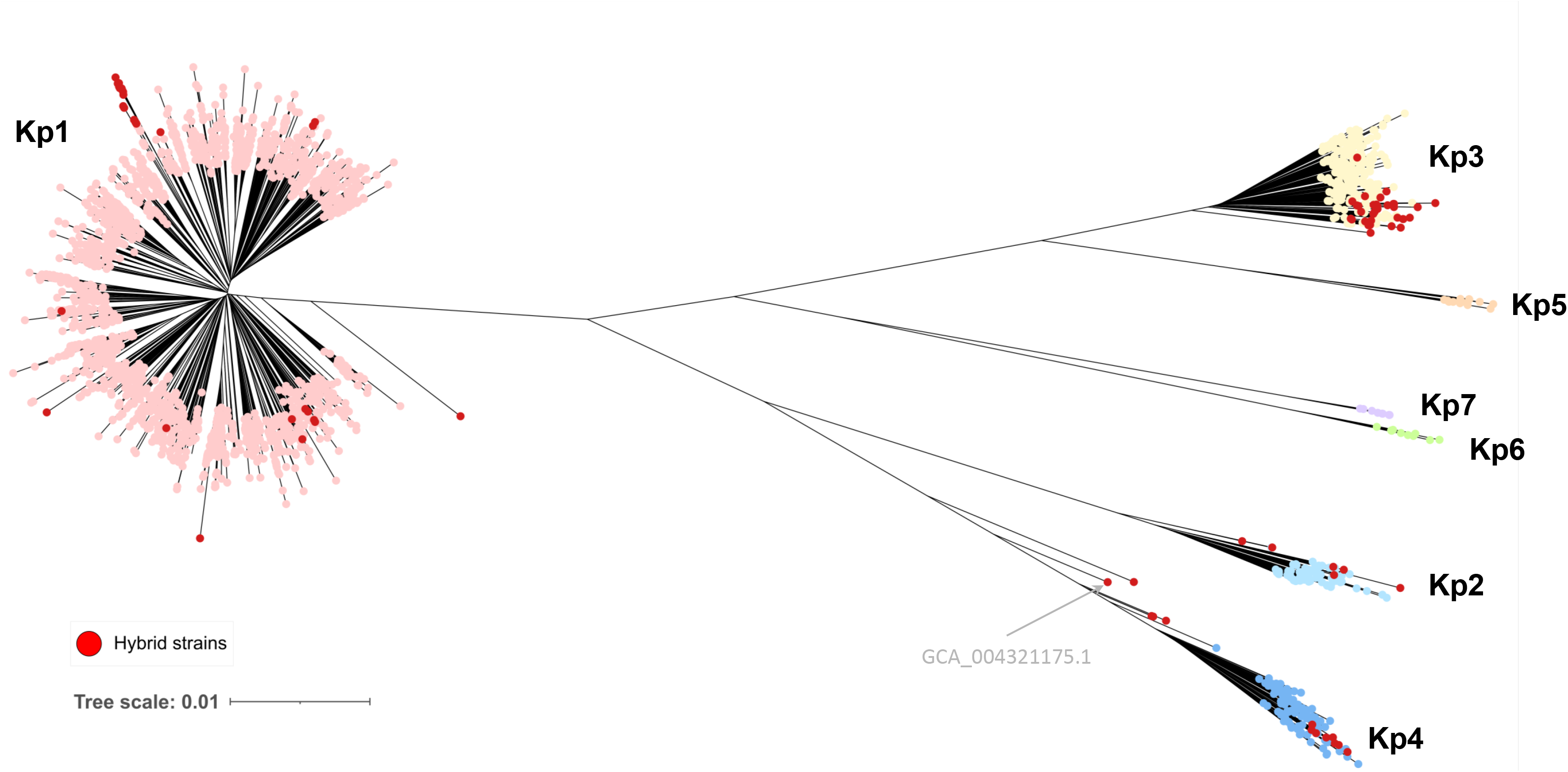
Genome-based phylogenetic tree of the *K. pneumoniae* complex. The distance-based tree was inferred using JolyTree. The seven phylogroups are indicated. Red dots correspond to strains defined as inter-phylogroup hybrids. Scale bar, 0.01 nucleotide substitutions per site.

**Figure 2.**
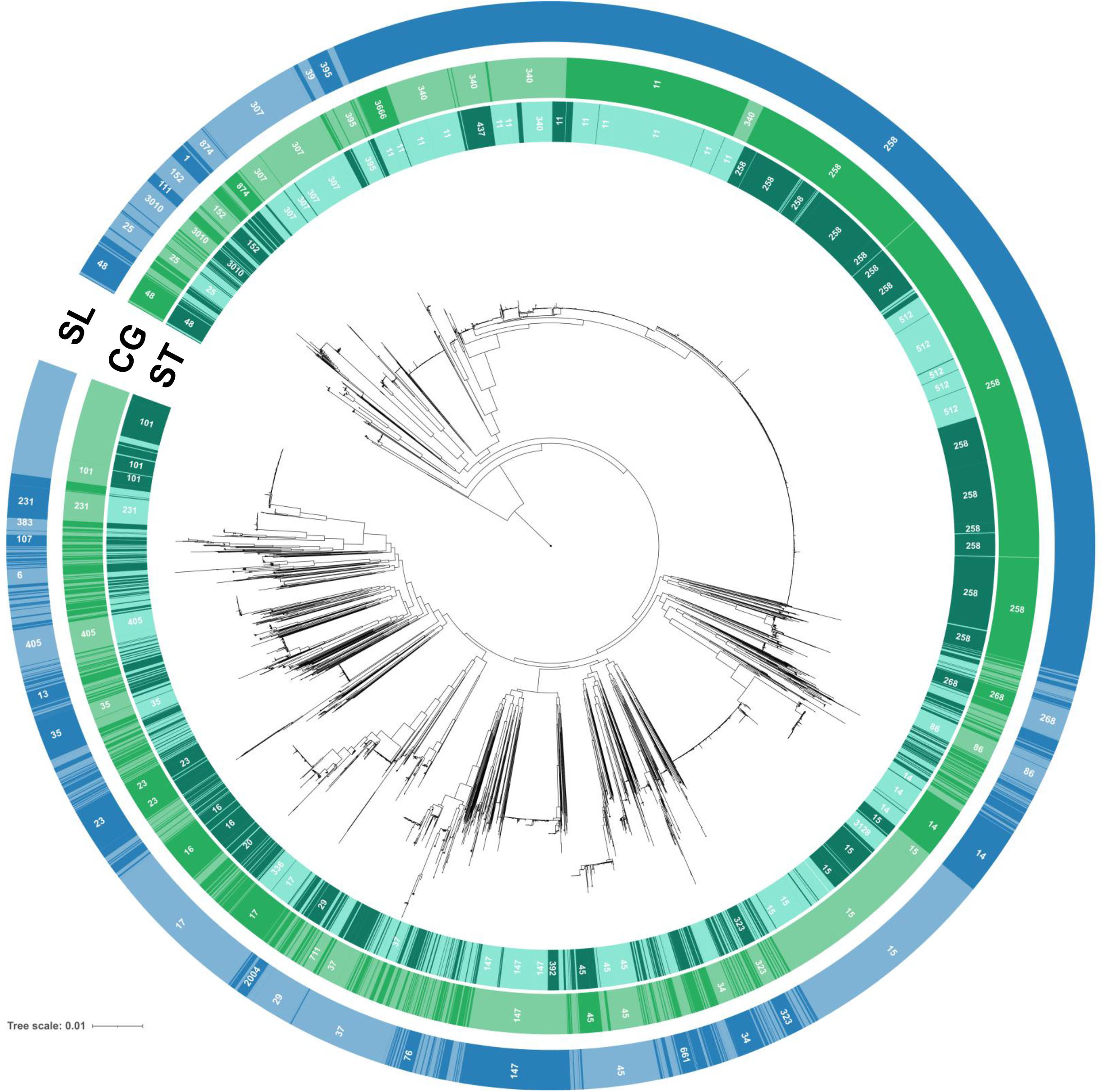
Phylogenetic structure within phylogroup Kp1 (*K. pneumoniae* sensu stricto). The circular tree was obtained using IQ-TREE based on the concatenation of the genes of the scgMLSTv2 scheme; 1,600 isolates are included (see Methods). Labels on the external first circle represent 7-gene MLST ST numbers (each alternation corresponds to a different ST and only ST with more than 20 strains are labelled). The second and third circles (light green and blue, respectively) show the alternation of clonal groups (CG) and sublineages (SL), respectively, labelling only groups with more than 20 isolates. Full correspondence between ST, SL and CG identifiers is given in the supplementary appendix.

*K. pneumoniae* strains can recombine large sections of their chromosome (Chen et al., 2014; Wyres et al., 2015). Large recombination events were detected in 1.9% (138) genomes and involved the phylogroups Kp1, Kp2 and Kp4 (supplementary appendix: Detection of hybrids**; Figure S1; Table S1; Table S2**). The phylogenetic impact of large-scale recombination is illustrated on **Figure 1**, with ‘hybrids’ occurring on atypically long branches.

### cgMLST analysis of the K. pneumoniae species complex

A previously defined strict core genome MLST (scgMLST) scheme (Bialek-Davenet et al., 2014) was updated (**Table S3**) and defined as scgMLSTv2. cgMLST allelic profiles were then determined for 7,433 genomic sequences (including 45 reference sequences; **Figure S2**). The mean number of missing alleles per profile was 8 (1.2%; standard deviation: 25; 4.0%), and most (7,198; 96.8%) isolates had a cgMLST profile with fewer than 30 (4.8%) missing alleles. Missing allele proportions did not vary significantly among phylogroups (**Table 1**). The transcription-repair coupling factor *mfd* gene was atypical, with 778 alleles and an average allele size of 3,447 nt; for the other loci, the number of distinct alleles varied from 8 to 626 (median: 243), and was strongly associated with locus size (range: 123 to 2,826 nt; median: 758 nt); (**Figure S3**). Locus-by-locus recombination analyses detected evidence of intra-gene recombination (PHI test; 5% *p*-value significance) in half of the loci (318/629; 50.6%) and these exhibited more alleles than non-recombining ones (**Table S3; Figure S3**).

The distribution of pairwise allelic mismatch proportions among non-hybrid cgMLST allelic profiles was discontinuous (**Figure 3**), with four major modes centered around values 99.7% (627 mismatches), 82.7% (520 mismatches), 12.4% (79 mismatches) and 2.0% (13 mismatches). ANI-based similarity values (**Figure S4**) varied from 92.8% to 100%, with two first modes at 93.5% and 95.5%, composed of inter-phylogroup strain comparisons. The corresponding genome pairs typically had only ~2% cgMLST similarity. In turn, whereas the range of ANI values was only 98% to 100% for intra-species pairs, their cgMLST similarities occupied the much broader 5%-100% range.

**Figure 3.**
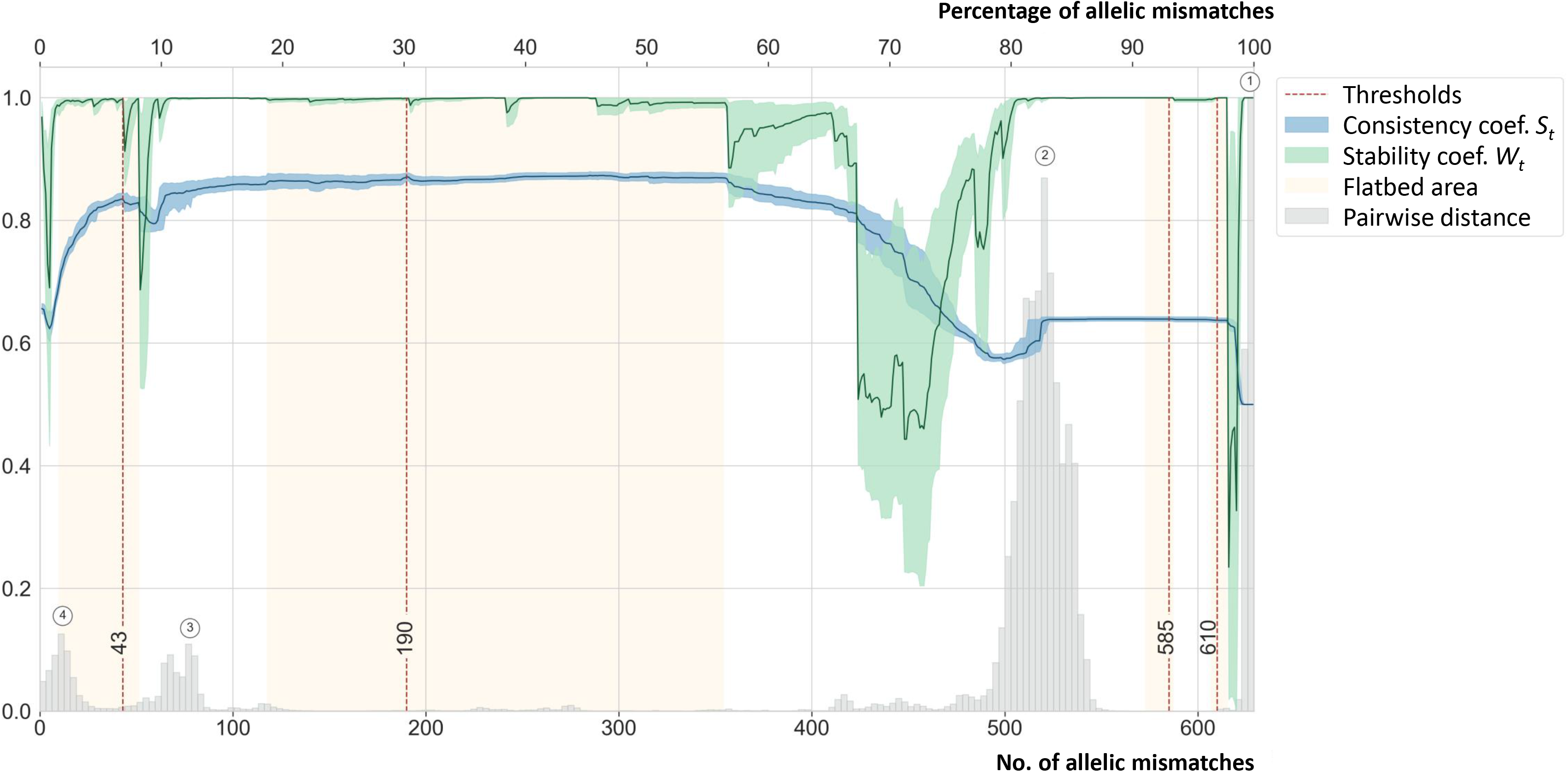
Distribution of pairwise cgMLST distances, clustering properties and phylogenetic congruence. Values are plotted for the 7,060 genomes dataset. Threshold values (*t*) are shown on the X-axis, corresponding to allelic profile mismatch values up to 629 (or 100%). Grey histograms: distribution of pairwise allelic mismatches. The curves represent the consistency (silhouette) and stability coefficients *S_t_* (blue) and *W_t_* (green), respectively, obtained with each threshold *t*; the corresponding scale is on the left Y-axis. The dotted vertical red lines at *t* = 43/629, 190/629, 585/629 and 610/629 represent the thresholds that correspond to clonal groups, sublineages, phylogroups and species, respectively.

The 627-mismatch mode corresponded mostly to pairs of strains belonging to distinct species of the Kp complex (**Figure 3; Figure S4**), while a minor peak centered on 591 mismatches (**Figure S5**) corresponded to comparisons between subspecies of *K. quasipneumoniae* and *K. variicola* (Kp3 and Kp5, and Kp2 and Kp4, respectively; **Figure S4**). Whereas the 520-mismatch mode corresponded to inter-ST comparisons in 99.9% cases, the 16-mismatch mode was largely dominated by comparisons of genomes with the same ST (68.2%; pairs of genomes within 402 distinct STs) or of single-locus variants (SLV; 30.8%). Finally, the 79-mismatch mode comprised a large proportion (48.0%) of ST258-ST11 comparisons and other comparisons of atypically closely-related STs (**Figure S5**).

### Definition of classification thresholds for phylogroups, sublineages and clonal groups

To determine optimal allelic mismatch thresholds that would reflect the Kp population structure, the consistency and stability properties of single-linkage clustering groups were assessed for every threshold value *t* from 1 to 629 allelic mismatches. The Silhouette coefficient *S_t_* had a plateau of optimal values in the range corresponding to 118/629 (18.8%) to 355/629 (56.4%) allelic mismatches (**Figure 3, blue curve**). Analysis of the stability to subsampling (*W_t_*; based on an adjusted Wallace coefficient; **Figure 3, green curve**) identified several ranges of allelic mismatch threshold values that were associated to maximal stability.

The above analyses led us to propose four deep classification levels. The two first thresholds, 610 and 585 allelic mismatches, enable species and subspecies separations, respectively. We next defined a threshold at the level of 190 allelic mismatches, corresponding to the optimal combination of consistency and stability coefficients *S_t_* and *W_t_*. The single-linkage clustering based on this threshold created 705 groups, which we define as ‘sublineages’ (SL). By design, this threshold separated into distinct groups, the pairs of genomes corresponding to the major mode (at 520 mismatches), *i.e*., the majority of genomes that have distinct STs within phylogroups. Finally, a threshold of 43 allelic mismatches was defined to separate genome pairs of the 79-mismatch mode. This value corresponded to local optima of both *S_t_* and *W_t_* coefficients. Interestingly, this threshold value was also located in the optimal range of compatibility with the classical 7-gene ST definitions (Rand index *R_t_* ≥ 0.70 was observed for 10 ≤ *t* ≤ 51). The use of this threshold resulted in 1,147 groups, which we propose to define as ‘clonal groups’ (CG).

Overall, approximately half (547/1,147; 47.7%) of the CGs corresponded one-to-one with the sublineage level (**Table S4**): 77.6% (547/705) sublineages contained a single CG, whereas 158 (22.4%) sublineages comprised at least two clonal groups (**Table S5; Figure 4; Figure S6**). Overall, CG compatibility with classical ST classification was high (*i.e*., *R_t_* = 0.72, whereas it was only 0.50 for sublineages).

**Figure 4.**
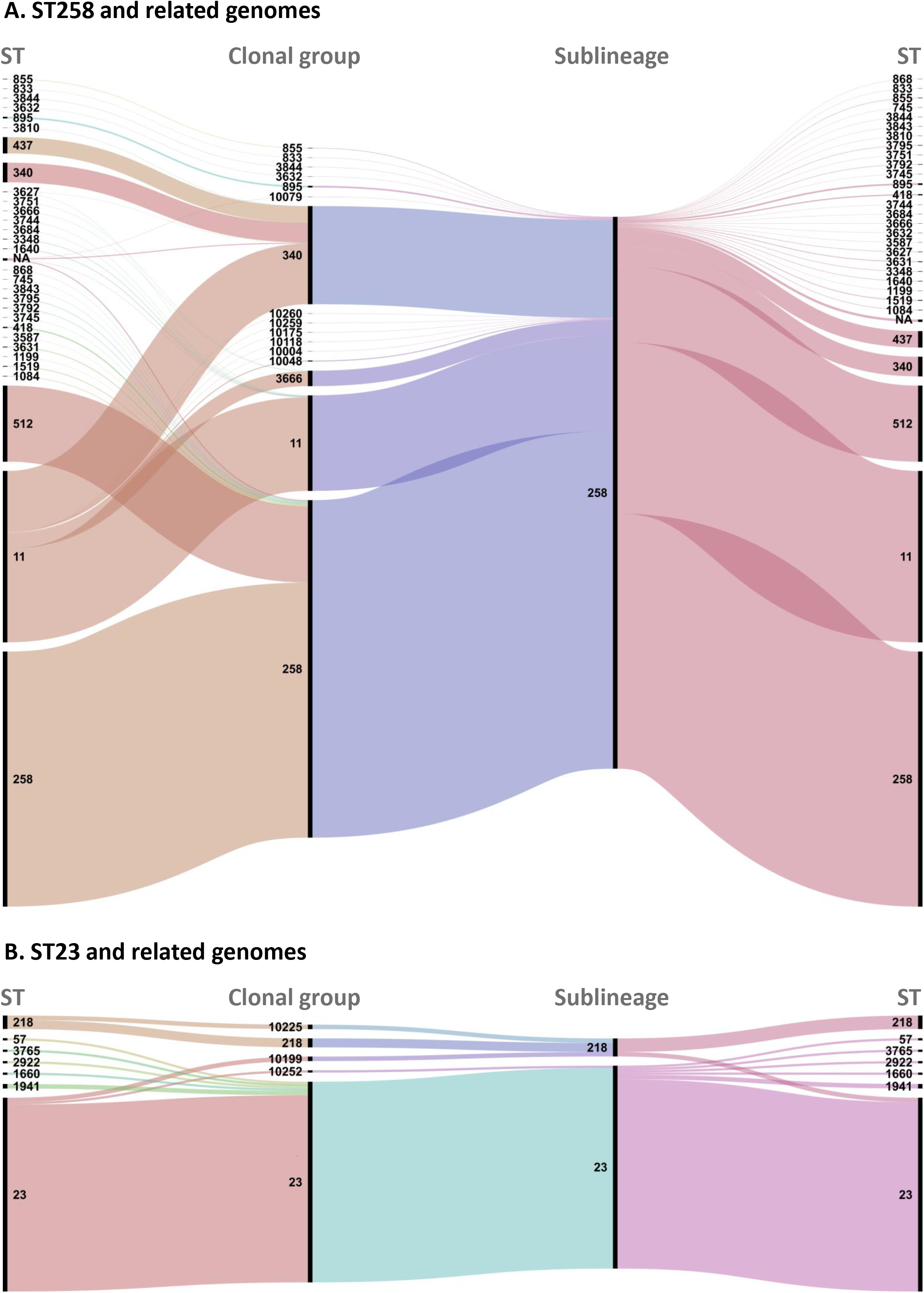
Concordance of sublineage, clonal group and 7-gene MLST classifications. Alluvial diagram obtained using RAWGraphs (Mauri *et al*. 2017: doi.org/10.1145/3125571.3125585) showing the correspondence between sequence types (ST; 7 genes identity), clonal groups (CG; 43 allelic mismatches threshold) and sublineages (SL; 190 allelic mismatches threshold). Colors are arbitrarily attributed by the software for readability.

The distribution of pairwise allelic mismatch values that involved hybrid genomes showed an additional peak around 39 shared alleles (*i.e*., 590 allelic mismatches; **Figure S7**). Therefore, these inter-phylogroup hybrids were placed into distinct partitions at the 585-mismatch level. However, as some of these hybrid genomes diverged by fewer than 585 allelic mismatches from two distinct phylogroups at the same time, they would cause fusion of phylogroup partitions upon single-linkage clustering. To highlight the impact of this phenomenon, hybrids were first filtered out, and were incorporated in a second single-linkage clustering step (supplementary appendix**; Figure S8**).

### Phylogenetic compatibility of sublineages and clonal groups

To estimate the congruence of classification groups with phylogenetic relationships among genomes, we quantified the proportion of monophyletic (single ancestor, exclusive group), paraphyletic (single ancestor, non-exclusive group) and polyphyletic (two or more distinct ancestors) groups. Regarding 7-gene MLST, 6985 (98.9%) genomes had a defined ST, *i.e*., an allele was called for the seven genes. Of the 992 distinct STs, 396 were non-singleton STs (*i.e*., comprised at least two isolates). Of these, 286 (72.2%) were monophyletic, nine were paraphyletic (2.3%) and 101 (25.5%) were polyphyletic. The monophyletic STs comprised only 22% of all genomes in non-singleton STs.

Regarding cgMLST-based classification, there were five and seven partitions at 610 and 585 allelic mismatch levels, respectively, and 100% of these were monophyletic. Among the 705 distinct sublineages, 317 (45.0%) were non-singleton, and most (310; 97.8%) of these were monophyletic (**Figure 2**); only three (0.9%) were paraphyletic, and four (1.3%) were polyphyletic (**Table S4**). The monophyletic sublineages comprised a large majority (5,961/6,672; 89.3%) of genomes in non-singleton STs.

Finally, 396 out of 1,147 (34.5%) CGs were non-singleton; most (362; 91.4%) were monophyletic (**Figure 2**), whereas eight (2.0%) were paraphyletic, and 26 (6.6%) were polyphyletic (**Table S5**). Monophyletic CGs comprised nearly half (3,030; 48.0%) of the genomes in the non-singleton CGs, whereas 3,224 (51.1%) were in polyphyletic groups, mostly in CG258, CG340 and CG15.

### Definition of shallow-level classification thresholds for Klebsiella epidemiology

Although the scgMLSTv2 scheme comprises only 629 loci, or ~10% of a typical *K. pneumoniae* genome length (512,856 nt out of 5,248,520 in the NTUH-K2044 genome), shallow-level classifications of genomic sequences might be useful for tentative outbreak delineation and epidemiological surveillance purposes. To provide flexible case cluster definitions, we classified Kp genomes using thresholds of 0, 1, 2, 4, 7 and 10 scgMLSTv2 allelic mismatches. The classification groups of identical cgMLST profiles (*i.e*., corresponding to the 0-mismatch threshold) are defined as cgST, analogous to the 7-gene ST. We observed that profiles of isolates involved in previously reported Kp outbreaks generally differed by no or 1 mismatch, with a maximum of five allelic mismatches (**Table S6**).

### Inheritance of the 7-gene ST identifiers into the cgMLST classification, and characteristics of main sublineages and clonal groups

To attribute SL and CG identifiers that corresponded maximally to the widely adopted 7-gene ST identifiers, we developed an inheritance algorithm to map MLST identifiers onto SL and CG partitions (see Methods). Of the 705 SLs, most (683; 96.9%) were named by inheritance and this was the case for 879 (76.6%) of the 1,047 CGs (**Table S4**). The resulting correspondence of cgMLST partitions with classical MLST was evident for the major groups (**Figure 4; Figure S6**). For instance, the multidrug resistant SL258 comprised isolates belonging to MLST sequence types ST258, ST11, ST512, ST340, ST437 and 25 others STs. SL258 consisted of 16 distinct clonal groups, of which the four most frequent were defined as CG258 (61.2%), CG340 (17.8%), CG11 (17.3%) and CG3666 (2.8%) (**Figure 4**). When compared to 7-gene MLST, most isolates of CG258 were ST258 (75.6%) or ST512 (22.6%), whereas CG11 mostly comprised ST11 genomes (98.0%). In turn, CG340 includes a large majority of ST11 genomes (61.8%) and only 20.0% ST340 genomes, and was named CG340 rather than CG11 because CG11 was already attributed. Likewise, the majority (83/86; 96.5%) of ST23 genomes, which are associated with pyogenic liver abscess (Lam et al., 2018), were classified into SL23, which itself consisted mainly (84/90, 93.3%) of ST23 genomes (**Figure 4**). The well-recognized emerging multidrug resistant Kp populations of ST147, ST307 and ST101 each corresponded largely to a single SL and CG (**Figure S6**).

The frequency of detection of virulence and resistance genes differed among the main sublineages and clonal groups (**Figure 5; Figure S9**). As expected (Lam et al., 2021), SL23 (median virulence score of 5) and SL86 (median score 3) were prominent ‘hypervirulent’ sublineages, and they were largely lacking resistance genes. In contrast, a majority of strains from sublineages 258, 147, 307 and 37 and of a large number of other SLs had a resistance score of 2 or more, indicative of BLSE/carbapenemases, but these groups had modest virulence scores. SL231 genomes stood out as combining high virulence and resistance scores. In some cases, clonal groups within single major sublineages had contrasted virulence and resistance gene contents (**Figure 5**).

**Figure 5.**
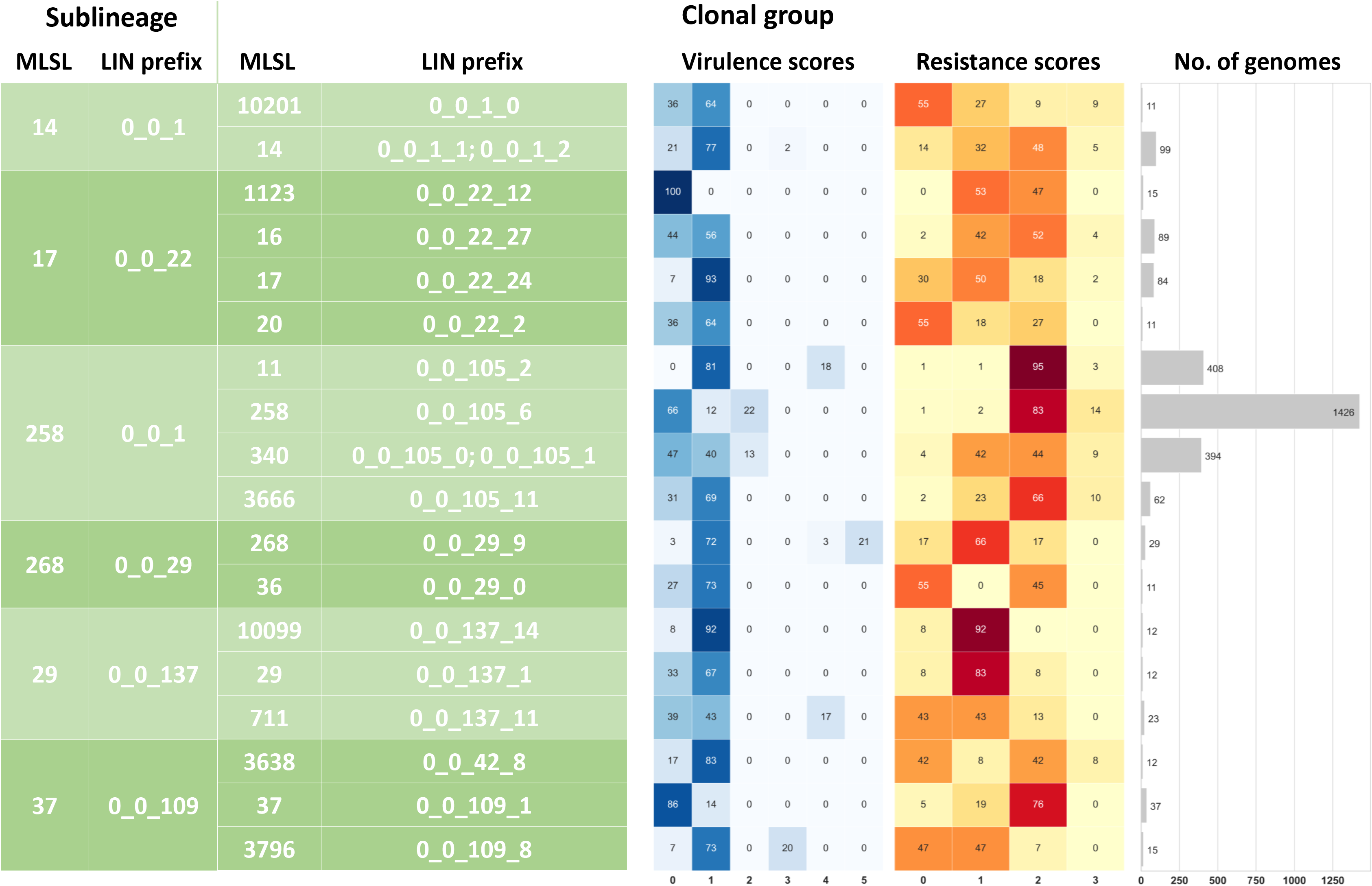
cgLIN code prefixes, virulence and resistance of some sublineages and their clonal groups. Left (green) panel: LIN prefixes of selected sublineages (SL) and clonal groups (CG). Right: heatmaps of virulence and resistance scores, and the number of genomes in each group.

### Development and implementation of a cgMLST-based LIN code system

Following the principle of the LIN code system, initially proposed based on the ANI similarity (Marakeby et al., 2014), we defined a cgMLST-based LIN (cgLIN) code approach. As LIN coding is performed sequentially, we first explored the impact on the resulting partitioning of genomes, of the order in which genomes are assigned. We confirmed that the number of partitions (hence their content too) varied according to input order (**Figure S10**). However, we determined that the order of genomes defined by a Minimum Spanning tree (MStree) (Prim, 1957) naturally induces a LIN encoding order that is optimal, *i.e*., most parsimonious with respect to the number of identifiers generated at each position of the code (see supplementary appendix). Using this MStree traversal strategy, we defined cgLIN codes for the 7,060 non-hybrid genomes (as a first step), resulting in 4,889 distinct cgLIN codes.

LIN codes can be displayed in the form of a prefix tree (**Figure 6**), which largely reflects the phylogenetic relationships among genomes. In addition, cgLIN code prefixes can be used to label particular phylogenetic lineages. For example, a single cgLIN code prefix defined each phylogroup (*e.g*., Kp1: prefix 0_0; Kp2: prefix 2_0; **Figure 6**). Likewise, a full one-to-one correspondence between prefixes and SLs was observed, and almost all (99.4%) CGs also had a unique prefix (**Table S4; Table S5; Figure 5)**.

**Figure 6.**
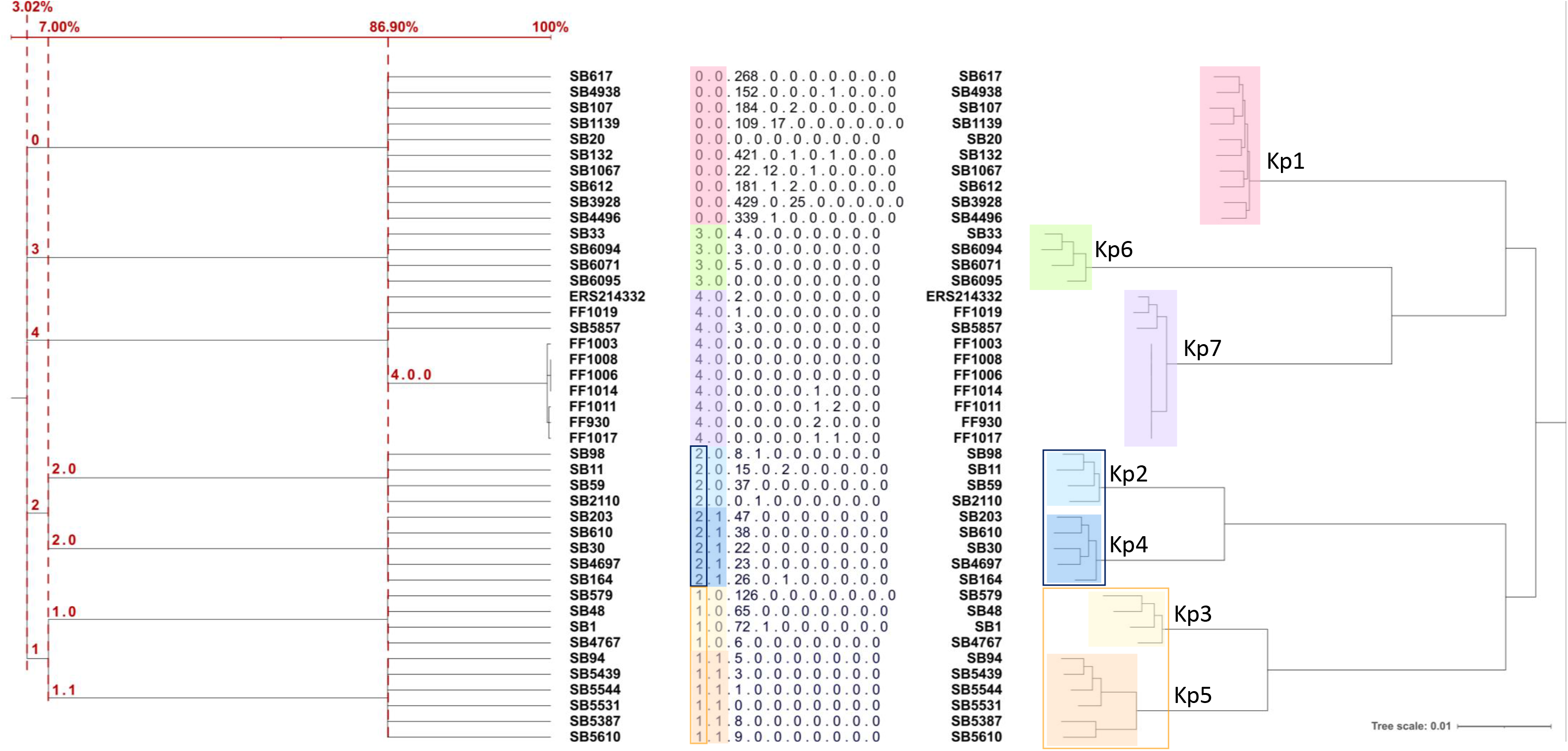
Phylogenetic relationships are reflected in cgLIN code prefixes. Left: the prefix tree generated from cgLIN codes; Right: phylogenetic relationships derived using IQ-TREE from a cgMSA. The cgLIN codes are also shown.

### Effect of hybrid genomes incorporation on the MLSL and cgLIN codes classifications

Because inter-phylogroup hybrid genomes have smaller distances to their parental phylogroups than the inter-phylogroup distances resulting from vertical evolutionary events, their incorporation into the MLSL classification may induce fusion of previously distinct single-linkage groups. To illustrate this chaining effect, the ‘hybrid’ genomes were included into the MLSL nomenclature in a second step, and fusions of previously existing partitions we observed; for example, at the 610 allelic mismatch threshold, group 2 (Kp2 and Kp4) and group 4 (Kp3 and Kp5) were merged with group 1 (Kp1). At the 585-mismatch threshold, group 5 (Kp2) and group 2 (Kp4) were merged with group 1 (Kp1). Last, at the 190-mismatch threshold, only one fusion was observed, between group 184 (SL113) and group 465 (SL1518; **Figure S11**). The partitions at other thresholds were not impacted by the addition of the hybrid genomes.

In contrast, the incorporation of hybrid genomes into the cgLIN code database left the cgLIN codes of the 7,060 previous genomes entirely unaffected; there were no merging of groups, as per design of the system. In particular, the seven phylogroup prefixes corresponding to species and subspecies remained unaffected (**Figure S11**); instead, additional prefixes were created for the hybrid genomes (**Table S1; Table S2**).

### Implementation of the genomic taxonomy in a publicly-accessible database

The MLSL nomenclature was incorporated into the Institut Pasteur *K. pneumoniae* MLST and whole genome MLST database (https://bigsdb.pasteur.fr/klebsiella) under the classification schemes functionality developed in BIGSdb version 1.21.0. In brief, the cgMLST profile of all isolates with fewer than 30 missing scgMLSTv2 alleles were assigned to a core genome sequence type (cgST), and these were grouped into single-linkage partitions for each of the 10 classification levels. For SLs and CGs, a custom classification group field (named SL or CG within the system) was additionally populated with identifiers inherited from 7-gene MLST. All cgMLST profiles and classification identifiers are freely available to use, open access.

To allow users to identify users to identify *K. pneumoniae* isolates easily, a profile matching functionality was developed, allowing to search for cgMLST profiles related to a query genome sequence. This was implemented and is available from the website sequence query page (https://bigsdb.readthedocs.io/en/latest/administration.html#scheme-profile-clustering-setting-up-classification-schemes). This functionality returns the classification identifiers (including MLST-inherited CG and SL identifiers) of the cgMLST profile that is most closely related to the query genomic sequence, along with its number of mismatches compared to the closest profile.

## Discussion

The existence within microbial species of sublineages with unique genotypic and phenotypic properties underlines the need for infra-specific nomenclatures (Lan and Reeves, 2001; Maiden et al., 1998; Rambaut et al., 2020). As for species and higher Linnaean taxonomic ranks, a strain taxonomy should: (i) recognize genetic discontinuities and capture the most relevant sublineages at different phylogenetic depths; (ii) provide an unambiguous naming system for sublineages; and (iii) provide identification methods for placement within the taxonomic framework. Here we developed a strain taxonomy consisting of a dual naming system that is grounded in population genetics and linked to an identification tool. The proposed system strains thus complies with the three fundamental pillars of taxonomy.

Although 7-gene MLST has been widely adopted as a taxonomic system of *Kp* strains, several limitations are apparent: besides its restricted resolution, MLST does not convey phylogenetic information; a single nucleotide substitution generates a different ST with unapparent relationships with its ancestor. Further, approximately half of the ST partitions were not monophyletic. The cgMLST approach extends the MLST concept to genome scale, providing much higher resolution and phylogenetic precision (Maiden et al., 2013; Zhou et al., 2021). Although other metrics such as whole-genome SNPs or ANI can be used to classify strains (Marakeby et al., 2014; Dallman et al., 2018), cgMLST presents advantages inherited from classical MLST, including standardization, reproducibility, portability and the conversion of sequences into human-readable allelic numbers. The high reproducibility and ease of interpretation of cgMLST are two critical characteristics for its adoption in epidemiological surveillance. We showed that cgMLST, based here on 629 genes, has a much broader dynamic range than ANI when considering intra-specific variation, and enables defining several hierarchical classification levels (Zhou et al., 2021). To define shallower genetic structure within sublineages resulting from recent clonal expansions, higher resolution could be achieved based *e.g*., on core gene sets of specific phylogroups or sublineages.

Optimization of threshold definitions based on population structure aims at optimizing cluster stability (Barker et al., 2018; Zhou et al., 2021). The density distribution of pairwise allelic mismatch dissimilarities within *K. pneumoniae* and related species exhibited genetic discontinuities at several phylogenetic depths. We took the benefit of this multimodal distribution to define optimal intra-specific classification thresholds, and combined a clustering consistency coefficient (Silhouette) with a newly developed strategy that evaluates cluster stability by subsampling the entire dataset. We defined four classifications at phylogenetic depths that reflected natural discontinuities within the population structure of *K. pneumoniae*, including the deep subdivisions of *K. pneumoniae sensu lato*, now recognized as seven species and/or subspecies (Rodrigues et al., 2019). The 190-mismatch sublineage level was designed to capture the numerous deep phylogenetic branches within phylogroups (Bialek-Davenet et al., 2014; Holt et al., 2015). In turn, the clonal group (CG) level was useful to capture the genetic structuration observed within these primary sublineages. For example, the CG-level nomenclature captures the evolutionary split of CG258 and CG11 from their SL258 ancestor, caused by a 1.1 Mb recombination event (Chen et al., 2014) (**Figure S5**).

Seven-locus MLST is a widely adopted nomenclature system, as illustrated by the widespread use of ST identifiers associated with hypervirulent or multidrug resistant sublineages (*e.g*., ‘*Klebsiella pneumoniae* ST258’: 293 PubMed hits; ST23: 117 hits; on July 20^th^, 2021). Backward nomenclatural compatibility is therefore critical. After applying our inheritance algorithm, most sublineages and clonal groups were labeled according to the 7-gene MLST identifier of the majority of their isolates. Widely adopted ST identifiers will therefore designate nearly the same strain groups within the proposed genomic taxonomy of *K. pneumoniae*, which should greatly facilitate its adoption. We note that the 7-gene MLST nomenclature will still have to be expanded, as this classical approach continues to be widely used. However, for practical reasons, upcoming MLST and cgMLST nomenclatural identifiers will be uncoupled, and we suggest that the cgMLST based identifiers of sublineages and clonal groups should be adopted as the reference nomenclature in the future.

Instability is a major limitation of single linkage clustering, caused by group fusion known as the chaining effect (Turner and Feil, 2007). This is particularly relevant over epidemiological timescales, where intermediate genotypes (*e.g*., a recent ancestor or recombinant) are often sampled (Feil, 2004). This issue is exacerbated in *K. pneumoniae*, where large scale recombination may result in truly intermediate genotypes (Chen et al., 2014; Holt et al., 2015), referred to as ‘hybrids’ by analogy to eukaryotic biology. Here this phenomenon was illustrated through our delayed introduction into our nomenclature, of 138 inter-phylogroup hybrid genomes. The merging of predefined classification groups can be handled by classification versioning or ad hoc rules (Zhou et al., 2021), but this is undeterministic and challenging in practice.

To address its stability issue, we complemented the single linkage clustering approach with a fully stable approach. LIN codes were proposed as a universal genome coding system (Marakeby et al., 2014; Tian et al., 2020), a key feature of which is the generation of definitive genome codes that are inherently stable. The original LIN code system was based on the ANI metric; here we noted that the ANI values that best correspond to some of the 10 cgMLST thresholds were highly similar (**Table S6**), casting doubt on the reliability of this metric for small-scale genetic distances. In addition, the ANI metric is non-reciprocal and highly dependent on comparison implementations and parameters. These shortfalls may lead to imprecision and non-reproducibility that are particularly impactful for comparisons between very similar genomes. We therefore adapted the LIN code concept to cgMLST-based similarity (*i.e*., one-complement of the allelic mismatch proportions) to classify strains into a cgLIN code system. This strategy leverages the benefits of cgMLST, and introduces more intuitive shallow-level classification thresholds. The multilevel similarity information embedded in MLSL and cgLIN ‘barcodes’ provides a human-readable snapshot of strain relationships, as nearly identical genomes have identical barcodes up to a position near the right end. In contrast to single linkage clustering partitions, one important limitation of LIN codes is that preexisting classification identifiers (e.g., ST258) cannot be mapped onto individual LIN code identifiers, because these are attributed with reference to the upper levels and are set to 0 for each downstream level. However, cgLIN code prefixes may represent useful labels for particular lineages.

A unified nomenclature of *K. pneumoniae* genotypes is required to enable communication around pathogens in the ‘One Health’ and ‘Global Health’ perspectives. This pathogen represents a rapidly growing public health threat, and the availability of a common language to designate its emerging sublineages is therefore highly timely. The proposed unified taxonomy of *K. pneumoniae* strains will facilitate advances on the biology of its sublineages across niches, time and space, and will endow surveillance networks with the capacity to efficiently monitor and control the emergence of sublineages of high public health relevance.

## Conclusions

On the bases of our analyses, we propose a dual barcoding approach to bacterial strain taxonomy, which combines the complementary advantages of stability provided by the cgLIN codes, with an unstable, but human readable multilevel single linkage nomenclature rooted in the popular 7-gene MLST nomenclature. Because they are definitive, cgLIN codes can be used for the traceability of cluster fusions that will occur occasionally in the MLSL arm of the dual taxonomy (**Figure S11**). Identification of users’ query genomes is made possible either through the BIGSdb platform that underlies the cgMLST website, or externally after export of the cgMLST profiles, both of which are publicly accessible. Further work will be needed to implement the cgLIN code approach within the BIGSdb platform or elsewhere (Tian et al., 2020) and to develop dual strain taxonomy systems for other bacterial species.

## Material & Methods

### Definition of an updated strict core genome MLST (scgMLST) genotyping scheme

We previously defined a strict core genome MLST (scgMLSTv1) scheme of 634 highly syntenic genes (Bialek-Davenet et al., 2014). Here, we updated the scgMLST scheme based on a reassessment of scgMLSTv1 loci, with the following improvements. First, two loci (KP1_2104 and *aceB*=KP1_0253) were removed because they were absent or truncated in multiple strains, based on 751 high-quality assemblies available in the BIGSdb-Pasteur *Klebsiella* database on October 16th, 2017 (project id 11 at https://bigsdb.web.pasteur.fr/cgi-bin/bigsdb/bigsdb.pl?db=pubmlst_klebsiella_isolates). Second, the remaining 632 loci templates were modified so that they would include the start and stop codons of the corresponding coding sequence (CDS); This was not the case for all CDSs in scgMLSTv1, as some loci corresponded to internal portions of CDSs. These template redefinitions were done to harmonize locus definitions across the scheme and because defining loci as complete CDSs facilitates genotyping, by allowing precise identification of the extremities of novel alleles, through the search of the corresponding start and stop codons. As a result of these locus template extensions, the 629 scgMLSTv2 genes have a summed length of 512,856 nt (9.8% of the genome of reference strain NTUH-K2044), as compared to 507,512 nt (9.7%) for the corresponding loci in scgMLSTv1.

### Genomic sequence dataset of 7060 isolates and extraction cgMLST profiles

The *K. pneumoniae* species complex (KpSC) comprises seven phylogroups that have been given taxonomic status in the prokaryotic nomenclature: *K. pneumoniae* subsp. *pneumoniae* (Kp1), *K. quasipneumoniae* subsp. *quasipneumoniae* (Kp2), *K. variicola* subsp. *variicola* (Kp3), *K. quasipneumoniae* subsp. *similipneumoniae* (Kp4), *K. variicola* subsp. *tropica* (Kp5), *K. quasivariicola* (Kp6) and *K. africana* (Kp7) (Rodrigues et al., 2019). We retrieved all KpSC genomes from the NCBI assembly database (GenBank) on March 15, 2019, corresponding to 8,125 assemblies. We then chose high-quality assemblies by excluding assemblies: (i) which contained more than 1,000 contigs of size >200 nt; (ii) for which the average nucleotide identity (ANI) values (estimated using FastANI v1.1) were < 96% against every reference strain of the taxonomic diversity of the KpSC (Rodrigues et al., 2019; **Table S8**); (iii) of size ≤ 4.5Mb or ≥ 6.5 Mb; and (iv) with G+C% content >59%. The three last criteria excluded possible contamination or non-KpSC genomes.

The resultant 7,388 ‘high-quality’ draft genomes (**Table S1**) were scanned for scgMLSTv2 alleles, using the BLASTN algorithm implemented in the BIGSdb platform (Jolley et al., 2018; Jolley and Maiden, 2010), with 90% identity, 90% length coverage, word size 30, with type alleles only (as defined below). After this step, 235 profiles (including the ‘*K. quasivariicola’* reference strain KPN1705, SB6096) were excluded because they had more than 30 missing alleles.

The resulting dataset comprised 36 taxonomic references of Kp1-Kp7 (Rodrigues et al., 2019) that, together with eight additional genomes of phylogroup Kp7, were considered as a reference dataset of the KpSC taxa. Besides these 44 reference genomes, 7,154 GenBank genomes were retained, resulting in a total dataset of 7,198 genomes (**Table S1; Table S2**). For some analyses, 138 genomes were set aside, defined as ‘hybrids’ between phylogroups (see below), resulting in a 7,060-genome dataset (**Figure S2**).

### Recording sequence variation at the cgMLST gene loci

Allelic variation at scgMLSTv2 loci was determined with the following strategy. First the sequence of strain NTUH-K2044 was used as the reference genome, with all its alleles defined as allele 1. Then, BLASTN searches (70% identity, 90% length coverage) were carried out using allele 1 as query against the genomic sequences of reference genomes 18A069, 342, 01A065, 07A044, CDC4241-71 and 08A119, representing major lineages (phylogroups Kp2 to Kp6, including two genomes of Kp3 and excluding Kp7, which was not discovered yet) of the KpSC (Blin et al., 2017). Only sequences with a complete CDS (start and stop, no internal frameshift) and within a plus/minus 5% range of the reference size were accepted. Alleles defined from these reference genomes and from NTUH-K2044 were then defined as type alleles.

New alleles in the database were identified by BLASTN searches using a 90% identity threshold, 90% length coverage and a word size of 30 and the above defined type alleles. The use of type alleles avoided expanding the sequence space of alleles in an uncontrolled way, at the cost of losing a few highly divergent alleles, which may have replaced original (vertically inherited) alleles by horizontal gene transfer (HGT) and homologous recombination. As for type alleles, novel alleles were accepted only if they (i) corresponded to a complete CDS (start and stop codons with no internal frameshift mutations) and (ii) were within a 5% (plus/minus) of the size of the type allele size. Novel allele sequences were also excluded if they came from assemblies with more than 500 contigs of size > 200 nt, as these may correspond to low quality assemblies and that might contain artifactual alleles. Genome assemblies based on 454 sequencing technology, which are prone to frameshifts, were also excluded for novel allele definitions. No genome assemblies based on IonTorrent sequencing technology were found.

In order to speed the scanning process, we used the fast scan option (-e -f) of the BIGSdb autotag.pl script (https://bigsdb.readthedocs.io/en/latest/offline_tools.html). This option limits the BLASTN search to a few exemplar alleles, which are used as query to find the genomic region corresponding to the locus. In a second step, a direct database lookup of the region was performed to identify the exact allele.

### Definition of MLST sequence types (ST) and core genome MLST sequence types (cgST)

Classical 7-gene MLST loci have been defined previously (Diancourt et al., 2005) as internal portions of the seven protein-coding genes *gapA*, *infB*, *mdh*, *pgi*, *phoE*, *rpoB* and *tonB*. Novel alleles were defined in the Institut Pasteur *Klebsiella* MLST and whole-genome MLST database https://bigdb.pasteur.fr/klebsiella. In 7-locus MLST, the combination of the seven allelic numbers determines the isolate profile, and each unique profile is attributed a sequence type (ST) number. Incomplete MLST profiles with one (or more) missing gene(s) are recorded in the isolates database but in these cases, no ST number can be attributed and the profiles are therefore not defined in the sequence definition database. The 7-locus MLST genes were not included in the scgMLSTv2 scheme.

Similar to ST identifiers used for unique 7-gene MLST allelic combinations, each distinct cgMLST profile can be assigned a unique identifier; however, when using draft genomes, cgMLST data can be partly incomplete due to *de novo* assembly shortcomings or missing loci. cgSTs were therefore defined for cgMLST profiles with no more than 30 uncalled alleles out of the 629 cgMLST loci. cgSTs were stored in the Institut Pasteur *Klebsiella* database.

### Phylogenetic analyses, recombination tests and screens for virulence and resistance genes

JolyTree v2.0 (Criscuolo, 2019, 2020) was used to reconstruct a phylogeny of the *K. pneumoniae* species complex. For this, first a single linkage (SL) was performed to cluster cgSTs into partitions. SL clustering was applied on the pairwise distances between allelic profiles, defined as the number of loci with different alleles, normalized by the number of loci with alleles called in both profiles. A threshold of 8 mismatches was defined, resulting in 2,417 clusters. One genome from each of these 2,417 clusters was used as an exemplar for phylogenetic analysis.

A core genome multiple sequence alignment (cg-MSA) of 7,060 cgMLST profiles free of evidence for inter-phylogroup ‘hybridization’ (see below) was constructed. The gene sequences were retrieved based on allele number in the sequence definition database, individual gene sequences were aligned (missing alleles were converted into gaps), and the alignments were concatenated into a cg-MSA. IQ-TREE v2.0.6 was used to infer a phylogenetic tree with the GTR+G model.

Locus-by-locus recombination analyses were computed with the PHI test (Bruen et al., 2006) using PhiPack v1.0. Kleborate v2.0.4 (Lam et al., 2021) was employed to identify known resistance genes in genomic sequences, based on CARD v3.0.8 database, with identity >80% and coverage >90%, with the --resistance option.

### Detection of hybrids

Horizontal gene transfer of large portions of the genome can occur among isolates belonging to distinct KpSC phylogroups (Holt et al., 2015). Additionally, MLST or scgMLST alleles may have been transferred horizontally from non-KpSC members, for example *E. coli*. For the purpose of phylogeny-based classification, putative hybrid genomes were excluded. To define genomes that result from large inter-phylogroup recombination events, the gene-by-gene approach was used to define an original strategy, outlined briefly here and more thoroughly in the supplementary appendix (Detection of hybrids): for each locus, each allele was unambiguously labelled by one of the seven KpSC phylogroup of origin, if possible; next, for each profile, a phylogroup homogeneity index (*i.e*., proportion of alleles labelled by the predominant phylogroup) was derived. The distributions of the phylogroup homogeneity indices allowed determining hybrid genomes (**Figure S12**). Exclusion of such hybrid genomes resulted in a genomic dataset of 7,060 isolates deemed as having a majority of alleles inherited from within a single phylogroup. Of the 44 reference genomes, one (SB1124, of phylogroup Kp2) was defined as having a hybrid origin: 414 alleles were attributed to Kp2, whereas 150 alleles originated from non-KpSC species; as a result, 73.4% of SB1124 alleles were part of the majority phylogroup, which was below the defined threshold of 78%. The quantification of recombination breakpoints was performed based on the position of cgMLST loci on the NTUH-K2044 reference genome (NC_012731), counting the number of recombination breakpoints in each successive 500 kb fragment along the reference genome.

### Identification of genetic discontinuities in the KpSc population structure

The tool MSTclust v0.21b (https://gitlab.pasteur.fr/GIPhy/MSTclust) was developed to ease the single-linkage clustering of cgMLST profiles from their pairwise allelic mismatch dissimilarities, as well as to assess the efficiency of the resulting profile partitioning (for details, see supplementary appendix: Minimum Spanning tree-based clustering of cgMLST profiles). Briefly, for each threshold *t* (= 0 to 629 allelic mismatches), the clustering consistency was assessed using the silhouette metrics *S_t_* (Rousseeuw, 1987), whereas its robustness to profile subsampling biases was assessed using a dedicated metrics *W_t_* based on the adjusted Wallace coefficients (Wallace, 1983; Severiano et al., 2011). Both consistency and stability coefficients *S_t_* and *W_t_* converge to 1 when the threshold *t* leads to a clustering that is consistent with the ‘natural’ grouping and is robust to subsampling biases, respectively.

The adjusted Rand index *R_t_* (Carrico et al., 2006; Hubert and Arabie, 1985) was used to assess the global concordance between SLC partitions and those induced by classifications into 7-gene MLST sequence types, subspecies and species.

### Diversity and phylogenetic compatibility indices

Simpson’s diversity index was computed using the www.comparingpartitions.info website (Carrico et al., 2006). The clade compatibility index of STs or other groups was calculated using the ETE Python library (http://etetoolkit.org/docs/latest/tutorial/tutorial_trees.html#checking-the-monophyly-of-attributes-within-a-tree), in order to define whether their constitutive genomes formed a monophyletic, paraphyletic or polyphyletic group within the recombination-purged sequence-based phylogeny of the core genome. We estimated clade compatibility as the proportion of non-singleton STs, sublineages or clonal groups that were monophyletic.

### Classification of cgMLST profiles into clonal groups and sublineages

The classification scheme functionality was implemented within BIGSdb v1.14.0 and relies on single linkage clustering. Briefly, cgSTs were defined in the sequence definitions (‘seqdef’) database as distinct profiles with fewer than 30 missing alleles over the scgMLST scheme, and their pairwise cgMLST distance was computed as the number of distinct alleles. To account for missing data, a relative threshold was used for clustering: the number of allelic mismatches was multiplied by the proportion of loci for which an allele was called in both strains. Hence, in order to be grouped, the number of matching alleles must exceed: (the number of loci called in both strains x (total loci - defined threshold)) / total loci. cgSTs and their corresponding sublineage (SL), clonal group (CG) and other levels partition identifiers, are stored in the seqdef database and are publicly available. Here, classification schemes were defined in the *Klebsiella* seqdef database on top of the scgMLSTv2 scheme, and host single linkage clustering group identifiers at the 10 defined cgMLST allelic mismatch thresholds (see Results). For classification groups defined using 43 and 190 allelic mismatch thresholds, scheme fields were defined and populated with the identifiers defined by inheritance from 7-gene MLST ST identifiers (see below).

### Nomenclature inheritance algorithm

In order to attribute to each sublineage (SL) and clonal group (CG), an identifier that would maximally reflect the widely adopted 7-gene ST identifier of the corresponding isolates, a set of naming rules were developed that prioritized the most abundant ST observed among isolates of each group. The formalization of the algorithm is given in supplementary appendix Nomenclature inheritance algorithm, and its implementation as a Python script is provided at https://gitlab.pasteur.fr/BEBP/inheritance-algorithm. Here the process for the CG level is described, but the algorithm was also applied to the SL level. Briefly, the data (*e.g*., a list of CG-ST pairs) can be formalized as a bipartite graph, in which each CG and ST are nodes, and each non-empty CG-ST intersection is an edge. The weight of each edge is equal to the number of isolates sharing the corresponding CG and ST identifiers. Based on this representation, the algorithm will consist of following all edges in the input graph, in the order of decreasing weight. The approach prioritizes the most frequent ST/CG pairs of isolates, *i.e*., those that are predominant in the dataset and thus naturally transfers to the CG nomenclature, the identifiers of the highest frequency STs. Rules were implemented to treat the cases of equality of representation of two or more STs connected to the same CG. Once all edges were removed from the graph, it may be that some CGs were not named, for example, because the identifier of their unique corresponding ST was already attributed to another CG. For these orphan CGs, iteratively, the attributed identifier corresponds to the maximal CG identifier already attributed, plus one.

### Adaptation of the LIN code approach to cgMLST: defining cgLIN codes

Vinatzer and colleagues proposed an original nomenclature method in which each genome is attributed a Life Identification Number code (LIN code), based on genetic similarity with the closest previously encoded member of the nomenclature (Marakeby et al., 2014; Weisberg et al., 2015; Vinatzer et al., 2016, 2017; Tian et al., 2020). In this proposal, the similarity between genomes was based on ANI (Average Nucleotide Identity; Konstantinidis and Tiedje, 2005; Goris et al., 2007), with a set of 24 thresholds corresponding to ANI percentages of 60, 70, 75, 80, 85, 90, 95, 98, 98.5, 99, 99.25, 99.5, 99.75, 99.9, 99.925, 99.95, 99.975, 99.99, 99.999 and 99.9999. Here, the method was adapted by replacing the ANI metric by the similarity between cgMLST profiles, defined as the proportion of loci with identical alleles normalized by the number of loci with alleles called in both profiles. These codes, which we refer to as cgLIN codes, are composed of a set of *p*-positions, each corresponding to a threshold of similarity between genomes. These similarity thresholds are sorted in ascending order (*i.e*., *s*_*p*_ < *s*_*p*+1_), the first positions of the code (on the left side) thus corresponding to low levels of similarity. Following the initial proposal, the codes are assigned as follows (**Figure S13**): (step 1) The code is initialized with the first strain being assigned the value “0” at all positions; (step 2) The encoding rule for a new genome *i* is based on the closest genome *j* already encoded as follows, from the similarity *s*_*ij*_ ∈ ]*s*_*p*−1_; *s*_*p*_]:

i. identical to code *j* up to and including position *p* − 1.
ii. for the position *p*: maximum value observed at this position (among the subset of codes sharing the same prefix at the position *p* − 1) incremented by 1.
iii. “0” to all downstream positions, from *p* + 1 included.

For each genome to be encoded, step 2 is repeated.

A set of 10 cgMLST thresholds were defined as follows: first, four thresholds were chosen above the similarity values peak observed between *Klebsiella* species (*s*_*p*_ = (629 − 610)/629 = 0.03), subspecies (*s*_*p*_ = (629 − 585)/629 = 0.07), main sublineages (*s*_*p*_ = (629 − 190)/629 = 0.70) and clonal groups (*s*_*p*_ = (629 − 43)/629 = 0.93). Second, we included six thresholds deemed useful for epidemical studies, corresponding to 10, 7, 4, 2, 1 and 0 cgMLST mismatches.

This encoding system conveys phylogenetic information, as two genomes with identical prefixes in their respective cgLIN codes can be understood as being similar, to an extent determined by the length of their common prefix. Isolates having cgMLST profiles with 100% identity (no mismatch at loci called in both genomes) will have exactly the same cgLIN code. For example, LIN codes 0_0_22_12_0_1_0_0_0_0 and 4_0_3_0_0_0_0_0_0_0 would denote two strains belonging to distinct species (as they differ by their first number in the code). LIN codes 0_0_105_6_0_0_75_1_1_0 and 0_0_105_6_0_0_75_1_0_0 correspond to strains from Kp1 (prefix 0_0) that differ by only 2 loci; they are identical up to the second bin, corresponding to 2 locus mismatches (**Figure S13**; note that 0 and 1 mismatches are both included in the last bin: genomes have an identical identifier when having 0 difference, and a different identifier when having 1 mismatch (**Figure S14**).

The impact of genome input order on the number of cgLIN code partitions at a given threshold was defined using the 7060 high-quality, non-hybrid cgMLST profiles, which were encoded 500 times with random input orders (see details in the supplementary appendix: Impact of strains input order on LIN codes, and use of Prim’s algorithm).

The scripts for cgLIN code database creation were made available via GitLab BEBP (https://gitlab.pasteur.fr/BEBP/LINcoding).

## Acknowledgements

This work builds partly on the classical 7-gene MLST database, which has been curated by Virginie Passet, Radek Izdebski, Carla Rodrigues and Federica Palma. We thank Adrien Le Meur for help with the genome dataset constitution and initial analyses of intra-outbreak variation.

## Funding

MH was supported financially by the PhD grant “Codes4strains” from the European Joint Programme One Health, which has received funding from the European Union’s Horizon 2020 Research and Innovation Programme under Grant Agreement No. 773830. This work was supported financially by the French Government’s Investissement d’Avenir program Laboratoire d’Excellence “Integrative Biology of Emerging Infectious Diseases” (ANR-10-LABX-62-IBEID). BIGSdb development is funded by a Wellcome Trust Biomedical Resource grant (218205/Z/19/Z). This work used the computational and storage services provided by the IT department at Institut Pasteur.

## Authors license statement

This research was funded, in whole or in part, by Institut Pasteur and by European Union’s Horizon 2020 research and innovation programme. For the purpose of open access, the authors have applied a CC-BY public copyright license to any Author Manuscript version arising from this submission.

## Declaration of interest statement

The authors declare no conflict of interest.

## Ethical approval statement

Not relevant.

## Author contributions

S.B. designed and coordinated the study. M.H. performed the genomic analyses, and cgLIN code developments and implementations. J.G. designed the novel version of the cgMLST scheme. A.C. developed the MSTclust tool (and associated metrics), and supervised cgLIN developments and phylogenetic analyses. K.A.J. and M.C.J.M. designed and developed the BIGSdb platform functionalities. S.B. and M.H. wrote the initial version of the manuscript, with input from A.C. All authors provided input to the manuscript and reviewed the final version.

## Notes

### Competing Interest Statement

The authors have declared no competing interest.

## References

Achtman, M., Wain, J., Weill, F.-X., Nair, S., Zhou, Z., Sangal, V., Krauland, M.G., Hale, J.L., Harbottle, H., Uesbeck, A., et al. (2012). Multilocus Sequence Typing as a Replacement for Serotyping in Salmonella enterica. PLoS Pathog 8, e1002776.

Barker, D.O., Carriço, J.A., Kruczkiewicz, P., Palma, F., Rossi, M., and Taboada, E.N. (2018). Rapid Identification of Stable Clusters in Bacterial Populations Using the Adjusted Wallace Coefficient. BioRxiv 299347.

van Belkum, A., Tassios, P.T., Dijkshoorn, L., Haeggman, S., Cookson, B., Fry, N.K., Fussing, V., Green, J., Feil, E., Gerner-Smidt, P., et al. (2007). Guidelines for the validation and application of typing methods for use in bacterial epidemiology. Clin Microbiol Infect 13 Suppl 3, 1–46.

Bialek-Davenet, S., Criscuolo, A., Ailloud, F., Passet, V., Jones, L., Delannoy-Vieillard, A.S., Garin, B., Le Hello, S., Arlet, G., Nicolas-Chanoine, M.H., et al. (2014). Genomic definition of hypervirulent and multidrug-resistant Klebsiella pneumoniae clonal groups. Emerg Infect Dis 20, 1812–1820.

Blin, C., Passet, V., Touchon, M., Rocha, E.P.C., and Brisse, S. (2017). Metabolic diversity of the emerging pathogenic lineages of Klebsiella pneumoniae. Environ Microbiol 19, 1881–1898.

Bowers, J.R., Kitchel, B., Driebe, E.M., MacCannell, D.R., Roe, C., Lemmer, D., de Man, T., Rasheed, J.K., Engelthaler, D.M., Keim, P., et al. (2015). Genomic Analysis of the Emergence and Rapid Global Dissemination of the Clonal Group 258 Klebsiella pneumoniae Pandemic. PLoS ONE 10, e0133727.

Brisse, S., and Verhoef, J. (2001). Phylogenetic diversity of Klebsiella pneumoniae and Klebsiella oxytoca clinical isolates revealed by randomly amplified polymorphic DNA, gyrA and parC genes sequencing and automated ribotyping. Int J Syst Evol Microbiol 51, 915–924.

Brisse, S., Grimont, F., and Grimont, P.A.D. (2006). The genus Klebsiella. In The Prokaryotes A Handbook on the Biology of Bacteria, (New York: Springer), pp. 159–196.

Bruen, T.C., Philippe, H., and Bryant, D. (2006). A simple and robust statistical test for detecting the presence of recombination. Genetics 172, 2665–2681.

Carrico, J.A., Silva-Costa, C., Melo-Cristino, J., Pinto, F.R., de Lencastre, H., Almeida, J.S., and Ramirez, M. (2006). Illustration of a common framework for relating multiple typing methods by application to macrolide-resistant Streptococcus pyogenes. Journal of Clinical Microbiology 44, 2524–2532.

Chen, L., Mathema, B., Pitout, J.D.D., DeLeo, F.R., and Kreiswirth, B.N. (2014). Epidemic Klebsiella pneumoniae ST258 is a hybrid strain. MBio 5, e01355–01314.

Cowan, S.T. (1965). PRINCIPLES AND PRACTICE OF BACTERIAL TAXONOMY--A FORWARD LOOK. J. Gen. Microbiol. 39, 143–153.

Criscuolo, A. (2019). A fast alignment-free bioinformatics procedure to infer accurate distance-based phylogenetic trees from genome assemblies. Research Ideas and Outcomes 5, e36178.

Criscuolo, A. (2020). On the transformation of MinHash-based uncorrected distances into proper evolutionary distances for phylogenetic inference. F1000Res 9, 1309.

Dallman, T., Ashton, P., Schafer, U., Jironkin, A., Painset, A., Shaaban, S., Hartman, H., Myers, R., Underwood, A., Jenkins, C., et al. (2018). SnapperDB: a database solution for routine sequencing analysis of bacterial isolates. Bioinformatics 34, 3028–3029.

Diancourt, L., Passet, V., Verhoef, J., Grimont, P.A., and Brisse, S. (2005). Multilocus sequence typing of Klebsiella pneumoniae nosocomial isolates. J Clin Microbiol 43, 4178–4182.

Feil, E.J. (2004). Small change: keeping pace with microevolution. Nat. Rev. Microbiol. 2, 483–495.

Fevre, C., Passet, V., Weill, F.X., Grimont, P.A., and Brisse, S. (2005). Variants of the Klebsiella pneumoniae OKP chromosomal beta-lactamase are divided into two main groups, OKP-A and OKP-B. Antimicrobial Agents and Chemotherapy 49, 5149–5152.

Goris, J., Konstantinidis, K.T., Klappenbach, J.A., Coenye, T., Vandamme, P., and Tiedje, J.M. (2007). DNA-DNA hybridization values and their relationship to whole-genome sequence similarities. International Journal of Systematic and Evolutionary Microbiology 57, 81–91.

Hacker, J., and Kaper, J.B. (2000). Pathogenicity islands and the evolution of microbes. Annu Rev Microbiol 54, 641–679.

Holt, K.E., Wertheim, H., Zadoks, R.N., Baker, S., Whitehouse, C.A., Dance, D., Jenney, A., Connor, T.R., Hsu, L.Y., Severin, J., et al. (2015). Genomic analysis of diversity, population structure, virulence, and antimicrobial resistance in Klebsiella pneumoniae, an urgent threat to public health. Proc Natl Acad Sci U S A 112, E3574–81.

Hubert, L., and Arabie, P. (1985). Comparing partitions. Journal of Classification 2, 193–218.

Jolley, K.A., and Maiden, M.C. (2010). BIGSdb: Scalable analysis of bacterial genome variation at the population level. BMC Bioinformatics 11, 595.

Jolley, K.A., Bray, J.E., and Maiden, M.C.J. (2018). Open-access bacterial population genomics: BIGSdb software, the PubMLST.org website and their applications. Wellcome Open Res 3, 124.

Konstantinidis, K.T., and Tiedje, J.M. (2005). Genomic insights that advance the species definition for prokaryotes. Proc Natl Acad Sci U S A 102, 2567–2572.

Lam, M.M.C., Wyres, K.L., Duchêne, S., Wick, R.R., Judd, L.M., Gan, Y.-H., Hoh, C.-H., Archuleta, S., Molton, J.S., Kalimuddin, S., et al. (2018). Population genomics of hypervirulent Klebsiella pneumoniae clonal-group 23 reveals early emergence and rapid global dissemination. Nat Commun 9, 2703.

Lam, M.M.C., Wick, R.R., Watts, S.C., Cerdeira, L.T., Wyres, K.L., and Holt, K.E. (2021). A genomic surveillance framework and genotyping tool for Klebsiella pneumoniae and its related species complex. Nat Commun 12, 4188.

Lan, R., and Reeves, P.R. (2001). When does a clone deserve a name? A perspective on bacterial species based on population genetics. Trends Microbiol 9, 419–424.

Long, S.W., Linson, S.E., Ojeda Saavedra, M., Cantu, C., Davis, J.J., Brettin, T., and Olsen, R.J. (2017). Whole-Genome Sequencing of a Human Clinical Isolate of the Novel Species Klebsiella quasivariicola sp. nov. Genome Announc 5.

Maiden, M.C., Bygraves, J.A., Feil, E., Morelli, G., Russell, J.E., Urwin, R., Zhang, Q., Zhou, J., Zurth, K., Caugant, D.A., et al. (1998). Multilocus sequence typing: a portable approach to the identification of clones within populations of pathogenic microorganisms. Proc. Natl. Acad. Sci. U. S. A. 95, 3140–3145.

Maiden, M.C., van Rensburg, M.J., Bray, J.E., Earle, S.G., Ford, S.A., Jolley, K.A., and McCarthy, N.D. (2013). MLST revisited: the gene-by-gene approach to bacterial genomics. Nature Reviews. Microbiology 11, 728–736.

Marakeby, H., Badr, E., Torkey, H., Song, Y., Leman, S., Monteil, C.L., Heath, L.S., and Vinatzer, B.A. (2014). A system to automatically classify and name any individual genome-sequenced organism independently of current biological classification and nomenclature. PLoS ONE 9, e89142.

Moura, A., Criscuolo, A., Pouseele, H., Maury, M.M., Leclercq, A., Tarr, C., Björkman, J.T., Dallman, T., Reimer, A., Enouf, V., et al. (2016). Whole genome-based population biology and epidemiological surveillance of Listeria monocytogenes. Nat Microbiol 2, 16185.

Prim, R.C. (1957). Shortest Connection Networks And Some Generalizations. Bell System Technical Journal 36, 1389–1401.

Rambaut, A., Holmes, E.C., O’Toole, Á., Hill, V., McCrone, J.T., Ruis, C., du Plessis, L., and Pybus, O.G. (2020). A dynamic nomenclature proposal for SARS-CoV-2 lineages to assist genomic epidemiology. Nat Microbiol 5, 1403–1407.

Rodrigues, C., Passet, V., Rakotondrasoa, A., Diallo, T.A., Criscuolo, A., and Brisse, S. (2019). Description of Klebsiella africanensis sp. nov., Klebsiella variicola subsp. tropicalensis subsp. nov. and Klebsiella variicola subsp. variicola subsp. nov. Res. Microbiol.

Rousseeuw, P.J. (1987). Silhouettes: A graphical aid to the interpretation and validation of cluster analysis. Journal of Computational and Applied Mathematics 20, 53–65.

Selander, R.K., and Levin, B.R. (1980). Genetic diversity and structure in Escherichia coli populations. Science 210, 545–547.

Severiano, A., Pinto, F.R., Ramirez, M., and Carriço, J.A. (2011). Adjusted Wallace Coefficient as a Measure of Congruence between Typing Methods. J. Clin. Microbiol. 49, 3997–4000.

Sneath, P.H.A. (1992). International Code of Nomenclature of Bacteria (Washington, D.C.: American Society for Microbiology).

Struelens, M.J., De Gheldre, Y., and Deplano, A. (1998). Comparative and library epidemiological typing systems: outbreak investigations versus surveillance systems. Infect Control Hosp Epidemiol 19, 565–569.

Struve, C., Roe, C.C., Stegger, M., Stahlhut, S.G., Hansen, D.S., Engelthaler, D.M., Andersen, P.S., Driebe, E.M., Keim, P., and Krogfelt, K.A. (2015). Mapping the Evolution of Hypervirulent Klebsiella pneumoniae. MBio 6, e00630.

Tian, L., Huang, C., Mazloom, R., Heath, L.S., and Vinatzer, B.A. (2020). LINbase: a web server for genome-based identification of prokaryotes as members of crowdsourced taxa. Nucleic Acids Research 48, W529–W537.

Turner, K.M., and Feil, E.J. (2007). The secret life of the multilocus sequence type. International Journal of Antimicrobial Agents 29, 129–135.

Vinatzer, B.A., Weisberg, A.J., Monteil, C.L., Elmarakeby, H.A., Sheppard, S.K., and Heath, L.S. (2016). A Proposal for a Genome Similarity-Based Taxonomy for Plant-Pathogenic Bacteria that Is Sufficiently Precise to Reflect Phylogeny, Host Range, and Outbreak Affiliation Applied to Pseudomonas syringae sensu lato as a Proof of Concept. Phytopathology^®^ 107, 18–28.

Vinatzer, B.A., Tian, L., and Heath, L.S. (2017). A proposal for a portal to make earth’s microbial diversity easily accessible and searchable. Antonie van Leeuwenhoek 110, 1271–1279.

Wallace, D.L. (1983). A Method for Comparing Two Hierarchical Clusterings: Comment. Journal of the American Statistical Association 78, 569–576.

Weisberg, A.J., Elmarakeby, H.A., Heath, L.S., and Vinatzer, B.A. (2015). Similarity-Based Codes Sequentially Assigned to Ebolavirus Genomes Are Informative of Species Membership, Associated Outbreaks, and Transmission Chains. Open Forum Infectious Diseases 2.

Wyres, K.L., Gorrie, C., Edwards, D.J., Wertheim, H.F.L., Hsu, L.Y., Van Kinh, N., Zadoks, R., Baker, S., and Holt, K.E. (2015). Extensive Capsule Locus Variation and Large-Scale Genomic Recombination within the Klebsiella pneumoniae Clonal Group 258. Genome Biol Evol 7, 1267–1279.

Wyres, K.L., Lam, M.M.C., and Holt, K.E. (2020a). Population genomics of Klebsiella pneumoniae. Nat Rev Microbiol 18, 344–359.

Wyres, K.L., Nguyen, T.N.T., Lam, M.M.C., Judd, L.M., van Vinh Chau, N., Dance, D.A.B., Ip, M., Karkey, A., Ling, C.L., Miliya, T., et al. (2020b). Genomic surveillance for hypervirulence and multi-drug resistance in invasive Klebsiella pneumoniae from South and Southeast Asia. Genome Med 12, 11.

Zhou, Z., Charlesworth, J., and Achtman, M. (2021). HierCC: A multi-level clustering scheme for population assignments based on core genome MLST. Bioinformatics btab234.

