## Supplementary appendix for "A dual barcoding approach to bacterial strain nomenclature: Genomic taxonomy of *Klebsiella pneumoniae* strains"

**Contents:**

*Detection of inter-phylogroup hybrids*

*Minimum Spanning tree-based clustering of cgMLST profiles*

*Nomenclature inheritance algorithm*

*Impact of strains input order on LIN codes, and use of Prim's algorithm*

*List of supplementary figure legends*

*List of supplementary tables*

*Supplementary references*

##### *Detection of inter-phylogroup hybrids*

To define hybrid genomes (*i.e.,* arising from multiple ancestral populations), the 45 reference genomes representative of the seven phylogroups Kp1-Kp7 (**Table S8**) were used as models. First, for each of the 7,388 other genome sequences, the closest reference genome was determined by estimating the average nucleotide identity (ANI) using FastANI v1.1 (Jain et al., 2018). Every genome *x* with ANI percentage > 99% against its closest reference genome *y* was then classified without ambiguity into the same phylogroup as the one of *y*. Second, for each genome classified into a phylogroup (Kp1-Kp7), all its cgMLST alleles were labelled with this phylogroup. Third, for each of the 629 scgMLSTv2 loci, every distinct allele associated to more than one phylogroup labels was unlabeled, given that such an allele cannot be considered as a reliable representative of a unique phylogroup (*e.g.*, it was too conserved, or involved in horizontal transfer between phylogroups). Fourth, for each locus, every unlabeled sequence identical to one of the remaining labelled alleles (i.e., sequence belonging to a genome that was not assigned to a phylogroup during step 1) was labelled accordingly. Such a procedure enabled the characterization of a large set of alleles that are each representative of one of the seven phylogroups Kp1-Kp7.

As a result, almost all cgMLST profiles were mostly made up by alleles belonging to only one phylogroup label (see **Figure S15**). However, notable exceptions were observed, with some cgMLST profiles being composed of alleles belonging to two phylogroup labels (see e.g., **Figure S15**). To define putative hybrid profiles, a phylogroup homogeneity index was determined for each profile, defined as the proportion of loci labelled with the predominant phylogroup (normalized by the number of non-missing alleles called in the profile). As expected, for each phylogroup Kp1-Kp7, most profiles are associated with high phylogroup homogeneity indices (see distributions in **Figure S12**). However, a total of 138 cgMLST profiles (1.9%; mainly within Kp1, Kp2 and Kp4) were associated with atypically smaller homogeneity indices, and mapping of allele phylogroup labels along the chromosome showed that many of these 138 cgMLST profiles appeared to result from large-scale inter-phylogroup recombination, while a few were made up by many unlabeled alleles (**Figure S1**).

Large recombination events were detected in 1.9% (138) genomes and mainly involved phylogroups Kp1, Kp2 and Kp4; we found 42 Kp4 genomes resulting from large-scale recombination of Kp1 out of the 50 hybrid Kp4 genomes (**Figure S1**), while others a multitude of small-scale recombination events. Next, 17 Kp1 genomes were observed with a large-scale recombination (12 with a Kp2 insertion, 4 with a Kp4 insertion and 1 with a Kp3 insertion), as well as 3 Kp2 genomes resulting from a large Kp1 insertion. In addition, 42 profiles of phylogroup Kp3 resulted from horizontal gene transfer (but not large-scale recombination events) from non-KpSC donors (**Table 1; Figure S2; Table S8**). The recombination breakpoints were non-randomly distributed along the genomes: most (109/126, 86.5%) were localized in the second half of the genome (3 Mb – 5.2 Mb of NTUH-K2044 genome coordinates), whereas in the first part (0 – 3 Mb) accounted for only 15 breakpoints.

These 138 hybrid profiles (or with multiple alleles of undefined origins) were therefore discarded during our initial population structure analyses and classification steps, which were based on the remaining 7060 profiles that likely arose from vertical evolution.

##### *Minimum Spanning tree-based clustering of cgMLST profiles: building and assessment*

A pairwise dissimilarity between two cgMLST profiles can be defined by the proportion of loci with two distinct alleles among the loci where alleles are defined in both profiles. A pairwise dissimilarity matrix can be computed from *n* cgMLST profiles, and can be used to build a minimum spanning tree (MStree; *e.g.*, Kruskal, 1956; Prim, 1957a; Dijkstra, 1959), allowing to infer a clustering of the cgMLST profiles, defined by the *k* different connected components obtained by removing from the MStree all edges of length larger than a specified threshold *t*. Such an MStree-based clustering is closely related to the single-linkage classification of the *n* cgMLST profiles (*e.g.*, Gower and Ross, 1969; Johnson, 1967).

In order to determine optimal thresholds *t*, several criteria can be used. Among these criteria, the average silhouette coefficient $S_t$ assesses the ability of an MStree-based clustering to consistently represents in *k* class(es) the 'natural' grouping of the cgMLST profiles (Rousseeuw, 1987; Lengyel and Botta-Dukát, 2019). When $S_t$ is close to 1, the clustering can be considered as accurate. A confidence interval for $S_t$ can be also obtained by considering the distribution of the average silhouette coefficients of different clustering computed from the distance matrix with 'noised' entries.

Further, in order to assess whether an MStree-based clustering $C_t$ (using threshold *t*) is robust to any subsampling biases, a simple approach is to build another MStree-based clustering $C_t'$ from a subsample of cgMLST profiles, and to measure the agreement between the cgMLST profile partitions induced by $C_t'$ and $C_t$. When the level of agreement remains high for different subsampling rates, the corresponding threshold *t* can be considered as being leading to stable clustering. Among different agreement metrics between partitions, the second adjusted Wallace coefficient *w* (Wallace, 1983; Severiano et al., 2011) estimates the probability of observing a pair of profiles in the same class in $C_t$ when they are clustered in the same class in $C_t'$. In order to derive a single coefficient $W_t$ from a range of different subsampling rates *r* (= 10% to 90%), different coefficients *w* were estimated and averaged for each rate *r*; the area under the resulting curve (*i.e.*, rates *r* on X-axis; coefficient *w* on Y-axis) was computed and normalized (using its maximum expected value). Such a normalized area $W_t$ is close to 1 when the different adjusted Wallace coefficients *w* (*i.e.*, derived from varying subsampling rates) are all close to their maximum value, therefore showing that the corresponding MStree-based clustering (based on the threshold *t*) is robust to any subsampling biases. A confidence interval for $W_t$ can be also obtained using the same approach as for $S_t$ (see above).

The MStree-based clustering of cgMLST profiles, as well as the two consistency and stability indices $S_t$ and $W_t$ were implemented in the tool MSTclust (https://gitlab.pasteur.fr/GIPhy/MSTclust). For more details, see https://gitlab.pasteur.fr/GIPhy/MSTclust/-/blob/0.21b/Technical.Notes.pdf).

##### *Nomenclature inheritance algorithm*

In order to attribute to each clonal group (CG), an identifier that would maximally reflect the widely adopted 7-gene ST identifier of the corresponding isolates, we developed a set of naming rules that prioritize the most abundant ST observed among isolates of each CG, as well as some supplementary rules in case of ties. This algorithm is summarized below, whereas an example is given in **Figure S15**.

<u>*Definitions and notations*</u>

Let $G = (U, V, E)$ be a weighted bipartite graph where:

- $U$ is a set of clonal groups (CG) inferred from a cgMLST scheme
- $V$ is the set of sequence types (ST) induced by a MLST scheme
- $E$ is the set of edges $\{u, v\}$ with $u \in U$ and $v \in V$
- $w(\{u, v\})$ is the weight of the edge $\{u, v\}$, *i.e.*, the number of isolates inside $u \cap v$

Let $L(v)$ be the label associated to node $v$ (*i.e.*, the ST identifier), and $L(u)$ the one to determine for $u$ (*i.e.*, the CG identifier).

Let $d_G(v)$ be the degree of a node $v$ inside the graph $G$, *i.e.*, the number of edges incident to node $v$.

Let $s(u) := \Sigma_{v\in V}\, w(\{u, v\})$ be the size of $u$, *i.e.*, the number of strains belonging to the CG $u$.

Let $\Gamma(G)$ be the edge-induced subgraph of a graph $G$ defined by the edge(s) of maximal weight in $G$.

Let $\Delta_G(u)$ be the set of nodes $v$ that are joined to $u$ inside the graph $G$ and of minimum degree, *i.e.*,

$$\Delta_G(u) := \{v' \in V : \{u, v'\} \in E,\ d_G(v') = \min_{\{u, v\}\in E} d_G(v)\}.$$

<u>*Algorithm*</u>

(a) ∘ $\lambda := \max_{v\in V} L(v)$
(b) ∘ **while** $E \neq \emptyset$
  **do**
(c)   ∘ **for each** connected component $G = (U', V', E')$ of $\Gamma(G)$
    **do**
(d)     ∘ **if** $U' = \{\mu$
      **then**
(e)       ∘ $v := \text{argmin}_{v'\in V'} L(v')$
      **else**
(f)       ∘ $U'' := \text{argmin}_{u'\in U'} s(u')$
(g)       ∘ **if** $U'' \neq \{\mu\}$
        **then**
(h)         ∘ $\mu := \text{argmin}_{u''\in U''} L(u'')$
(i)       ∘ $v := \text{argmin}_{v'\in\Delta_{G'}(\mu)} L(v')$
(j)     ∘ $L(\mu) := L(v)$
(k)     ∘ removing $\mu$ and v from $G$, as well as nodes $v$ such that $w(\{\mu, v\}) = w(\{\mu, v\})$
(l) ∘ **for each** $\mu \in U$
  **do**
(l)   ∘ $\lambda := \lambda + 1$
(m)   ∘ $L(\mu) := \lambda$

##### *Impact of strains input order on LIN codes, and use of Prim's algorithm*

By design, the input order of genomes into the cgLINcode nomenclature system influences their attributed code, as is the case for the original LIN code system (Marakeby et al., 2014). We evaluated this impact by quantifying the variation in the number of partitions at a given threshold, as defined by the number of distinct prefixes: for each threshold varying from 1% to 99%, a LIN encoding was defined using this threshold, and the 7,060 high-quality, non-hybrid cgMLST profiles were encoded 500 times with random input orders. This experiment made it possible to determine (i) the threshold values associated with a stronger variability in the final number of values; and (ii) the magnitude of this variability. In the example illustrated in **Figure S10**, we observed that the number of distinct prefixes was affected by the order of encoding, especially in the 450 - 530 mismatches range. Note that this experiment can help to select position thresholds, for example by favoring those that minimize the variance of the number of partitions (*i.e.*, are less affected by input order).

We next sought to minimize this problem by defining an optimal input order. The one that answered our expectations is the input order guided by a Prim's algorithm (Prim, 1957). More precisely, the number of categories in a given LIN encoding bin is minimal (*i.e.*, identical to the number of groups created by a single-linkage clustering using the threshold associated to the bin) when the profiles are encoded following the order induced by the traversal of an MStree. Indeed, when following such an order, when a new profile is considered for encoding, then its closest profile is already encoded (by definition of a tree traversal). The optimal order we suggest is therefore verified by noting that the Prim's (1957) algorithm to infer a MStree induces such an MStree traversal. A comparison between the MLSL approach and the cgLIN codes was performed (cgLIN codes in optimal *versus* arbitrary order). We found that the partitioning created by the MLSL approach and that created by the cgLIN codes according to the optimal order, were identical (**Table S1**).

We generated 500 random input orders and then generated cgLIN codes with two bins (the first varies from 1 to 100% with a step of one allelic difference, i.e. 1/629*100, and the second fixed at 100%). Then we counted the number of prefixes, up to the first bin, that were created. We observed that the 10 identifier bins differ in their sensitivity to input order (**Figure S10**). The most affected bins correspond to regions of the pairwise distance distribution with high density; in particular around 485 mismatches, before the mode that corresponds to inter-sublineage distances.

The algorithm below was used to define the input order, without even having to construct an MStree. Indeed, thanks to a simple traversal of the matrix of dissimilarities between profiles, the algorithm makes it possible to quickly determine the optimal order for LIN encoding.

*Algorithm*

(a) ◦ Create a set "mstSet" that keeps track of vertices already included in MST

(b) ◦ Assign a key value to all vertices in the input graph. Initialize all key values as $\infty$. Assign key value as $0$ for the first vertex so that it is picked first.

(c) ◦ **while** "mstSet" doesn't include all vertices

(d) ◦ Pick a vertex u which is not there in "mstSet" and has minimum key value.

(e) ◦ Include u to "mstSet".

(f) ◦ Update key value of all adjacent vertices of u. To update the key values, iterate through all adjacent vertices. For every adjacent vertex $v$, if weight of edge $u - v$ is less than the previous key value of $v$, update the key value as weight of $u - v$.

Note that using key values enables to pick the minimum weight edge from cut. The key values are used only for vertices which are not yet included in MStree; the key value for these vertices indicate the minimum weight edges connecting them to the set of vertices included in MStree. The time complexity required by Prim's (1957) algorithm is $O(E\ log\ V)$.

#### *List of supplementary figures*


Figure S1. cgMLST profile painting illustrates large recombinations

Figure S2. Genomes inclusion flowchart

Figure S3. Characteristics of the 629 loci of the cgMLST scheme

Figure S4. The distribution of pairwise distances based on Average Nucleotide Identity (ANI) and cgMLST

Figure S5. Details of cgMLST pairwise distances distributions

Figure S6. Correspondence of ST, sublineage and clonal group classifications for 9 major K. pneumoniae sublineages

Figure S7. The distribution of pairwise distances based on Average Nucleotide Identity (ANI) and cgMLST, with hybrid genomes

Figure S8. Impact of inter-phylogroup hybrid genomes on cgMLST classification groups

Figure S9. Virulence and resistance scores in major sublineages

Figure S10. Impact of input order on the number of partitions in the resulting LIN codes

Figure S11. Relationships between ST, cgMLST and cgLIN codes, and their behavior upon novel genomes inclusion

Figure S12. Distribution of the phylogroup homogeneity index

Figure S13. Principle of cgLIN code implementation

Figure S14. cgLIN codes implementation for nearly-identical cgMLST profiles

Figure S15. Step-by-step illustration of the taxonomic inheritance algorithm

#### *List of supplementary tables*

Table S1. Dataset of 7,433 genomes

Table S2. Hybrid genomes breakdown by phylogroup

Table S3. Characteristics of the 629 loci of the cgMLST scheme

Table S4. Correspondence between SLs, CGs and STs

Table S5. Characteristics of the clonal groups

Table S6. Outbreak dataset

Table S7. Correspondence between ANI and cgMLST distance thresholds

Table S8. Reference genomes

##### *Supplementary references*


Dijkstra, E.W. (1959). A note on two problems in connexion with graphs. Numer. Math. *1*, 269–271.

Gower, J.C., and Ross, G.J.S. (1969). Minimum Spanning Trees and Single Linkage Cluster Analysis. Applied Statistics *18*, 54.

Holt, K.E., Wertheim, H., Zadoks, R.N., Baker, S., Whitehouse, C.A., Dance, D., Jenney, A., Connor, T.R., Hsu, L.Y., Severin, J., et al. (2015). Genomic analysis of diversity, population structure, virulence, and antimicrobial resistance in Klebsiella pneumoniae, an urgent threat to public health. Proc Natl Acad Sci U S A *112*, E3574-81.

Jain, C., Rodriguez-R, L.M., Phillippy, A.M., Konstantinidis, K.T., and Aluru, S. (2018). High throughput ANI analysis of 90K prokaryotic genomes reveals clear species boundaries. Nat Commun *9*, 5114.

Johnson, S.C. (1967). Hierarchical clustering schemes. Psychometrika *32*, 241–254.

Kruskal, J.B. (1956). On the shortest spanning subtree of a graph and the traveling salesman problem. Proc. Amer. Math. Soc. *7*, 48–48.

Lengyel, A., and Botta-Dukát, Z. (2019). Silhouette width using generalized mean—A flexible method for assessing clustering efficiency. Ecol Evol *9*, 13231–13243.

Marakeby, H., Badr, E., Torkey, H., Song, Y., Leman, S., Monteil, C.L., Heath, L.S., and Vinatzer, B.A. (2014). A system to automatically classify and name any individual genome-sequenced organism independently of current biological classification and nomenclature. PLoS One *9*, e89142.

Prim, R.C. (1957). Shortest Connection Networks And Some Generalizations. Bell System Technical Journal *36*, 1389–1401.

Rousseeuw, P.J. (1987). Silhouettes: A graphical aid to the interpretation and validation of cluster analysis. Journal of Computational and Applied Mathematics *20*, 53–65.

Severiano, A., Pinto, F.R., Ramirez, M., and Carriço, J.A. (2011). Adjusted Wallace Coefficient as a Measure of Congruence between Typing Methods. J. Clin. Microbiol. *49*, 3997–4000.

Wallace, D.L. (1983). A Method for Comparing Two Hierarchical Clusterings: Comment. Journal of the American Statistical Association *78*, 569–576.

#### Figure S1. cgMLST profile painting illustrates large recombinations

The 7198 total (upper right) and 138 hybrid (bottom) cgMLST profiles are represented. Loci are represented in their order along the reference genome NTUH-K2044. Each allele is colored according to its attribution to a phylogroup (see color key; white: unattributed). Gene loci are indicated at the bottom of the figure. Blocs with distinctive colors within some profiles correspond to large recombination events.

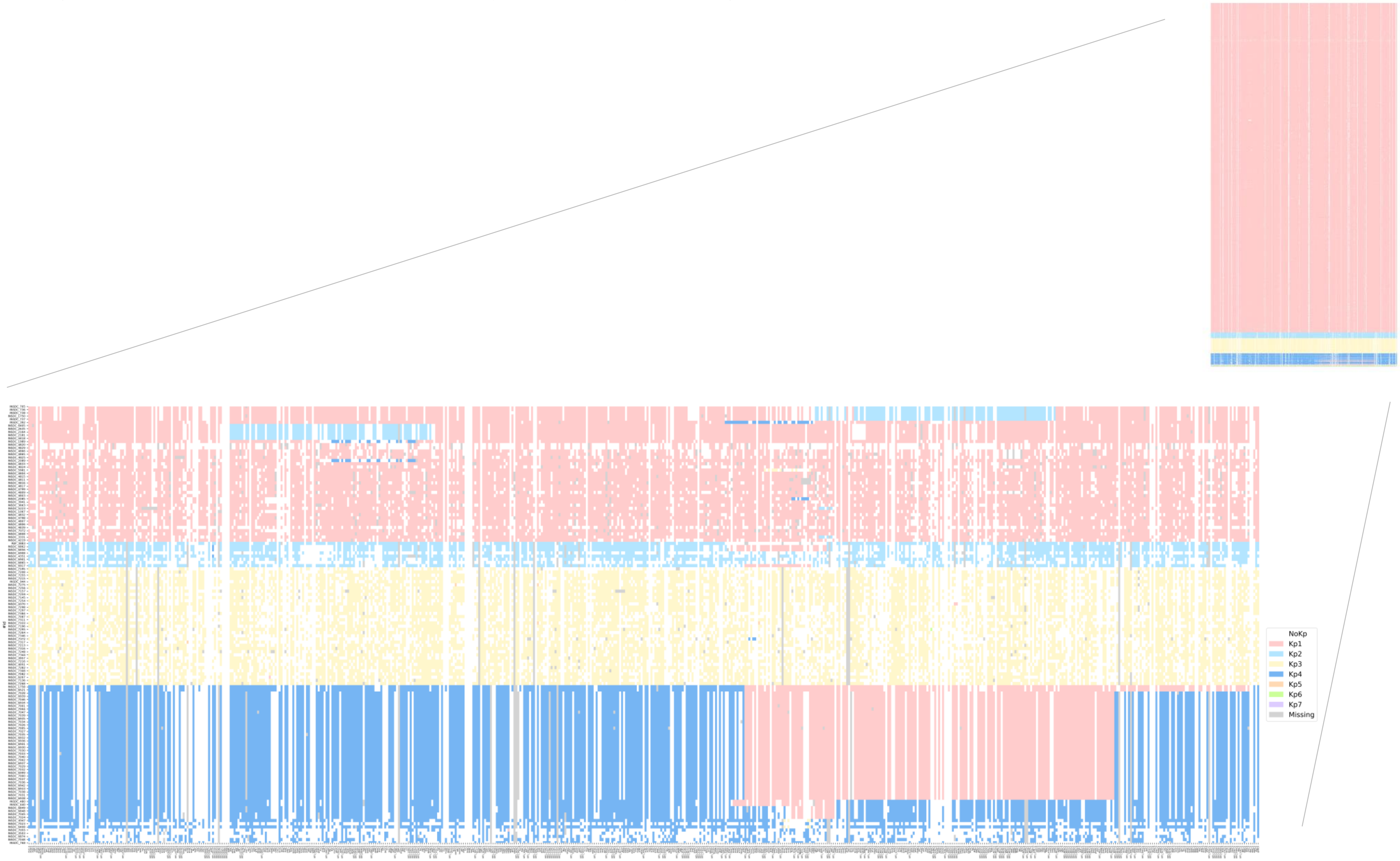

### Figure S2. Genome inclusion flowchart

The flowchart summarizes the genomes inclusion process.

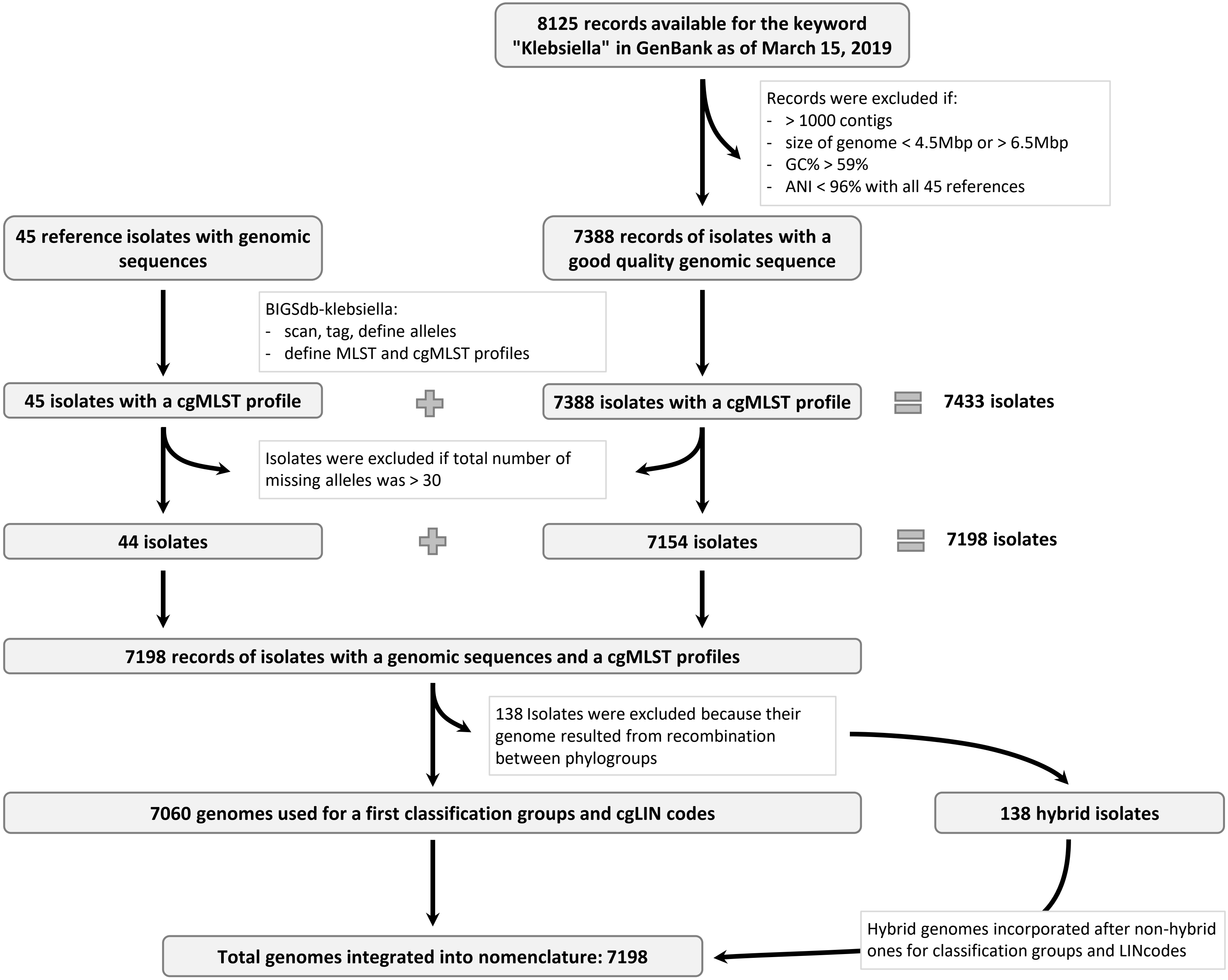

**Figure S3. Characteristics of the 629 loci of the cgMLST scheme**

(A) Effect of locus size on allele number. Central panel: each point represents a locus; the X-axis corresponds to the locus length and the Y-axis to the number of alleles; top/bottom and right/left panels: the distributions and the boxplots of each parameter are shown. (B) Effect of intra-gene recombination on allele diversity. Boxplots show the distribution of the number of alleles according to the different significance levels of the PHI statistic. n.a.: not applicable (polymorphism was too low); n.s.: loci for which no significant recombination was detected.

**A. Locus length *versus* number of alleles**

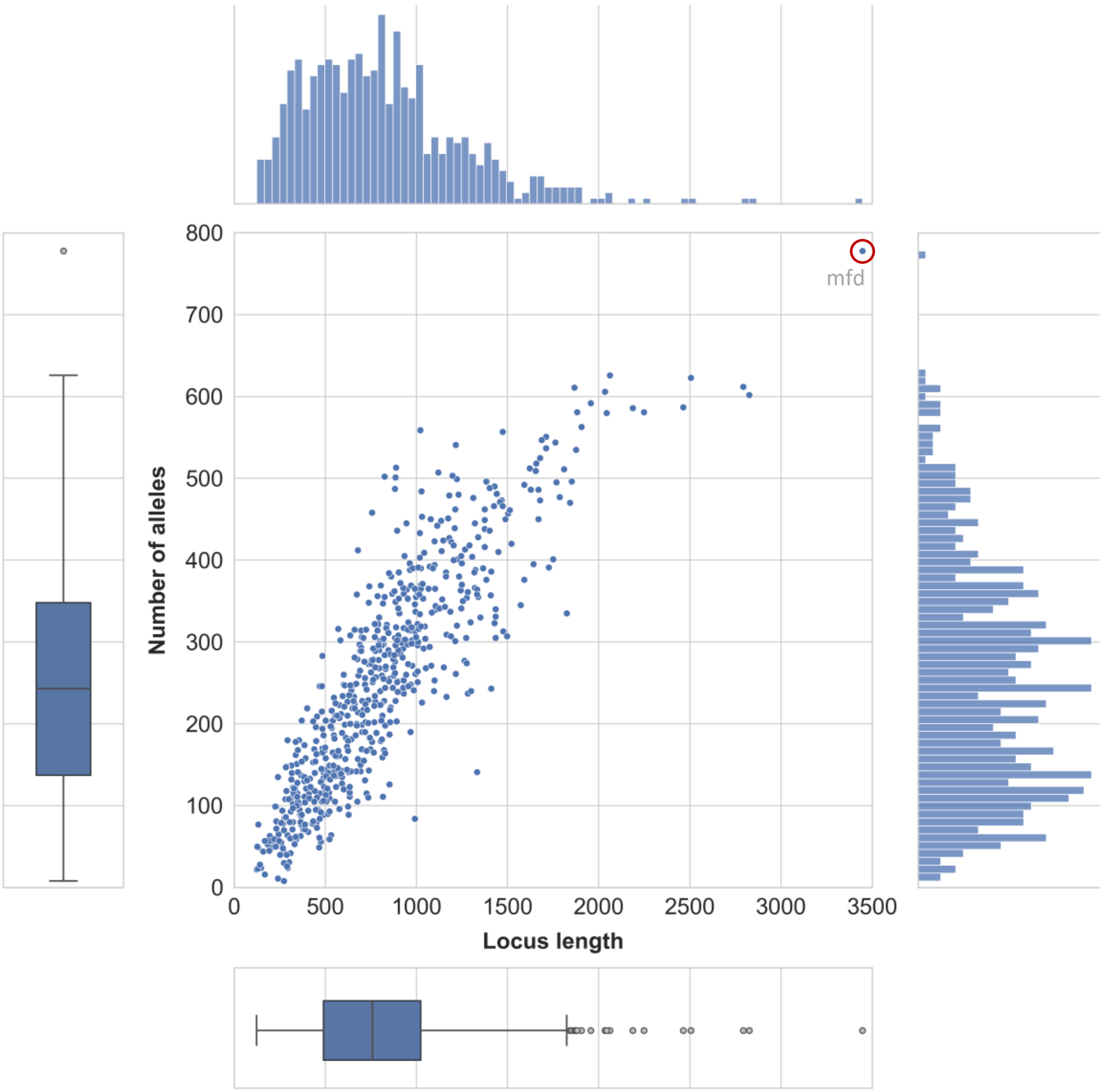


**B. Effect of recombination on allele number per locus**

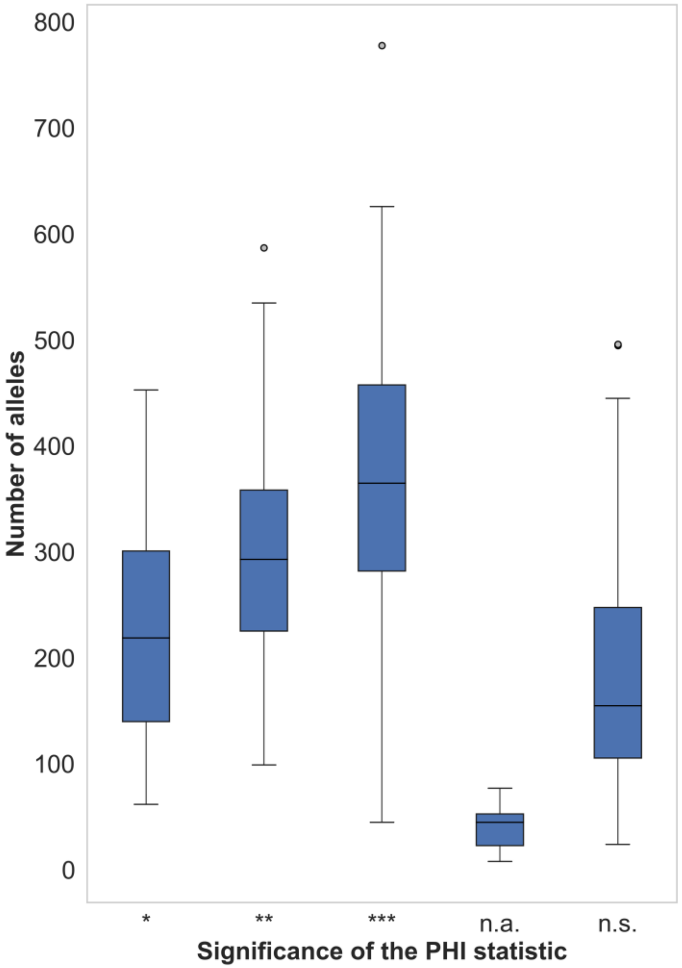

**Figure S4. The distribution of pairwise distances based on Average Nucleotide Identity (ANI) and cgMLST**

(A) cgMLST similarity versus ANI. In the central panel, each point represents a pair of strains; the X-axis corresponds to the cgMLST profile similarity whereas the Y-axis corresponds to the ANI; the corresponding density distributions are shown on the outside of the graph. Colors correspond to inter-phylogroup (orange) and intra-phylogroup (blue) genome pairs. (B) Distribution of ANI values with box-plots of phylogroup comparisons. (C) Distribution of cgMLST similarity values with box-plots of phylogroup comparisons.

**A. cgMLST similarity *versus* ANI**

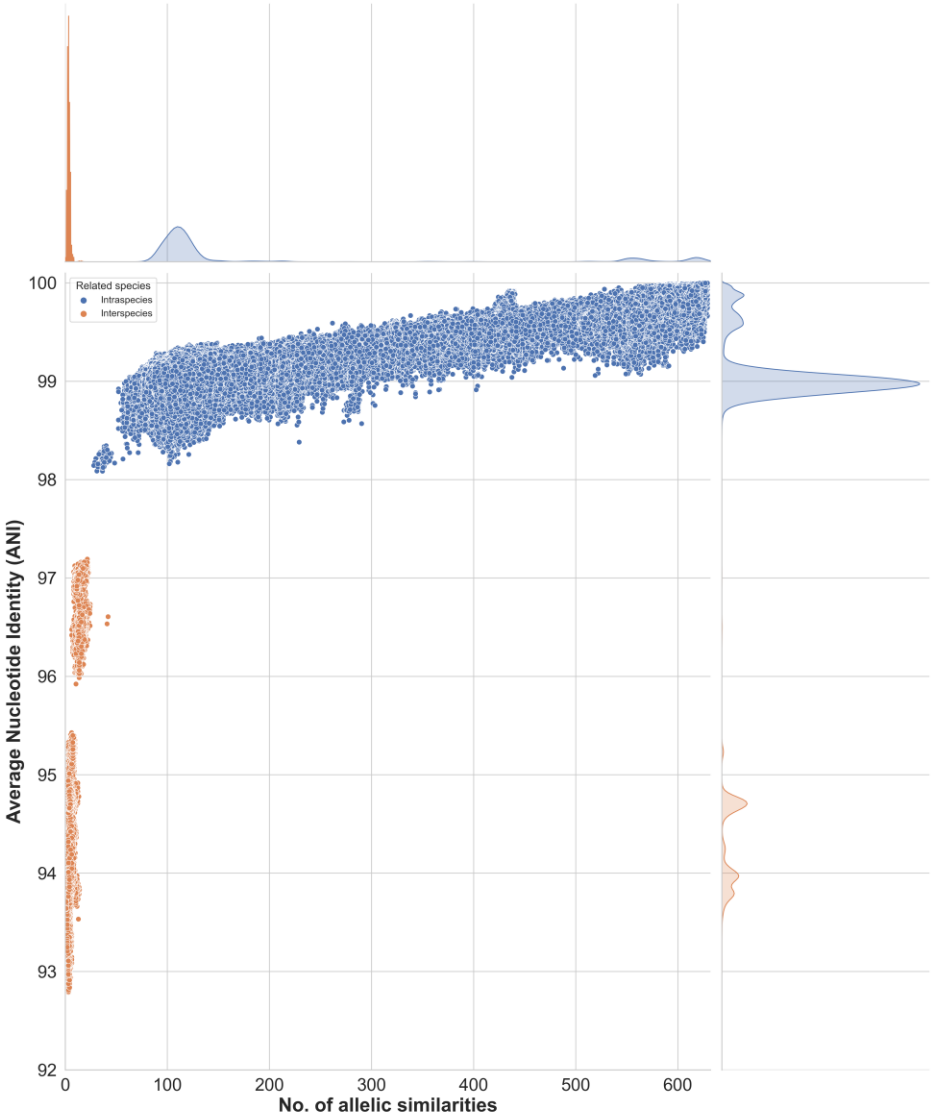


**B. ANI**

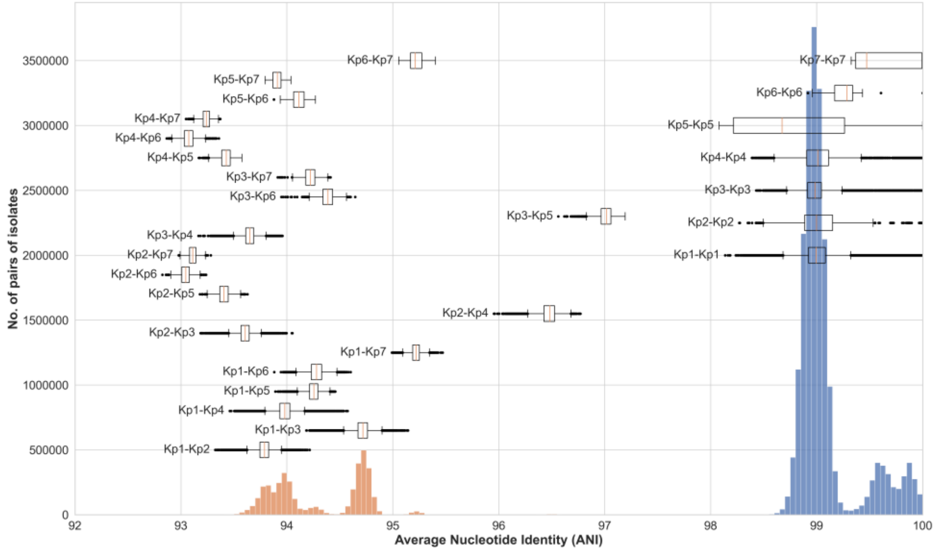


**C. cgMLST**

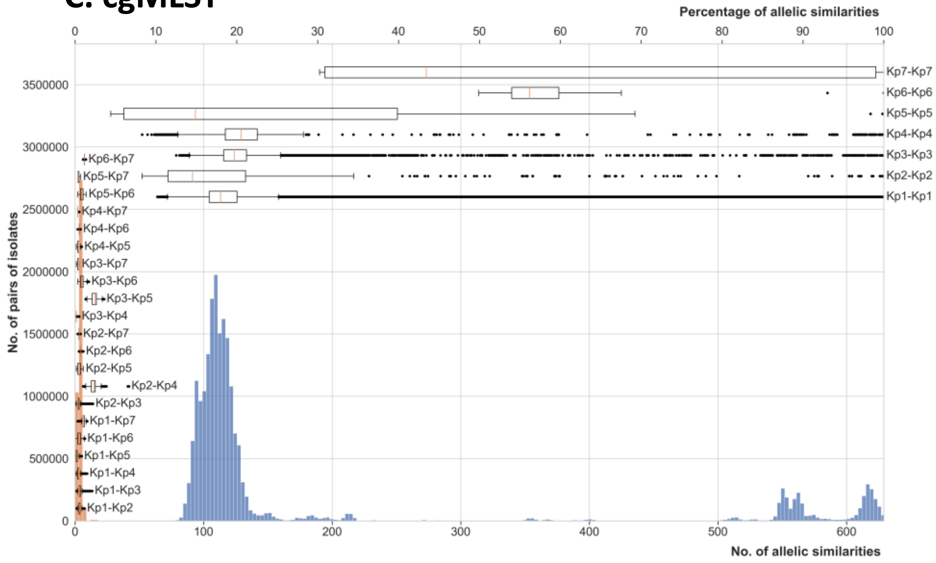

### Figure S5. Details of cgMLST pairwise distances distributions

(A) cgMLST pairwise distance distribution, with a zoom on the range of 575-605 allelic differences (inter-subspecies comparisons). The chosen threshold of 585 for subspecies delineation is indicated by a red vertical line. (B) cgMLST pairwise distance distribution, with a zoom on the range of 40-100 allelic differences. Sectors of the bars are colored according to the 7-gene MLST sequence type (ST) of the compared genomes (see key). The 79-mismatch mode of the distribution is mostly composed of ST258-ST11 pairs.

Distribution of cgMLST profiles pairwise distance of all dataset (A) and cgMLST profiles pairwise distance with ST repartitions (B). Note that the 79 allelic mismatches mode comprised a large proportion (48.0%) of ST258-ST11 comparisons. This observation is consistent with ST258 having evolved through a 1.1 MB large-scale recombination event of an ST11 ancestor with a ST442 donor (Chen et al., 2014): these two STs differ by 57 cgMLST alleles in this 1.1 MB region (computed for GCA_000445405.1 JM45 and SB4938_Kp13), and 21 additional allelic mismatches are observed on average between ST258 (GCA_000597905.1 NJST258_2) and the ST11 (GCA_000445405.1 JM45) ancestor outside the recombined region. Other important contributors to this third mode were comparisons between ST11-ST512, ST258-ST340, ST258-ST437.

### Figure S5. Details of cgMLST pairwise distances distributions

#### A. cgMLST pairwise distance distribution, with a zoom in range 575-605 allelic differences

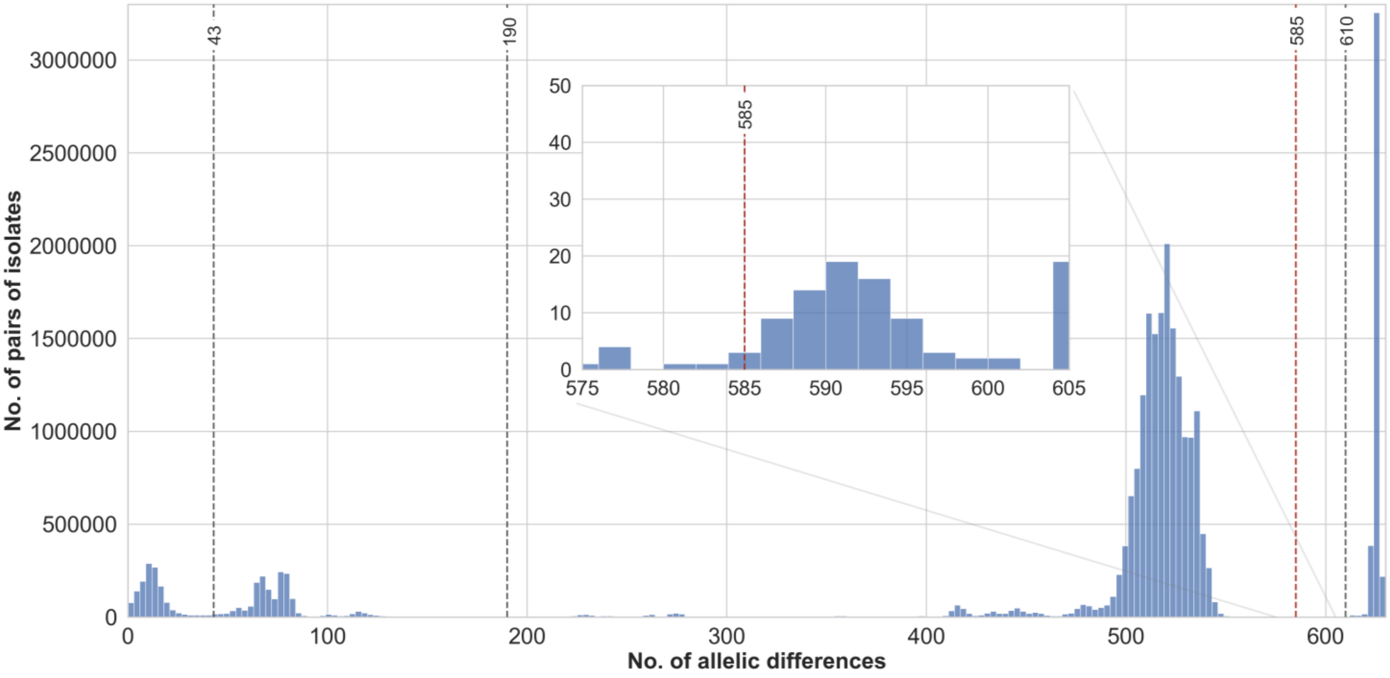


#### B. cgMLST pairwise distance distribution, with a zoom in range 40-100 allelic differences

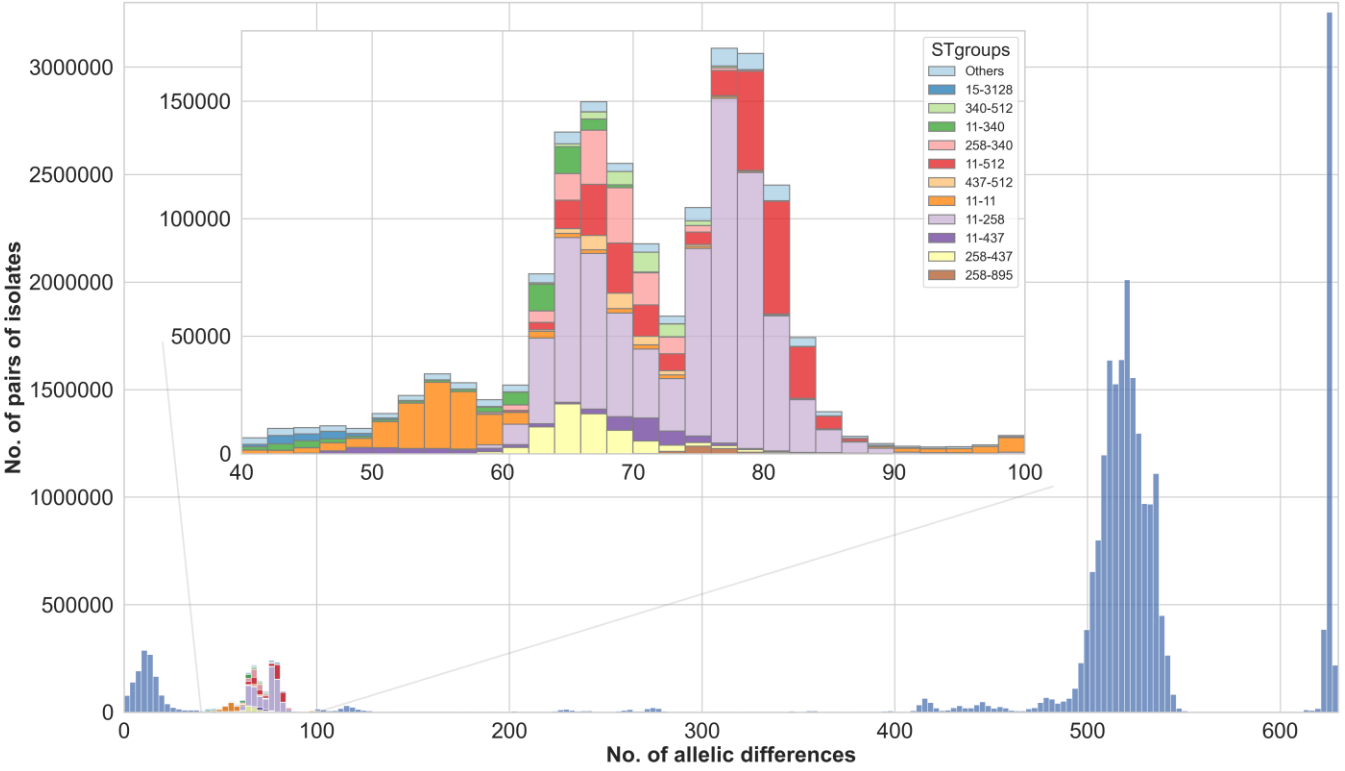

### Figure S6. Correspondence of ST, sublineage and clonal group classifications for 9 major K. pneumoniae sublineages

ST: 7-gene MLST sequence type. The identifiers of the sublineages and clonal groups classifications are those inherited from MLST using our mapping algorithm.

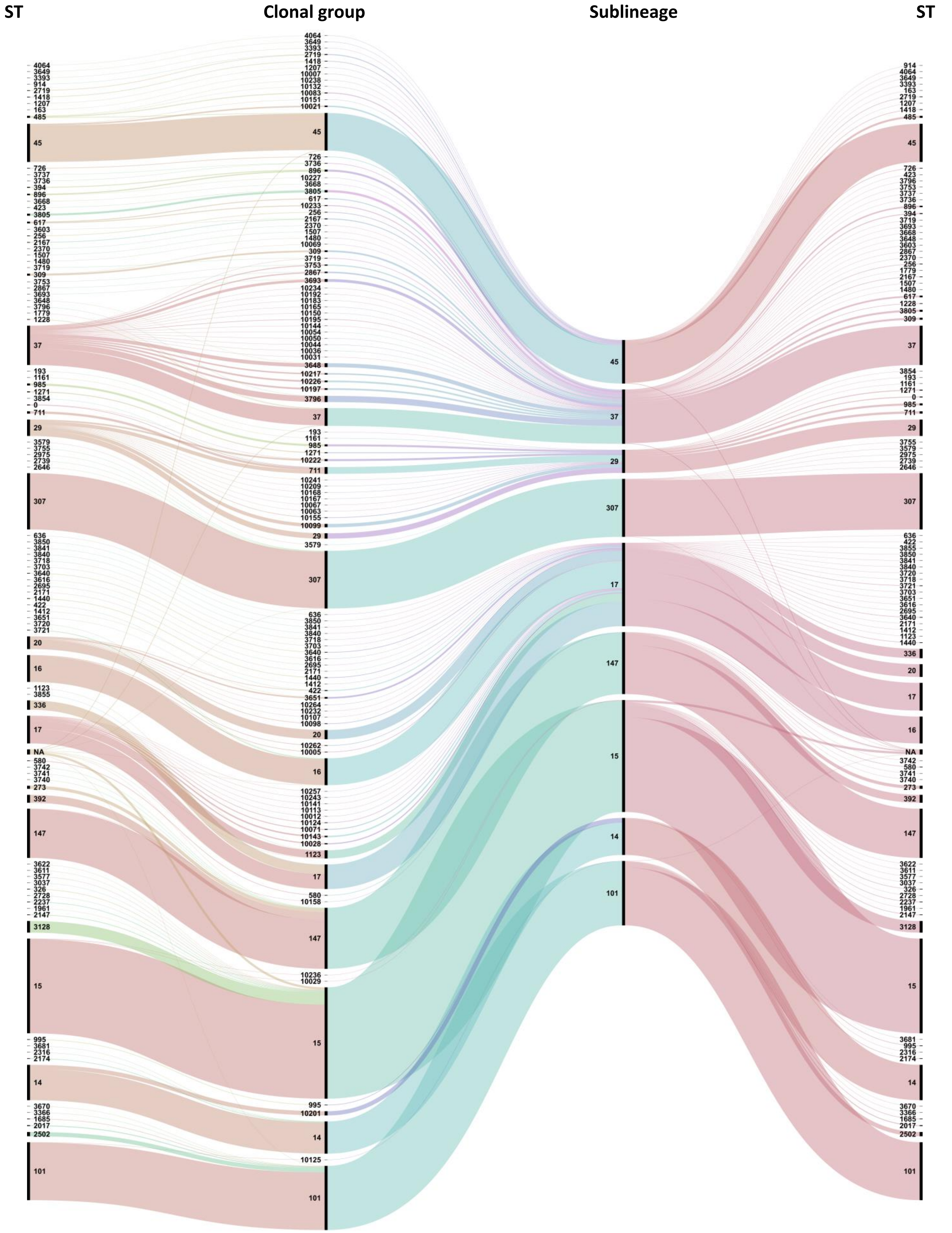

**Figure S7. The distribution of pairwise distances based on Average Nucleotide Identity (ANI) and cgMLST, with hybrid genomes**

(A) cgMLST similarity versus ANI. In the central panel, each point represents a pair of strains; the X-axis corresponds to the cgMLST profile similarity whereas the Y-axis corresponds to the ANI; the corresponding density distributions are shown on the outside of the graph. Colors correspond to genome pairs involving only non-hybrid genomes (green) or at least one hybrid genome (red). (B) Distribution of ANI values. (C) Distribution of cgMLST similarity values.

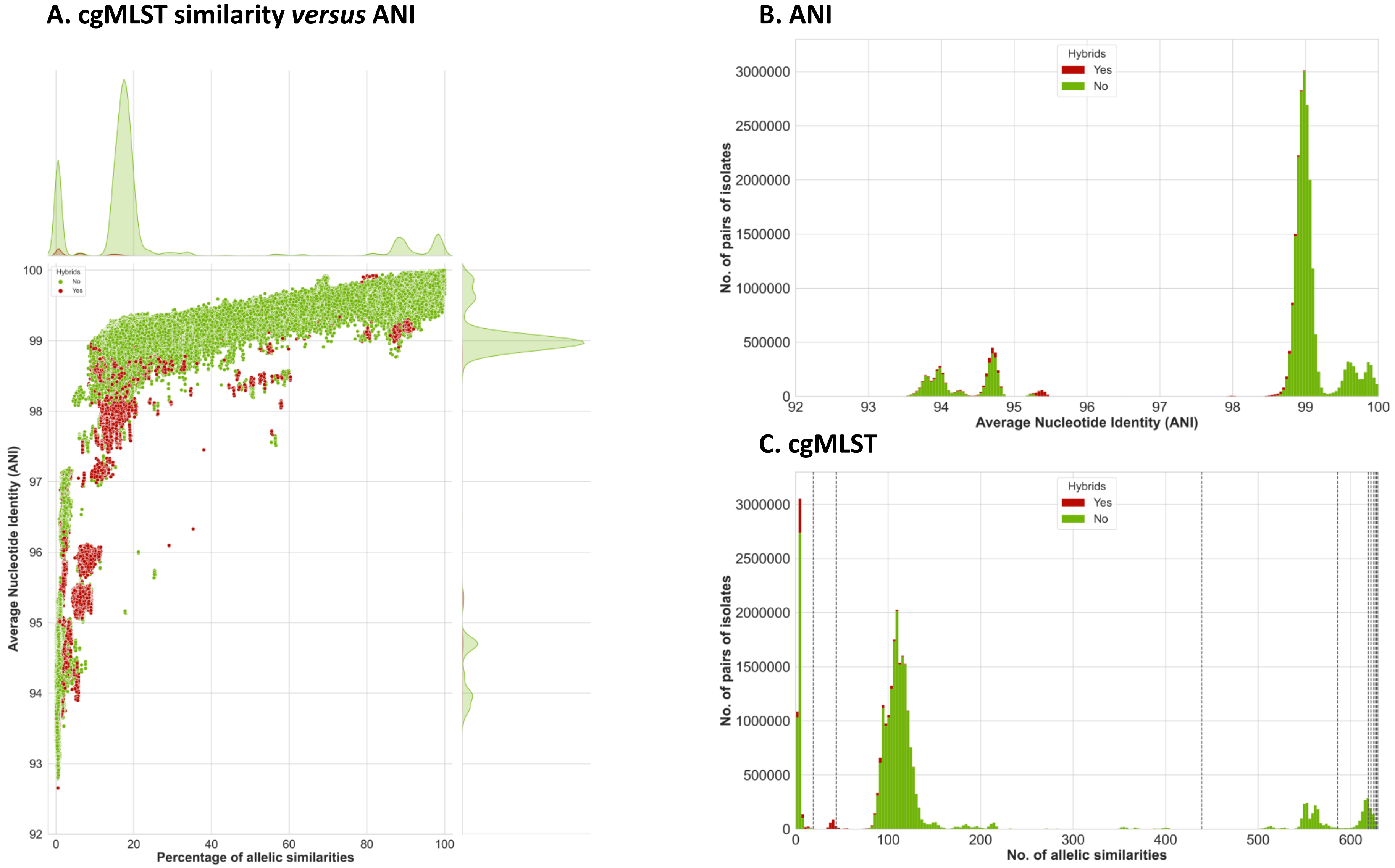

#### Figure S8. Impact of inter-phylogroup hybrid genomes on cgMLST classification groups

The effect of incorporating the hybrid genomes on MLSL and cgLIN codes is illustrated by comparing example genome codes before (left) and after (right) hybrid were included into the nomenclature. Version 1 of nomenclature was created from the 7060 non-hybrid genomes; version 2 was obtained after the 138 hybrid genomes were included. Note that the example MLSL genome codes are unstable, as the 610 and 585 threshold levels were affected by the incorporation of hybrids; in contrast, cgLIN codes were unaffected, as expected by design.

### Figure S8. Impact of inter-phylogroup hybrid genomes on cgMLST classification groups

#### Version 1

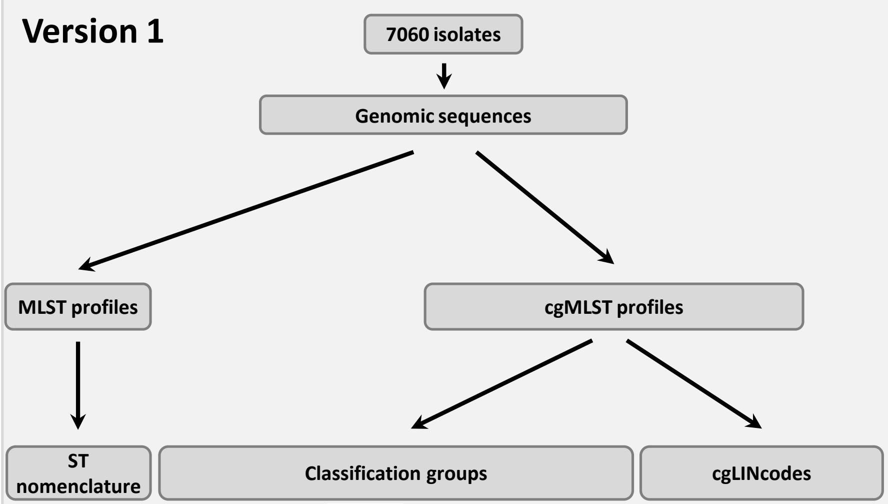


| Isolates | ST (MLST) |
|---|---|
| SB20 | 15 |
| SB1067 | 17 |
| SB4767 | 1220 |
| SB48 | 2263 |
| SB5531 | 2561 |
| SB5544 | 2605 |
| SB2110 | 526 |
| SB98 | 622 |
| SB30 | 1215 |
| SB4697 | 2275 |
| SB6095 | 3303 |
| SB6094 | 2830 |
| FF1003 | 4938 |
| FF1006 | 4938 |

| | | | SL | CG | | | | | | |
|---|---|---|---|---|---|---|---|---|---|---|
| Threshold | 610 | 585 | 190 | 43 | 10 | 7 | 4 | 2 | 1 | 0 |
| SB20 | 1 | 1 | 1 | 1 | 1 | 1 | 1 | 1 | 1 | 1 |
| SB1067 | 1 | 1 | 4 | 4 | 4 | 4 | 4 | 4 | 4 | 4 |
| SB4767 | 4 | 4 | 35 | 35 | 35 | 35 | 35 | 35 | 35 | 35 |
| SB48 | 4 | 4 | 18 | 18 | 18 | 18 | 18 | 18 | 18 | 18 |
| SB5531 | 4 | 6 | 27 | 27 | 27 | 27 | 27 | 27 | 27 | 27 |
| SB5544 | 4 | 6 | 28 | 28 | 28 | 28 | 28 | 28 | 28 | 28 |
| SB2110 | 2 | 5 | 21 | 21 | 21 | 21 | 21 | 21 | 21 | 21 |
| SB98 | 2 | 5 | 20 | 20 | 20 | 20 | 20 | 20 | 20 | 20 |
| SB30 | 2 | 2 | 14 | 14 | 14 | 14 | 14 | 14 | 14 | 14 |
| SB4697 | 2 | 2 | 23 | 23 | 23 | 23 | 23 | 23 | 23 | 23 |
| SB6095 | 3 | 3 | 33 | 33 | 33 | 33 | 33 | 33 | 33 | 33 |
| SB6094 | 3 | 3 | 34 | 34 | 34 | 34 | 34 | 34 | 34 | 34 |
| FF1006 | 7 | 8 | 36 | 36 | 36 | 36 | 37 | 37 | 37 | 37 |
| FF1003 | 7 | 8 | 36 | 36 | 36 | 36 | 37 | 37 | 37 | 37 |

| Isolates | cgLINcodes |
|---|---|
| SB20 | 0.0.0.0.0.0.0.0.0.0 |
| SB1067 | 0.0.22.12.0.1.0.0.0.0 |
| SB4767 | 1.0.6.0.0.0.0.0.0.0 |
| SB48 | 1.0.65.0.0.0.0.0.0.0 |
| SB5531 | 1.1.0.0.0.0.0.0.0.0 |
| SB5544 | 1.1.1.0.0.0.0.0.0.0 |
| SB2110 | 2.0.0.1.0.0.0.0.0.0 |
| SB98 | 2.0.8.1.0.0.0.0.0.0 |
| SB30 | 2.1.22.0.0.0.0.0.0.0 |
| SB4697 | 2.1.23.0.0.0.0.0.0.0 |
| SB6095 | 3.0.0.0.0.0.0.0.0.0 |
| SB6094 | 3.0.3.0.0.0.0.0.0.0 |
| FF1006 | 4.0.0.0.0.0.0.0.0.0 |
| FF1003 | 4.0.0.0.0.0.0.0.0.0 |

Unstable

Stable

#### Version 2

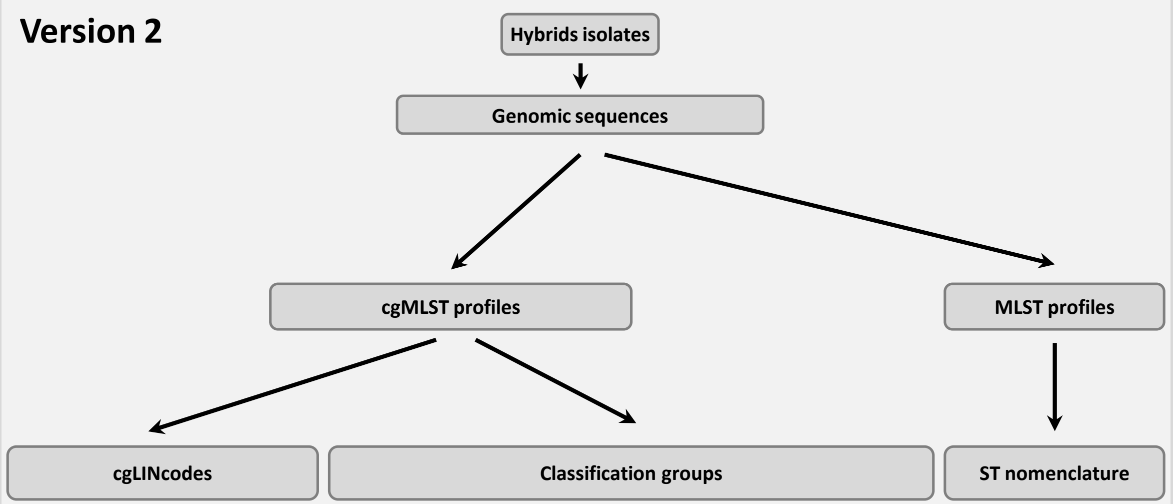


| Isolates | cgLINcodes |
|---|---|
| SB20 | 0.0.0.0.0.0.0.0.0.0 |
| SB1067 | 0.0.22.12.0.1.0.0.0.0 |
| SB4767 | 1.0.6.0.0.0.0.0.0.0 |
| SB48 | 1.0.65.0.0.0.0.0.0.0 |
| SB5531 | 1.1.0.0.0.0.0.0.0.0 |
| SB5544 | 1.1.1.0.0.0.0.0.0.0 |
| SB2110 | 2.0.0.1.0.0.0.0.0.0 |
| SB98 | 2.0.8.1.0.0.0.0.0.0 |
| SB30 | 2.1.22.0.0.0.0.0.0.0 |
| SB4697 | 2.1.23.0.0.0.0.0.0.0 |
| SB6095 | 3.0.0.0.0.0.0.0.0.0 |
| SB6094 | 3.0.3.0.0.0.0.0.0.0 |
| FF1006 | 4.0.0.0.0.0.0.0.0.0 |
| FF1003 | 4.0.0.0.0.0.0.0.0.0 |
| SB1124 | 2.0.32.1.0.0.0.0.0.0 |
| GCA_900494485 | 0.0.105.17.0.0.0.0.0.0 |
| GCA_900492755 | 2.1.36.1.0.0.0.0.0.0 |
| GCA_001874875 | 0.0.440.0.0.0.0.0.0.0 |
| GCA_002752865 | 2.1.61.0.0.0.0.0.0.0 |

| | | | SL | CG | | | | | | |
|---|---|---|---|---|---|---|---|---|---|---|
| Threshold | 610 | 585 | 190 | 43 | 10 | 7 | 4 | 2 | 1 | 0 |
| SB20 | 1 | 1 | 1 | 1 | 1 | 1 | 1 | 1 | 1 | 1 |
| SB1067 | 1 | 1 | 4 | 4 | 4 | 4 | 4 | 4 | 4 | 4 |
| SB4767 | 1 | 4 | 35 | 35 | 35 | 35 | 35 | 35 | 35 | 35 |
| SB48 | 1 | 4 | 18 | 18 | 18 | 18 | 18 | 18 | 18 | 18 |
| SB5531 | 1 | 6 | 27 | 27 | 27 | 27 | 27 | 27 | 27 | 27 |
| SB5544 | 1 | 6 | 28 | 28 | 28 | 28 | 28 | 28 | 28 | 28 |
| SB2110 | 1 | 1 | 21 | 21 | 21 | 21 | 21 | 21 | 21 | 21 |
| SB98 | 1 | 1 | 20 | 20 | 20 | 20 | 20 | 20 | 20 | 20 |
| SB30 | 1 | 1 | 14 | 14 | 14 | 14 | 14 | 14 | 14 | 14 |
| SB4697 | 1 | 1 | 23 | 23 | 23 | 23 | 23 | 23 | 23 | 23 |
| SB6095 | 3 | 3 | 33 | 33 | 33 | 33 | 33 | 33 | 33 | 33 |
| SB6094 | 3 | 3 | 34 | 34 | 34 | 34 | 34 | 34 | 34 | 34 |
| FF1006 | 7 | 8 | 36 | 36 | 36 | 36 | 37 | 37 | 37 | 37 |
| FF1003 | 7 | 8 | 36 | 36 | 36 | 36 | 37 | 37 | 37 | 37 |
| SB1124 | 1 | 1 | 548 | 1176 | 2138 | 2401 | 2865 | 3471 | 3987 | 4903 |
| GCA_900494485 | 1 | 1 | 57 | 1177 | 2139 | 2402 | 2866 | 3472 | 3988 | 4904 |
| GCA_900492755 | 1 | 1 | 138 | 1178 | 2140 | 2403 | 2867 | 3473 | 3989 | 4905 |
| GCA_001874875 | 1 | 1 | 720 | 1179 | 2141 | 2404 | 2868 | 3474 | 3990 | 4906 |
| GCA_002752865 | 1 | 1 | 723 | 1183 | 2145 | 2408 | 2872 | 3478 | 3994 | 4911 |

| Isolates | ST (MLST) |
|---|---|
| SB20 | 15 |
| SB1067 | 17 |
| SB4767 | 1220 |
| SB48 | 2263 |
| SB5531 | 2561 |
| SB5544 | 2605 |
| SB2110 | 526 |
| SB98 | 622 |
| SB30 | 1215 |
| SB4697 | 2275 |
| SB6095 | 3303 |
| SB6094 | 2830 |
| FF1003 | 4938 |
| FF1006 | 4938 |
| SB1124 | 2274 |
| GCA_900494485 | 2948 |
| GCA_900492755 | 1584 |
| GCA_001874875 | 34 |
| GCA_002752865 | 2542 |

#### Figure S9. Virulence and resistance scores of major sublineages

Left: Heatmap of the number of strains classified by virulence and resistance scores, broken down by sublineage; middle panel: Median of the virulence and resistance scores (the scale for the virulence score is negativized). Right panel: number of strains present in each sublineage

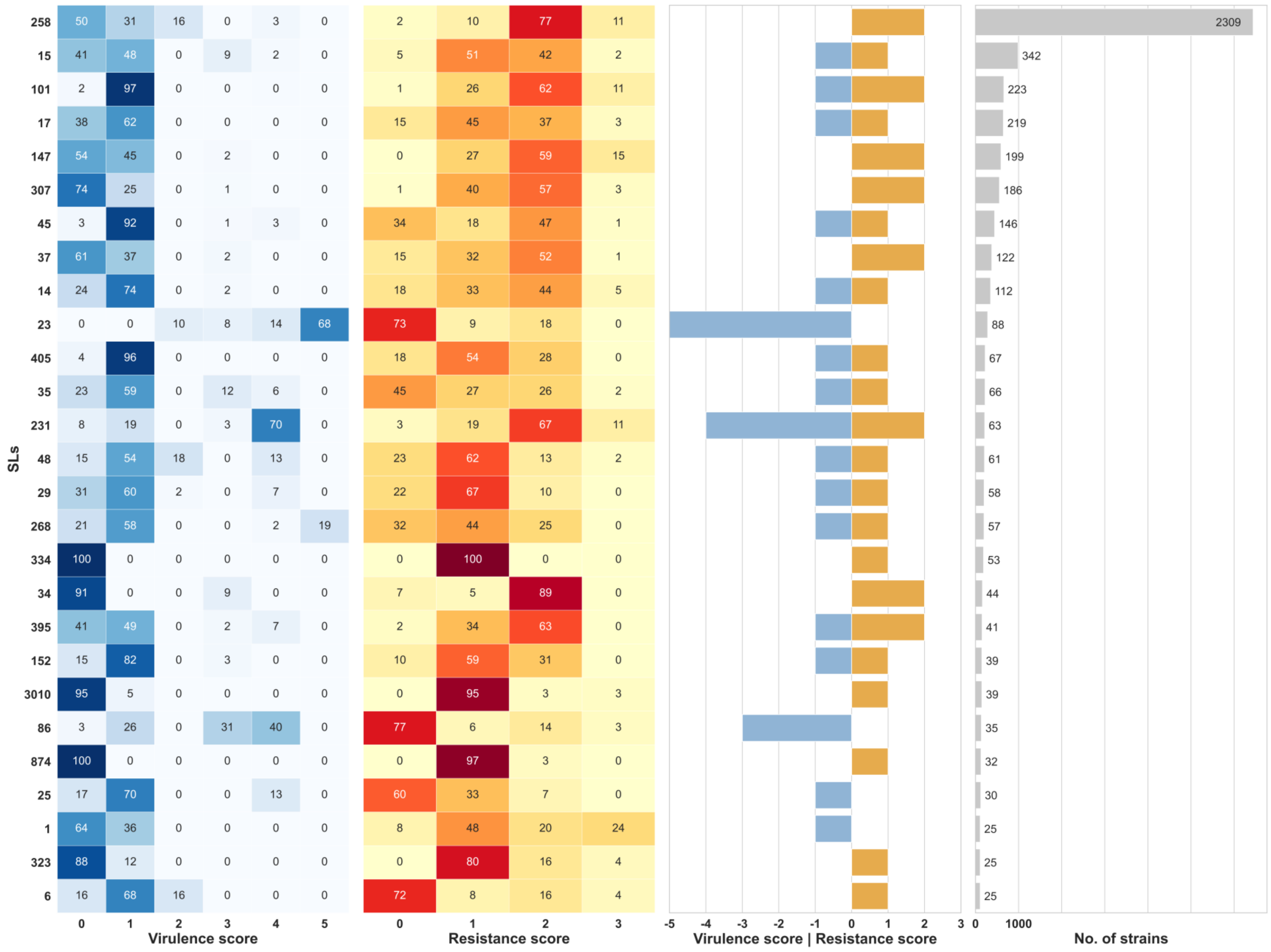

**Figure S10. Impact of input order on the number of partitions in the resulting LIN codes**

The variation in the number of partitions at a given threshold, as defined by the number of distinct prefixes, was quantified. For each threshold ranging from 1% to 99%, a cgLIN encoding was defined, and the 7,060 high-quality non-hybrid cgMLST profiles were encoded 500 times with random input orders. Blue: number of partitions created; red: the variance of this number.

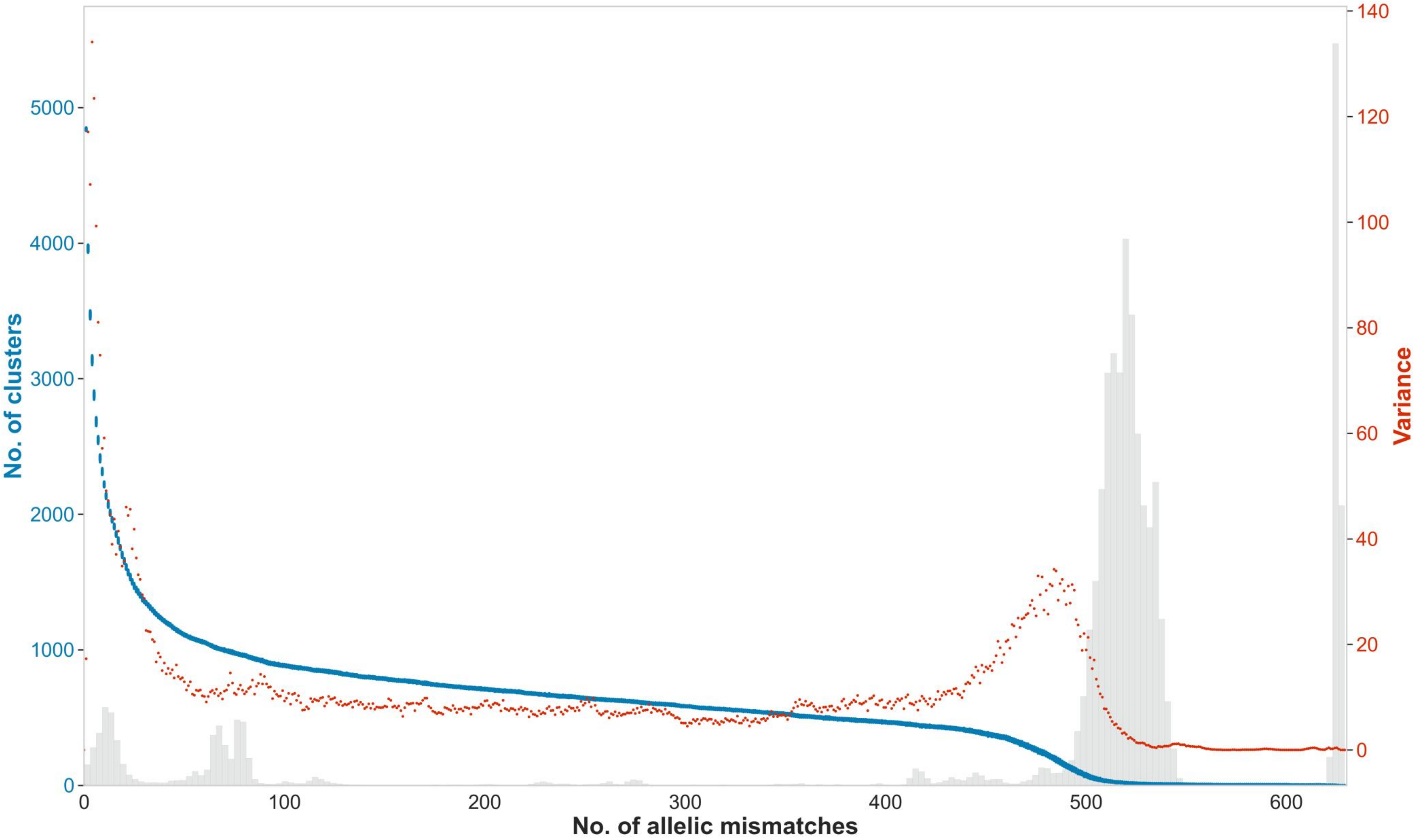

#### Figure S11. Relationships between ST, cgMLST and cgLIN codes, and their behavior upon novel genomes inclusion

From the cgMLST profiles, MLSL classification groups and cgLIN codes are generated. On the right, a second batch was submitted, leading to MLST classification groups to evolve, unlike the cgLIN codes, which are stable. A table of correspondence between MLSL classification and cgLIN codes can be used to follow MLSL classifications attached to each cgLIN code over time.

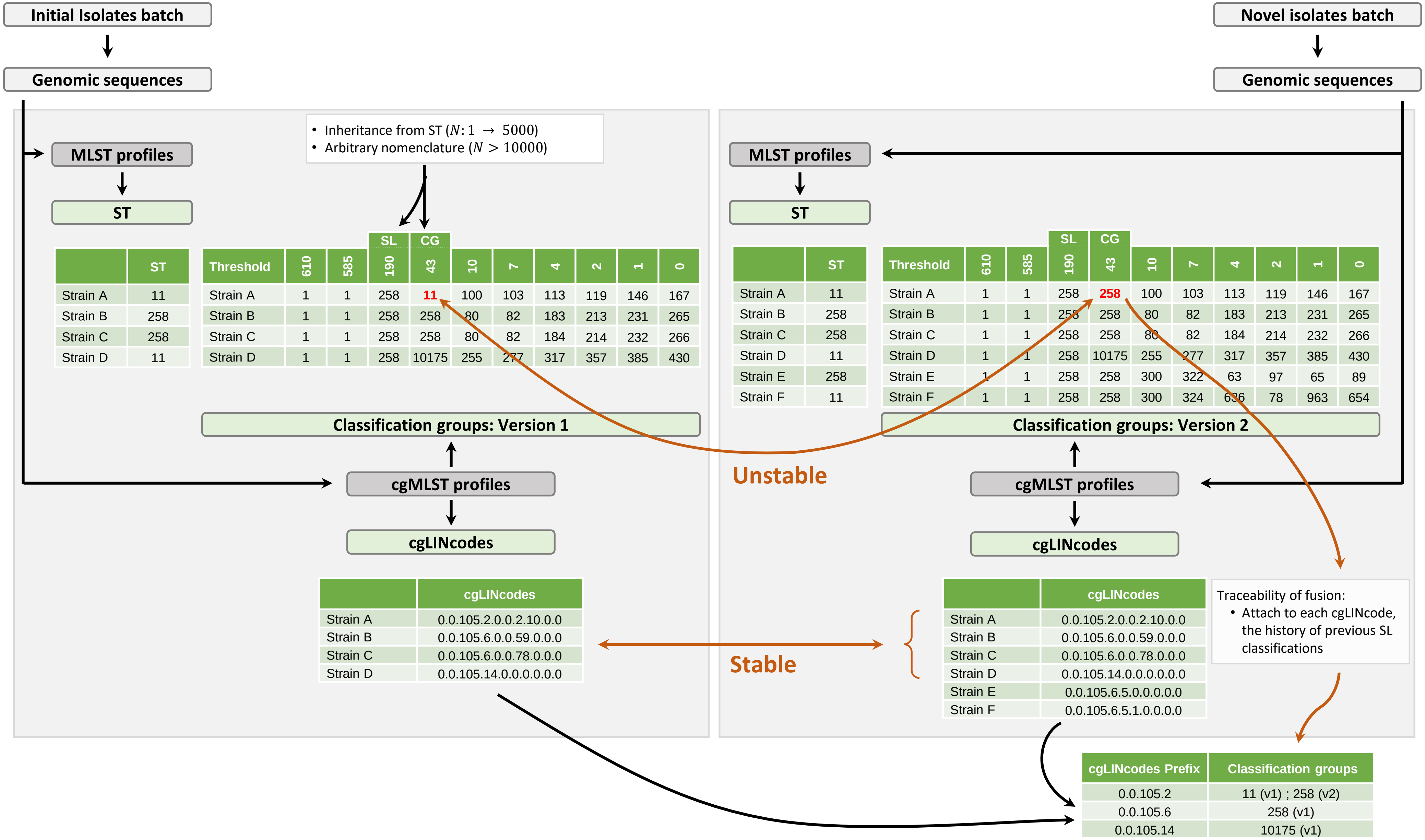


Initial isolates batch:

| | ST |
|---|---|
| Strain A | 11 |
| Strain B | 258 |
| Strain C | 258 |
| Strain D | 11 |

| Threshold | 610 | 585 | 190 (SL) | 43 (CG) | 10 | 7 | 4 | 2 | 1 | 0 |
|---|---|---|---|---|---|---|---|---|---|---|
| Strain A | 1 | 1 | 258 | 11 | 100 | 103 | 113 | 119 | 146 | 167 |
| Strain B | 1 | 1 | 258 | 258 | 80 | 82 | 183 | 213 | 231 | 265 |
| Strain C | 1 | 1 | 258 | 258 | 80 | 82 | 184 | 214 | 232 | 266 |
| Strain D | 1 | 1 | 258 | 10175 | 255 | 277 | 317 | 357 | 385 | 430 |

| | cgLINcodes |
|---|---|
| Strain A | 0.0.105.2.0.0.2.10.0.0 |
| Strain B | 0.0.105.6.0.0.59.0.0.0 |
| Strain C | 0.0.105.6.0.0.78.0.0.0 |
| Strain D | 0.0.105.14.0.0.0.0.0.0 |

Novel isolates batch:

| | ST |
|---|---|
| Strain A | 11 |
| Strain B | 258 |
| Strain C | 258 |
| Strain D | 11 |
| Strain E | 258 |
| Strain F | 11 |

| Threshold | 610 | 585 | 190 (SL) | 43 (CG) | 10 | 7 | 4 | 2 | 1 | 0 |
|---|---|---|---|---|---|---|---|---|---|---|
| Strain A | 1 | 1 | 258 | 258 | 100 | 103 | 113 | 119 | 146 | 167 |
| Strain B | 1 | 1 | 258 | 258 | 80 | 82 | 183 | 213 | 231 | 265 |
| Strain C | 1 | 1 | 258 | 258 | 80 | 82 | 184 | 214 | 232 | 266 |
| Strain D | 1 | 1 | 258 | 10175 | 255 | 277 | 317 | 357 | 385 | 430 |
| Strain E | 1 | 1 | 258 | 258 | 300 | 322 | 63 | 97 | 65 | 89 |
| Strain F | 1 | 1 | 258 | 258 | 300 | 324 | 636 | 78 | 963 | 654 |

| | cgLINcodes |
|---|---|
| Strain A | 0.0.105.2.0.0.2.10.0.0 |
| Strain B | 0.0.105.6.0.0.59.0.0.0 |
| Strain C | 0.0.105.6.0.0.78.0.0.0 |
| Strain D | 0.0.105.14.0.0.0.0.0.0 |
| Strain E | 0.0.105.6.5.0.0.0.0.0 |
| Strain F | 0.0.105.6.5.1.0.0.0.0 |

| cgLINcodes Prefix | Classification groups |
|---|---|
| 0.0.105.2 | 11 (v1) ; 258 (v2) |
| 0.0.105.6 | 258 (v1) |
| 0.0.105.14 | 10175 (v1) |

**Figure S12. Distribution of the phylogroup homogeneity index**

Each graph is based on the population of strains belonging to the indicated phylogoup. The Y-axis correspond to the number of strains (genomes) ; the X-axis corresponds to the percentage of alleles per cgMLST profile attributed to the corresponding phylogroup (see Methods). An additional panel provides details for phylogroup Kp1. Red vertical lines indicate the threshold used to define genomes as ‘hybrids’ (none were defined in phylogroups Kp5, Kp6 and Kp7).

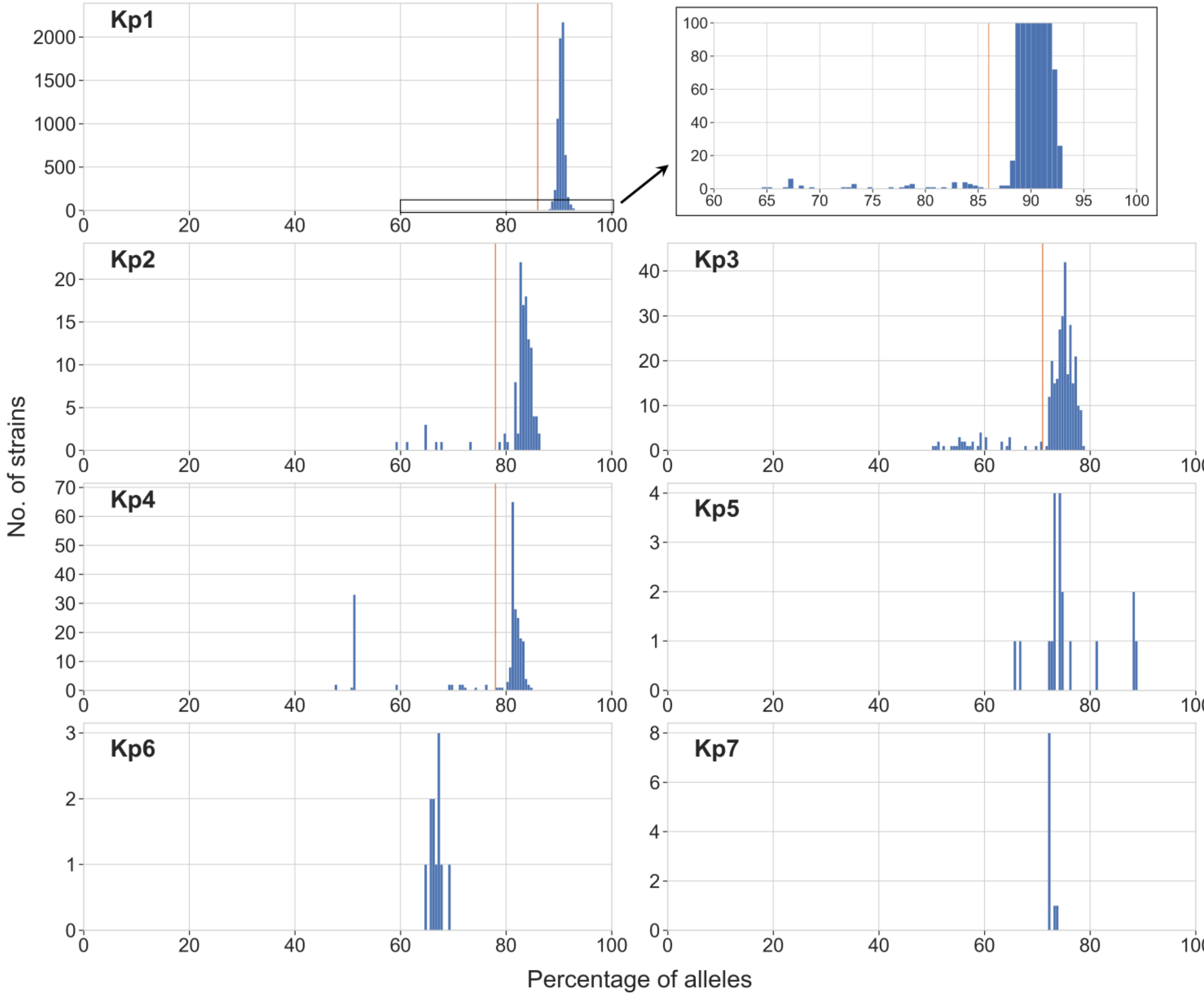

#### Figure S13. Principle of cgLIN code implementation

The principle is derived from Marakeby et al. (Marakeby et al., 2014), using cgMLST distances instead of ANI. Step 1: the code is initialized with the first genome being assigned the value "0" at all positions. Step 2: the encoding for an incoming genome is based on the closest genome already encoded In the chosen example, when the genome of strain B is submitted, the only genome in the database is that of strain A. Therefore, the cgLIN code of strain B is assigned based on the cgLIN code of strain A. By the time strain C is submitted, strains A and B are in the database. Based on the distance between the cgMLST profile of strain C and those present in the database, strain B is found to be the most similar. Therefore, the cgLIN code of strain C is assigned based on the cgLIN code of strain B and a value corresponding to the value in the cgLIN code of strain B, plus 1 (here, 0+1=1) is attributed to the bin corresponding to the lowest bin with a higher identity threshold that the similarity between B and C. Downstream bins are attributed “0” in the cgLIN code of strain C. Upstream bins are attributed the same value as for strain B. When D is introduced, it is coded according to strain C, its closest genome already encoded.

### Figure S13. Principle of cgLIN code implementation

#### Step 1: initialization

Database

| Number of mismatch threshold | 610 | 585 | 190 | 43 | 10 | 7 | 4 | 2 | 1 | 0 |
|---|---|---|---|---|---|---|---|---|---|---|
| Number of match threshold | 19 | 44 | 439 | 586 | 619 | 622 | 625 | 627 | 628 | 629 |
| % identity threshold | 3.0207 | 6.9952 | 69.7933 | 93.1638 | 98.4102 | 98.8871 | 99.3641 | 99.6820 | 99.8410 | 100.000 |
| Strain A | 0 | 0 | 0 | 0 | 0 | 0 | 0 | 0 | 0 | 0 |

#### Step 2: incrementation

Assignment of the cgLINcode of next strain

| Strain | Closest genome | Number of mismatch (β) | Number of loci called in both strains (Ω) | % identity |
|---|---|---|---|---|
| Strain B | Strain A | 8 | 621 | $\frac{\Omega - \beta}{\Omega}$ = 98.711 |

Lowest bin with higher identity threshold

Database

| Number of mismatch threshold | 610 | 585 | 190 | 43 | 10 | 7 | 4 | 2 | 1 | 0 |
|---|---|---|---|---|---|---|---|---|---|---|
| % identity threshold | 3.0207 | 6.9952 | 69.7933 | 93.1638 | 98.4102 | 98.8871 | 99.3641 | 99.6820 | 99.8410 | 100.000 |
| Strain A | 0 | 0 | 0 | 0 | 0 | 0 | 0 | 0 | 0 | 0 |
| Strain B | 0 | 0 | 0 | 0 | 0 | 1 | 0 | 0 | 0 | 0 |

Assignment of the cgLINcode of next strain

| Strain | Closest genome | Number of mismatch (β) | Number of loci called in both strains (Ω) | % identity |
|---|---|---|---|---|
| Strain C | Strain B | 2 | 625 | $\frac{\Omega - \beta}{\Omega}$ = 99.680 |

Database

| Number of mismatch threshold | 610 | 585 | 190 | 43 | 10 | 7 | 4 | 2 | 1 | 0 |
|---|---|---|---|---|---|---|---|---|---|---|
| % identity threshold | 3.0207 | 6.9952 | 69.7933 | 93.1638 | 98.4102 | 98.8871 | 99.3641 | 99.6820 | 99.8410 | 100.000 |
| Strain A | 0 | 0 | 0 | 0 | 0 | 0 | 0 | 0 | 0 | 0 |
| Strain B | 0 | 0 | 0 | 0 | 0 | 1 | 0 | 0 | 0 | 0 |
| Strain C | 0 | 0 | 0 | 0 | 0 | 1 | 0 | 1 | 0 | 0 |

Assignment of the cgLINcode of next strain

| Strain | Closest genome | Number of mismatch (β) | Number of loci called in both strains (Ω) | % identity |
|---|---|---|---|---|
| Strain D | Strain C | 4 | 627 | $\frac{\Omega - \beta}{\Omega}$ = 99.362 |

Database

| Number of mismatch threshold | 610 | 585 | 190 | 43 | 10 | 7 | 4 | 2 | 1 | 0 |
|---|---|---|---|---|---|---|---|---|---|---|
| % identity threshold | 3.0207 | 6.9952 | 69.7933 | 93.1638 | 98.4102 | 98.8871 | 99.3641 | 99.6820 | 99.8410 | 100.000 |
| Strain A | 0 | 0 | 0 | 0 | 0 | 0 | 0 | 0 | 0 | 0 |
| Strain B | 0 | 0 | 0 | 0 | 0 | 1 | 0 | 0 | 0 | 0 |
| Strain C | 0 | 0 | 0 | 0 | 0 | 1 | 0 | 1 | 0 | 0 |
| Strain D | 0 | 0 | 0 | 0 | 0 | 1 | 1 | 0 | 0 | 0 |

#### Figure S14. cgLIN codes implementation for nearly-identical cgMLST profiles

This figure illustrates the principle of cgLIN code implementation, when the genomes are very closely related to each other, as well as when there are missing data in the cgMLST profiles. The encoding process is similar to the general case (Figure S13). In case of complete identity and no missing data, cgLIN codes are exactly identical (see strains X and W). In cases where there is identity at all called loci, but with some loci being uncalled, the similarity would still be 100% resulting in identical cgLIN codes (not shown). In case of a single mismatch and no missing data, the cgLIN codes differ only at their last position (strains Y and X). In case of a single or several mismatches, the effect of missing data at some other loci will be to decrease the similarity ratio compared to the case where no data would be missing at other loci (strain Z therefore differs from Y already at the penultimate bin, even with a single allelic mismatch).

### Figure S14. cgLIN codes implementation for nearly-identical cgMLST profiles

**Sequential assignment of cgLIN codes of strains**

**Database**

**Strain W**

| Number of mismatch threshold | 610 | 585 | 190 | 43 | 10 | 7 | 4 | 2 | 1 | 0 |
|---|---|---|---|---|---|---|---|---|---|---|
| Number of match threshold | 19 | 44 | 439 | 586 | 619 | 622 | 625 | 627 | 628 | 629 |
| % identity threshold | 3.0207 | 6.9952 | 69.7933 | 93.1638 | 98.4102 | 98.8871 | 99.3641 | 99.6820 | 99.8410 | 100.000 |
| Strain W | 0 | 0 | 396 | 0 | 0 | 0 | 0 | 10 | 0 | 0 |

**Strain X**

| Strain | Closest genome | Number of mismatch (β) | Number of loci called in both strains (Ω) | % identity |
|---|---|---|---|---|
| Strain X | Strain W | 0 | 629 | $\frac{\Omega - \beta}{\Omega} = 100$ |

| Number of mismatch threshold | 610 | 585 | 190 | 43 | 10 | 7 | 4 | 2 | 1 | 0 |
|---|---|---|---|---|---|---|---|---|---|---|
| % identity threshold | 3.0207 | 6.9952 | 69.7933 | 93.1638 | 98.4102 | 98.8871 | 99.3641 | 99.6820 | 99.8410 | 100.000 |
| Strain W | 0 | 0 | 396 | 0 | 0 | 0 | 0 | 10 | 0 | 0 |
| Strain X | 0 | 0 | 396 | 0 | 0 | 0 | 0 | 10 | 0 | 0 |

**Strain Y**

| Strain | Closest genome | Number of mismatch (β) | Number of loci called in both strains (Ω) | % identity |
|---|---|---|---|---|
| Strain Y | Strain X | 1 | 629 | $\frac{\Omega - \beta}{\Omega} = 99.8410$ |

| Number of mismatch threshold | 610 | 585 | 190 | 43 | 10 | 7 | 4 | 2 | 1 | 0 |
|---|---|---|---|---|---|---|---|---|---|---|
| % identity threshold | 3.0207 | 6.9952 | 69.7933 | 93.1638 | 98.4102 | 98.8871 | 99.3641 | 99.6820 | 99.8410 | 100.000 |
| Strain W | 0 | 0 | 396 | 0 | 0 | 0 | 0 | 10 | 0 | 0 |
| Strain X | 0 | 0 | 396 | 0 | 0 | 0 | 0 | 10 | 0 | 0 |
| Strain Y | 0 | 0 | 396 | 0 | 0 | 0 | 0 | 10 | 0 | 1 |

**Strain Z**

| Strain | Closest genome | Number of mismatch (β) | Number of loci called in both strains (Ω) | % identity |
|---|---|---|---|---|
| Strain Z | Strain Y | 1 | $600 \leq \Omega \leq 628$ | $99{,}83 \leq \frac{\Omega - \beta}{\Omega} \leq 99.840$ |

| Number of mismatch threshold | 610 | 585 | 190 | 43 | 10 | 7 | 4 | 2 | 1 | 0 |
|---|---|---|---|---|---|---|---|---|---|---|
| % identity threshold | 3.0207 | 6.9952 | 69.7933 | 93.1638 | 98.4102 | 98.8871 | 99.3641 | 99.6820 | 99.8410 | 100.000 |
| Strain W | 0 | 0 | 396 | 0 | 0 | 0 | 0 | 10 | 0 | 0 |
| Strain X | 0 | 0 | 396 | 0 | 0 | 0 | 0 | 10 | 0 | 0 |
| Strain Y | 0 | 0 | 396 | 0 | 0 | 0 | 0 | 10 | 0 | 1 |
| Strain Z | 0 | 0 | 396 | 0 | 0 | 0 | 0 | 10 | 1 | 0 |

#### Figure S15. Step-by-step illustration of the taxonomic inheritance algorithm

Given 12 hypothetical clonal group (CG, named A-L) and 16 sequence types (ST, labelled from 1 to 16), a weighted bipartite graph *G* is shown, as well as the use of *G* to label the CG based on their relation to their related ST. For eacj step illustration, the corresponding lines of the algorithm pseudo-code (see SupMat) are specified (bottom of each sub-figure).

Figure S15. (continued)

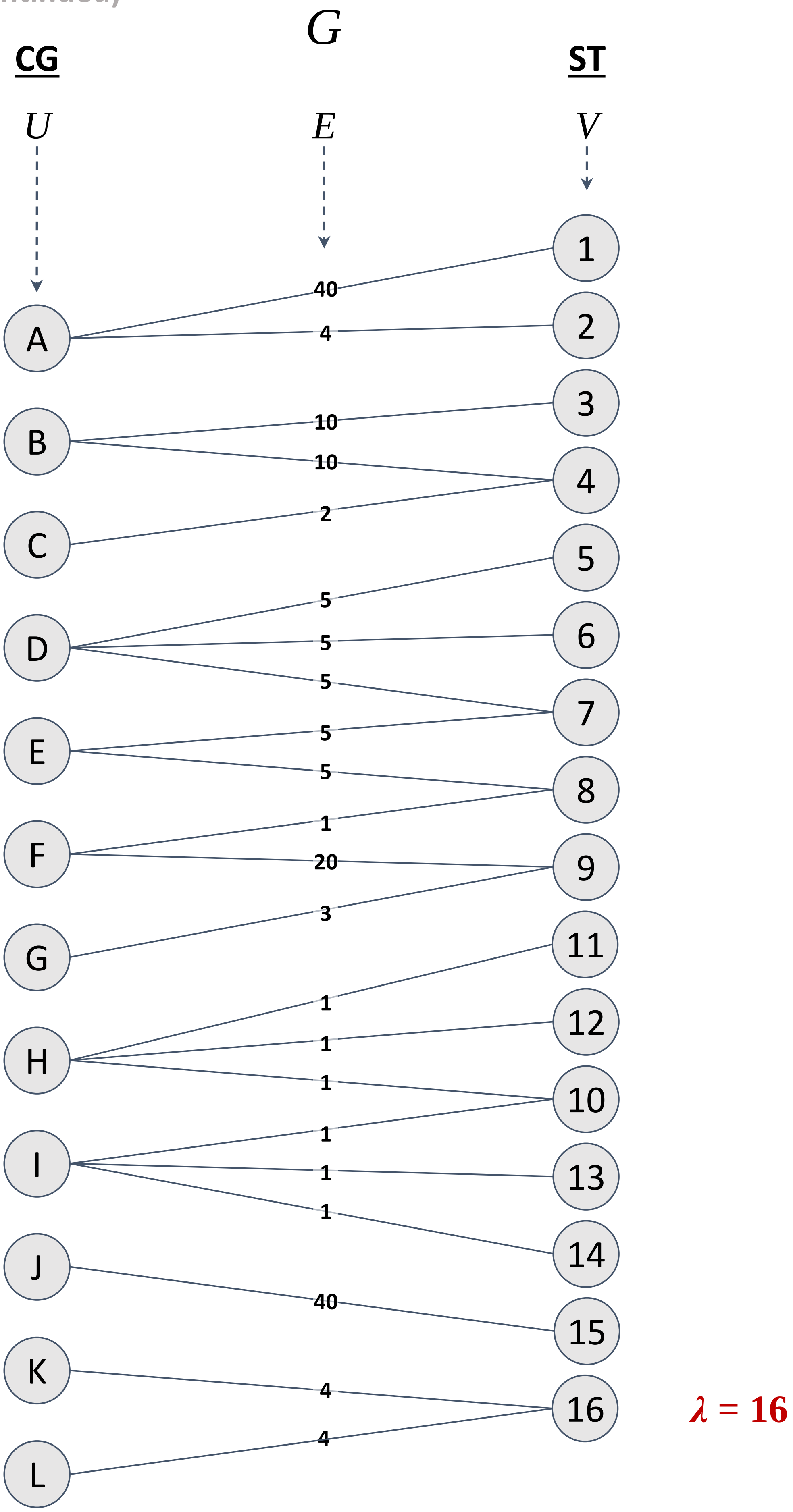


(a)

Figure S15. (continued)

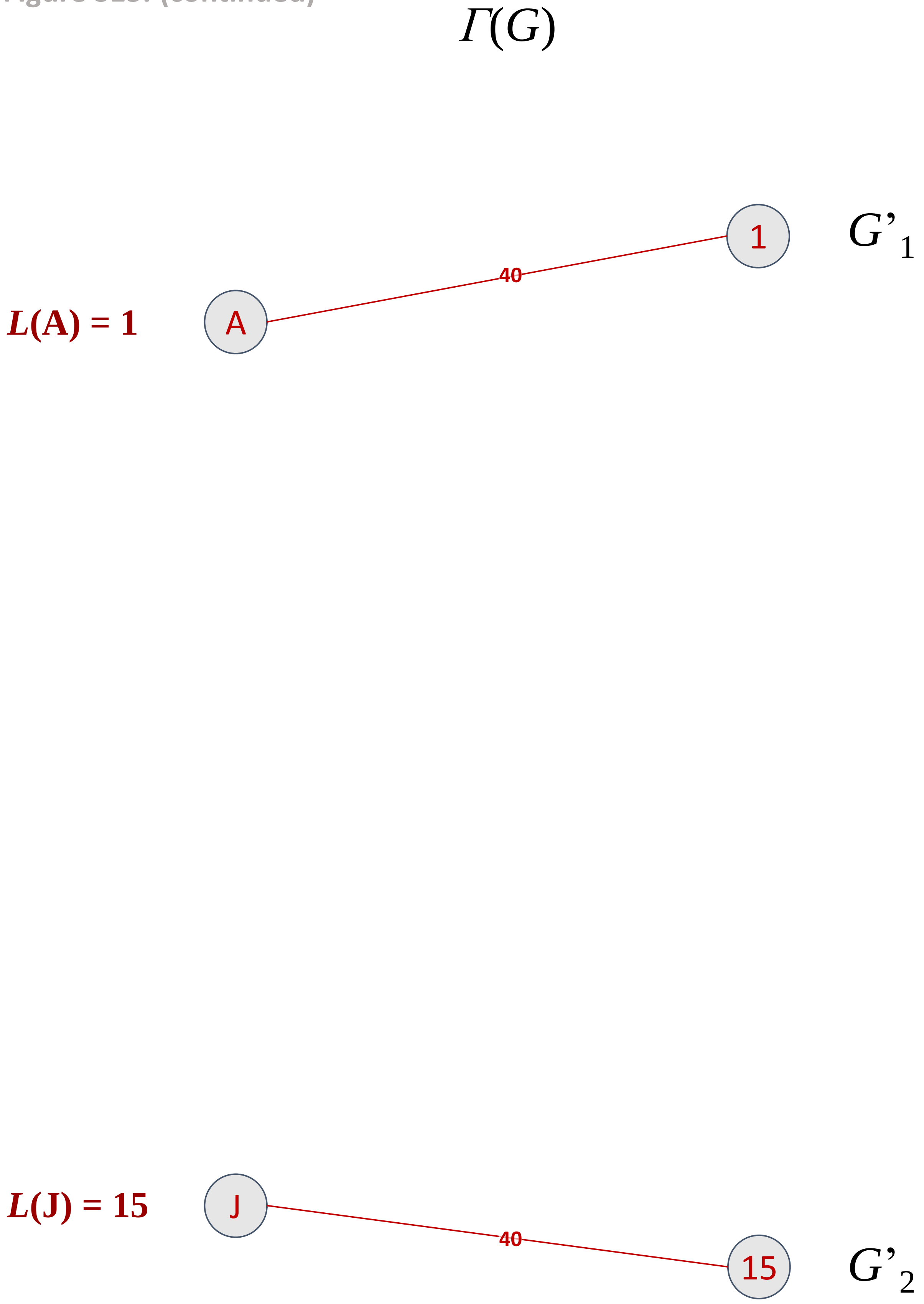


(c), (d), (e) and (j)

Figure S15. (continued)

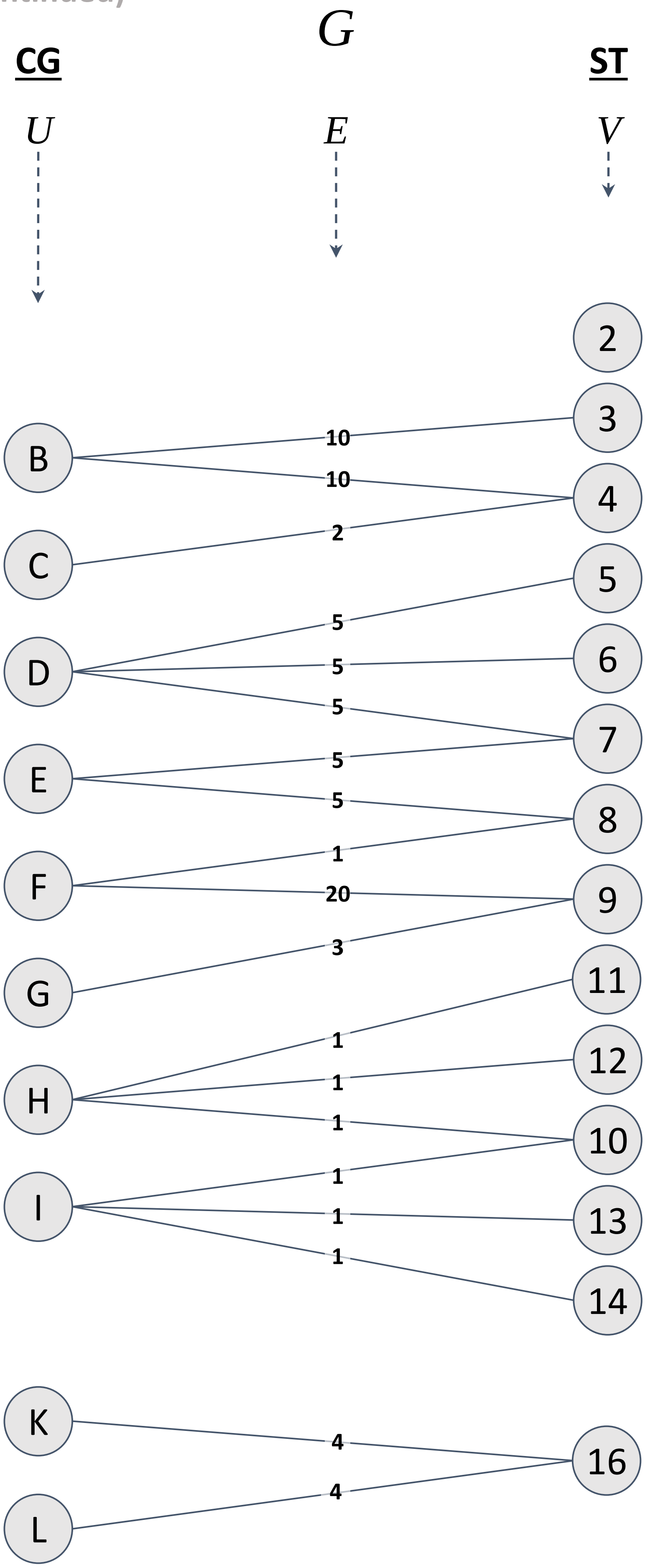


(i)

Figure S15. (continued)

$\Gamma(G)$

**$L$(F) = 9**

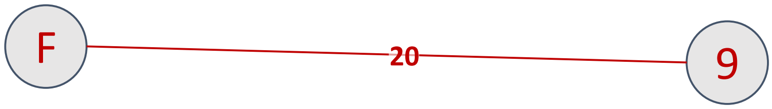


(c), (d), (e) and (j)

Figure S15. (continued)

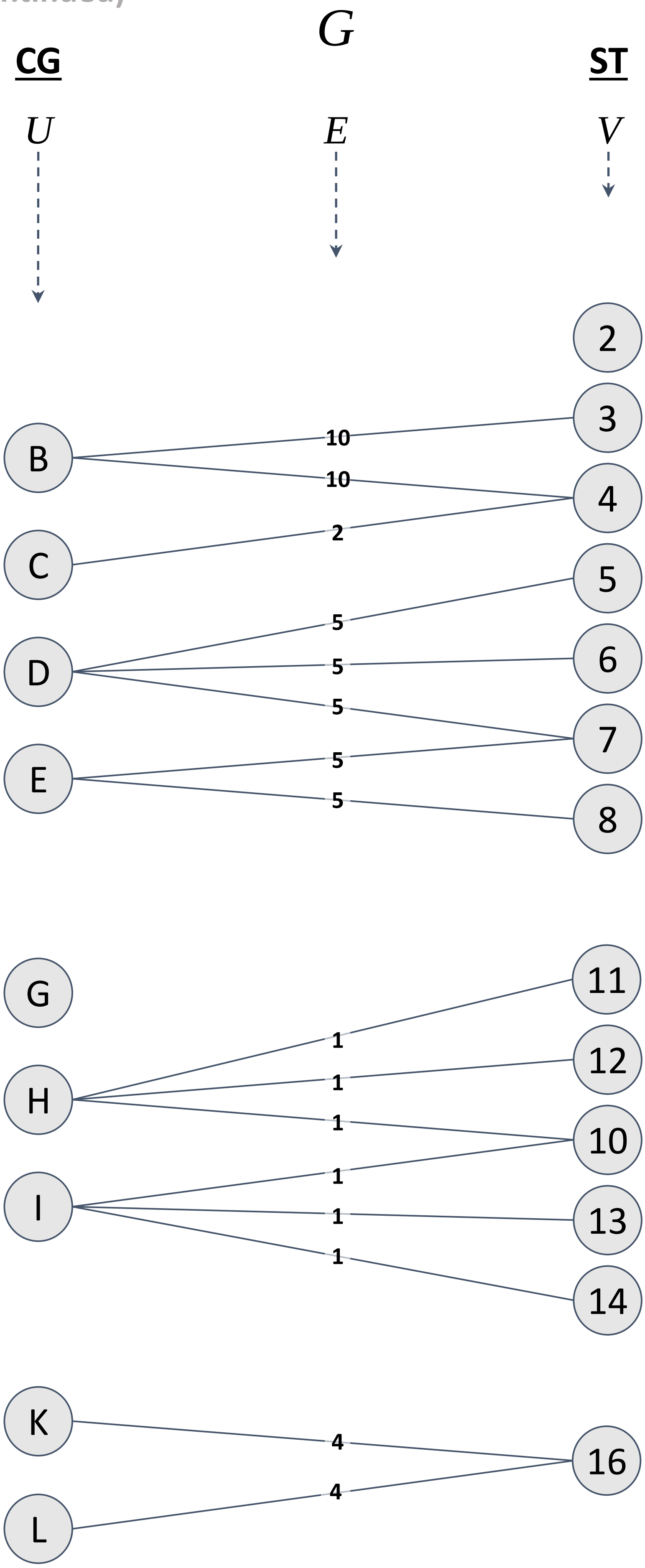


(i)

Figure S15. (continued)

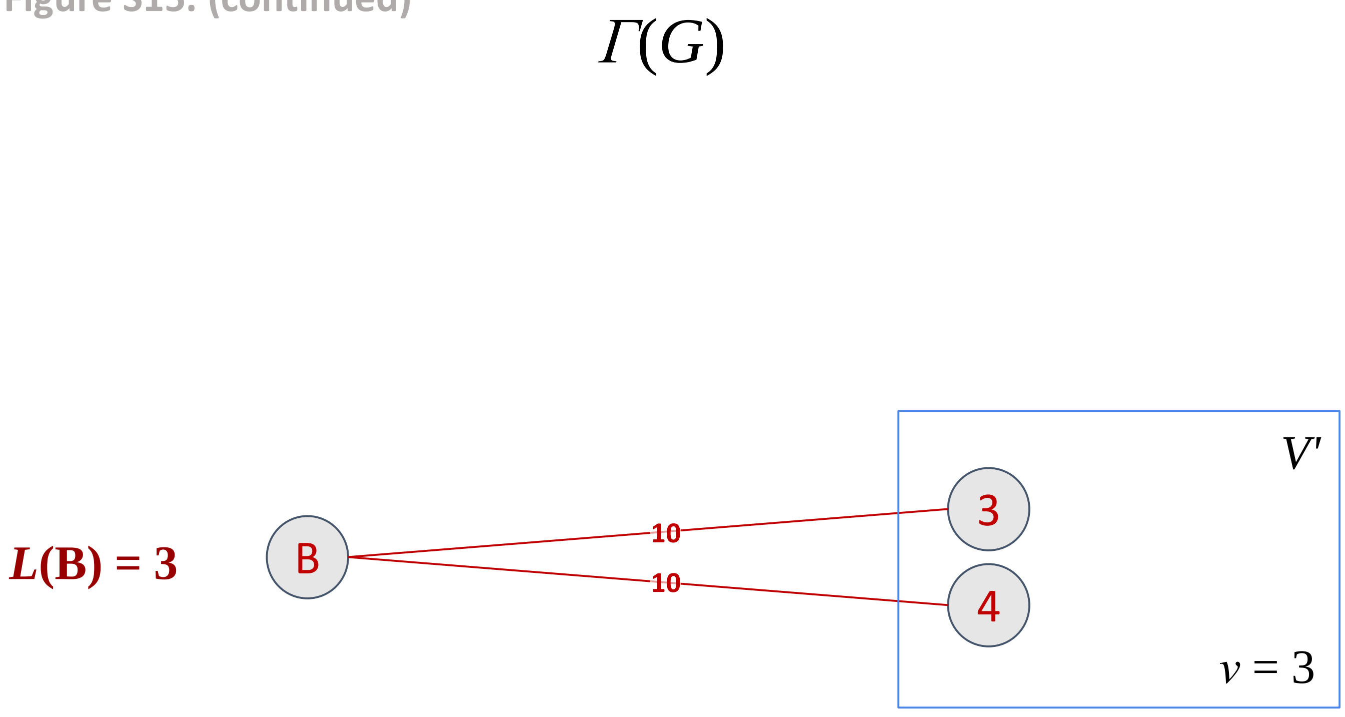


(c), (d), (e) and (j)

Figure S15. (continued)

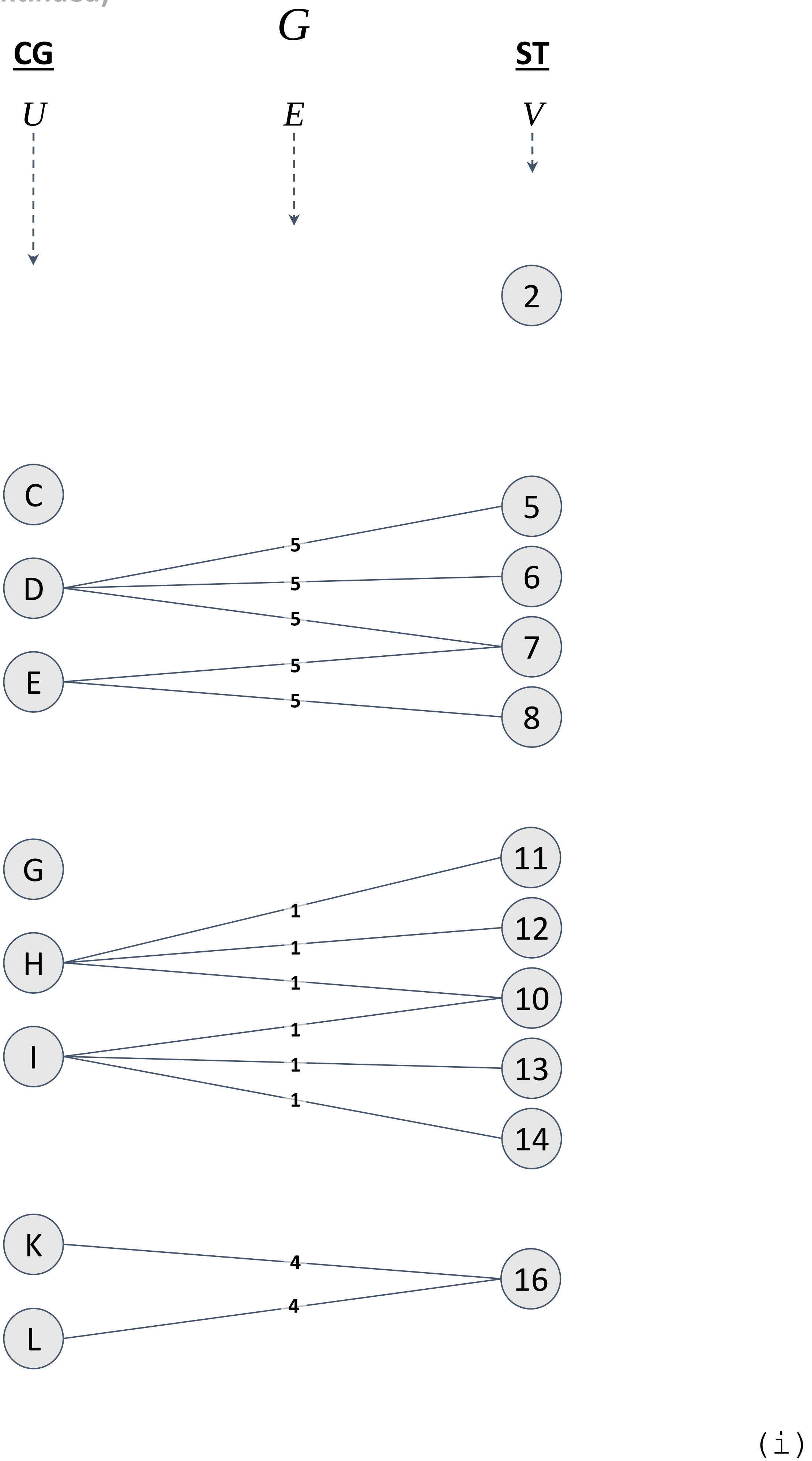

Figure S15. (continued)

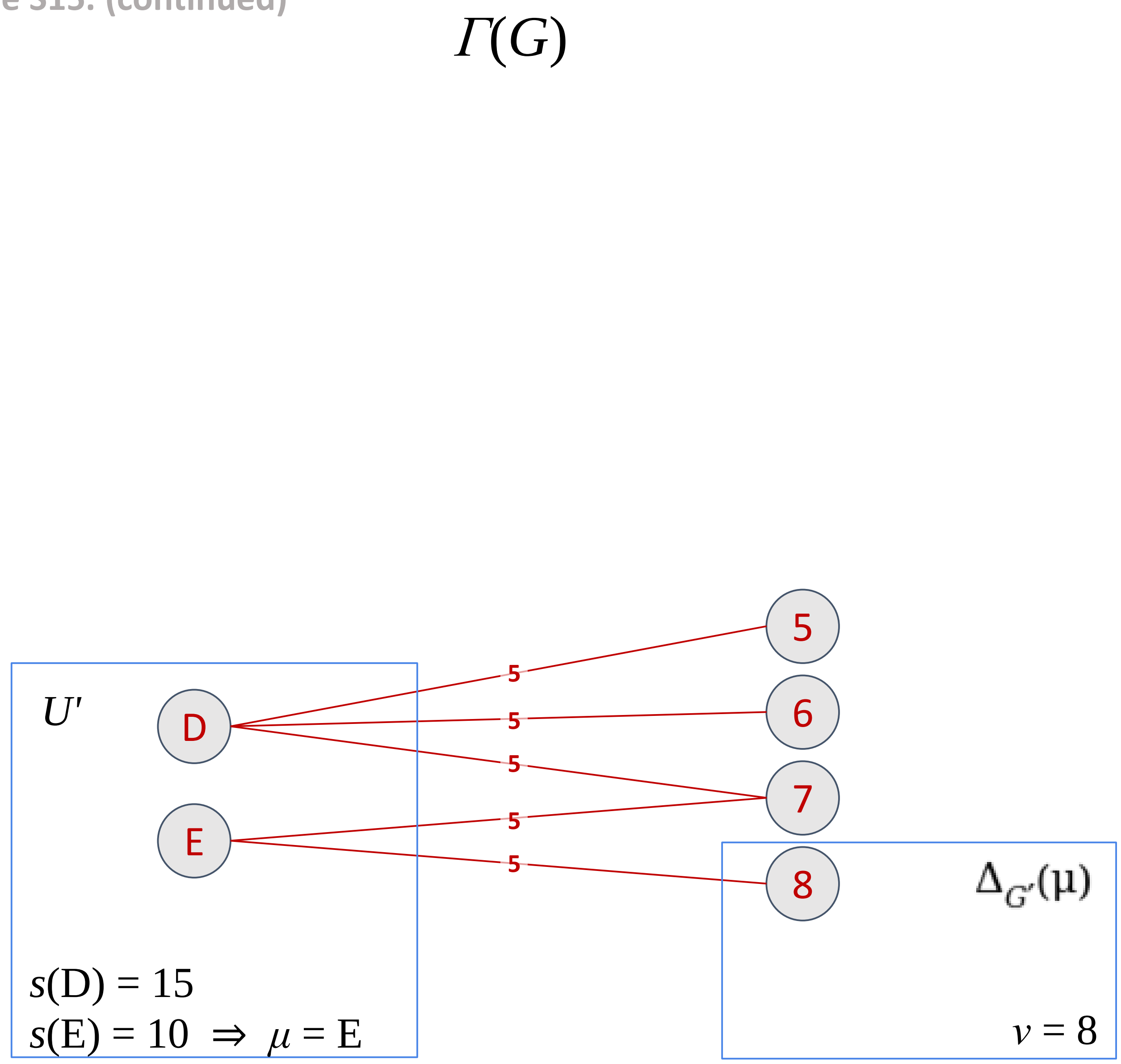


**$L(\text{E}) = 8$**

`(c), (f), (i)` and `(j)`

Figure S15. (continued)

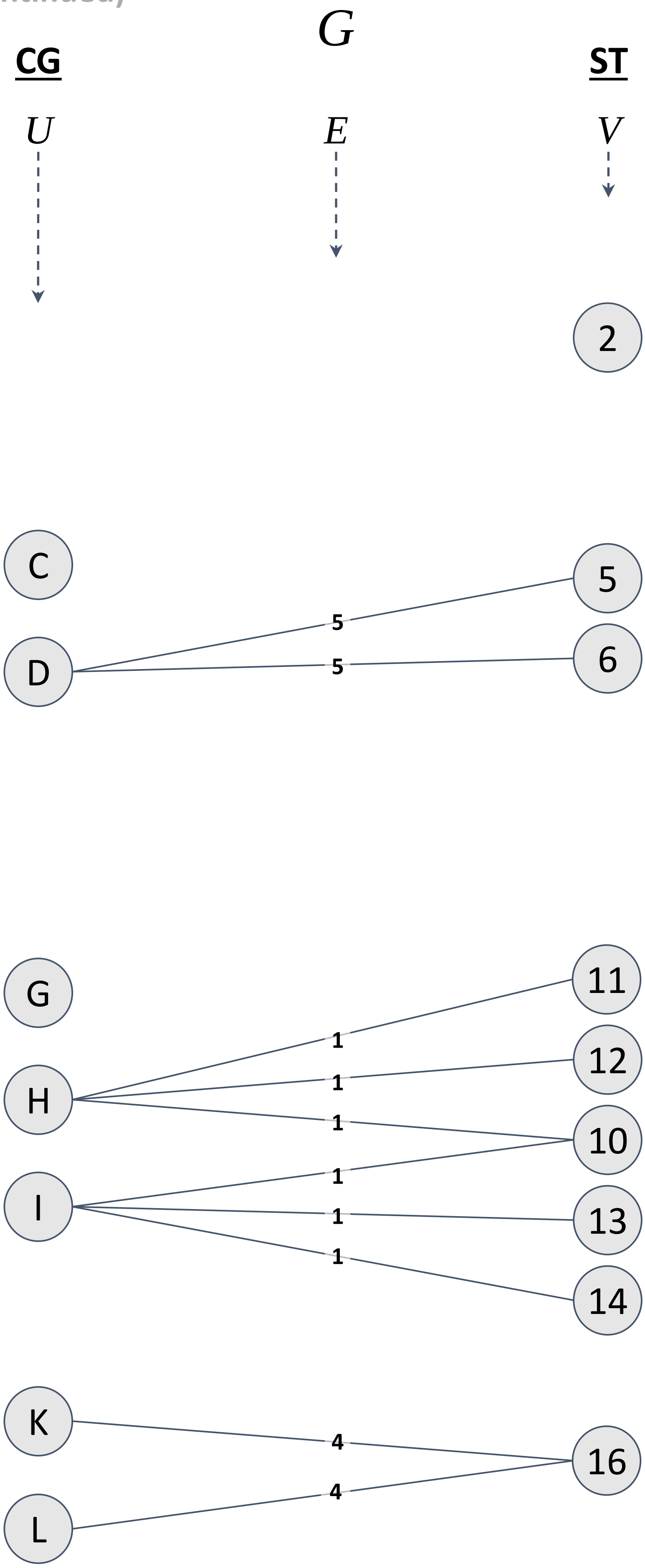


(i)

Figure S15. (continued)

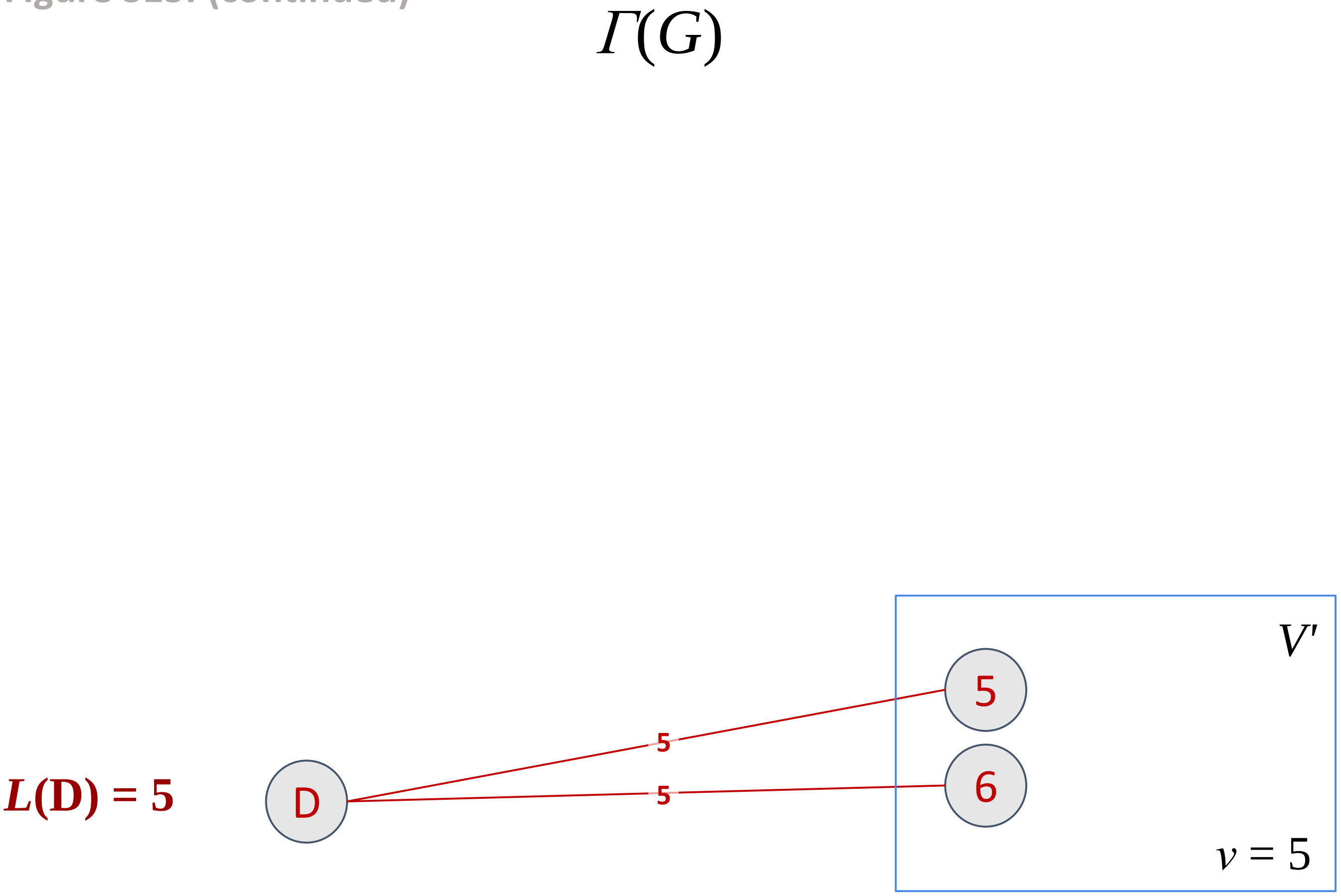


(c), (d), (e) and (j)

Figure S15. (continued)

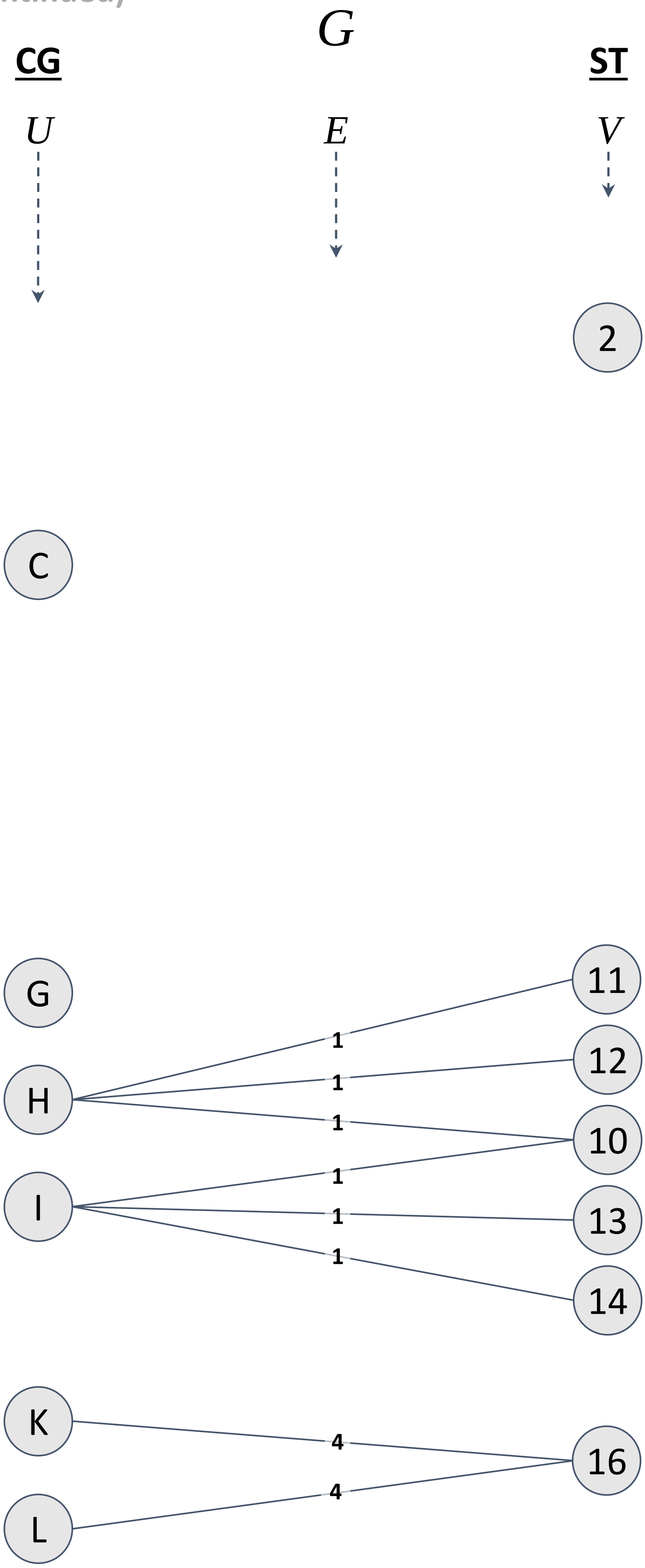


(i)

Figure S15. (continued)

$\Gamma(G)$

$L(\mathrm{K}) = 16$

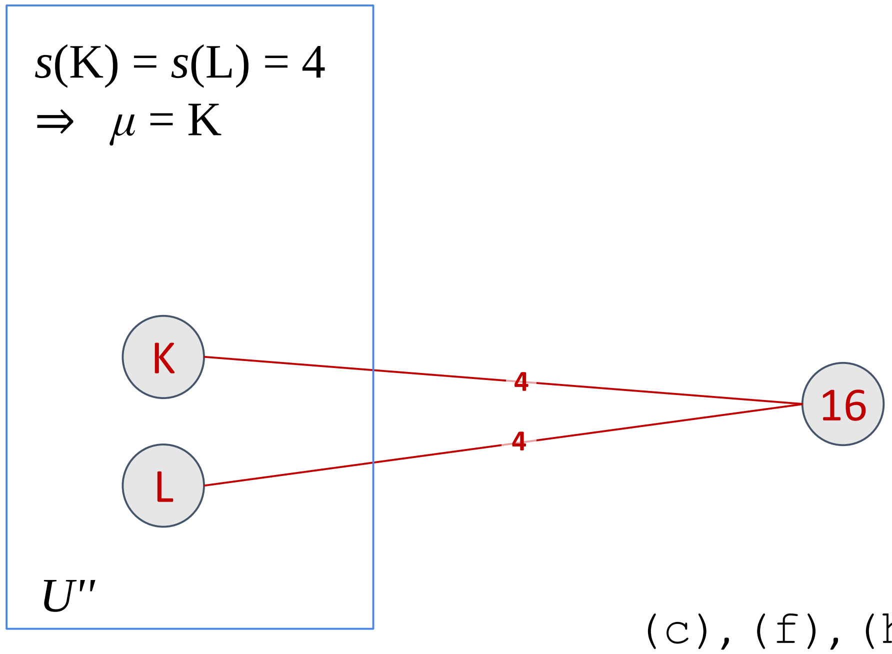


(c), (f), (h), (i) and (j)

Figure S15. (continued)

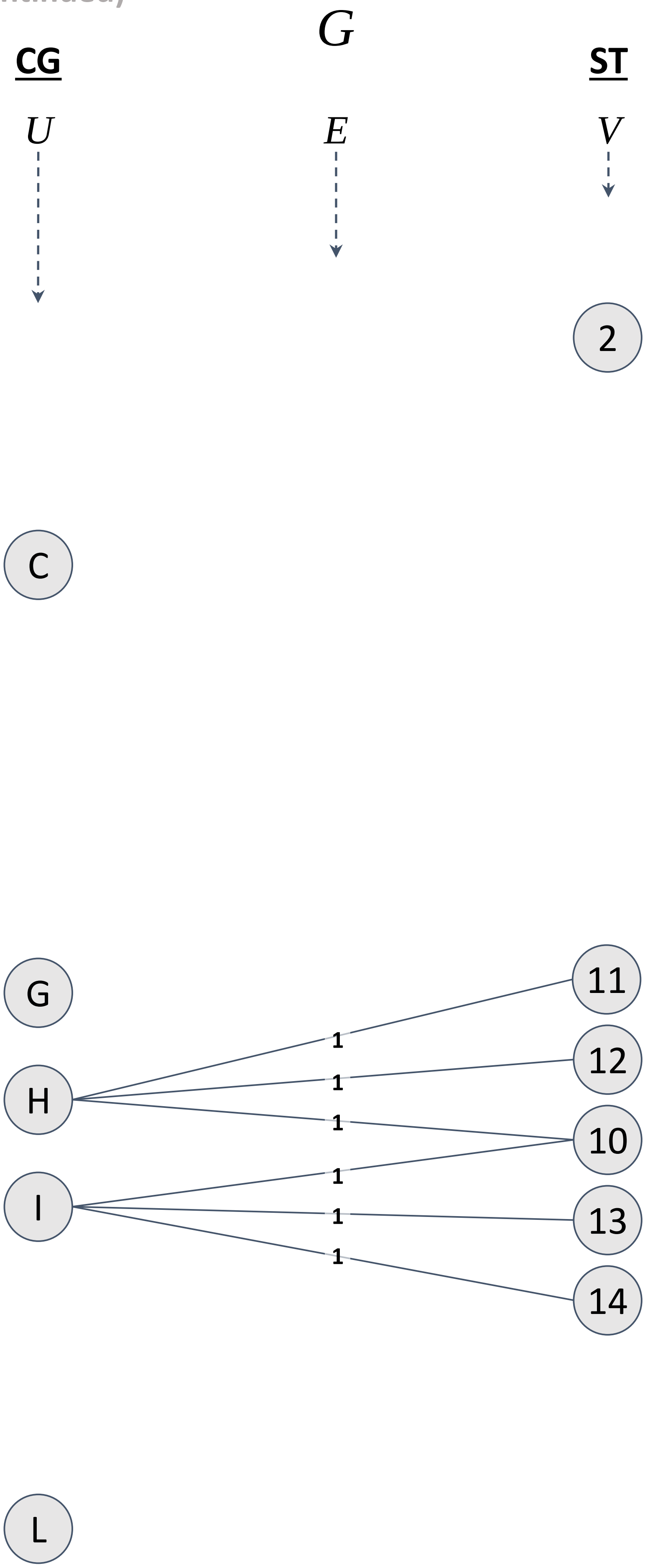


(i)

$\Gamma(G)$

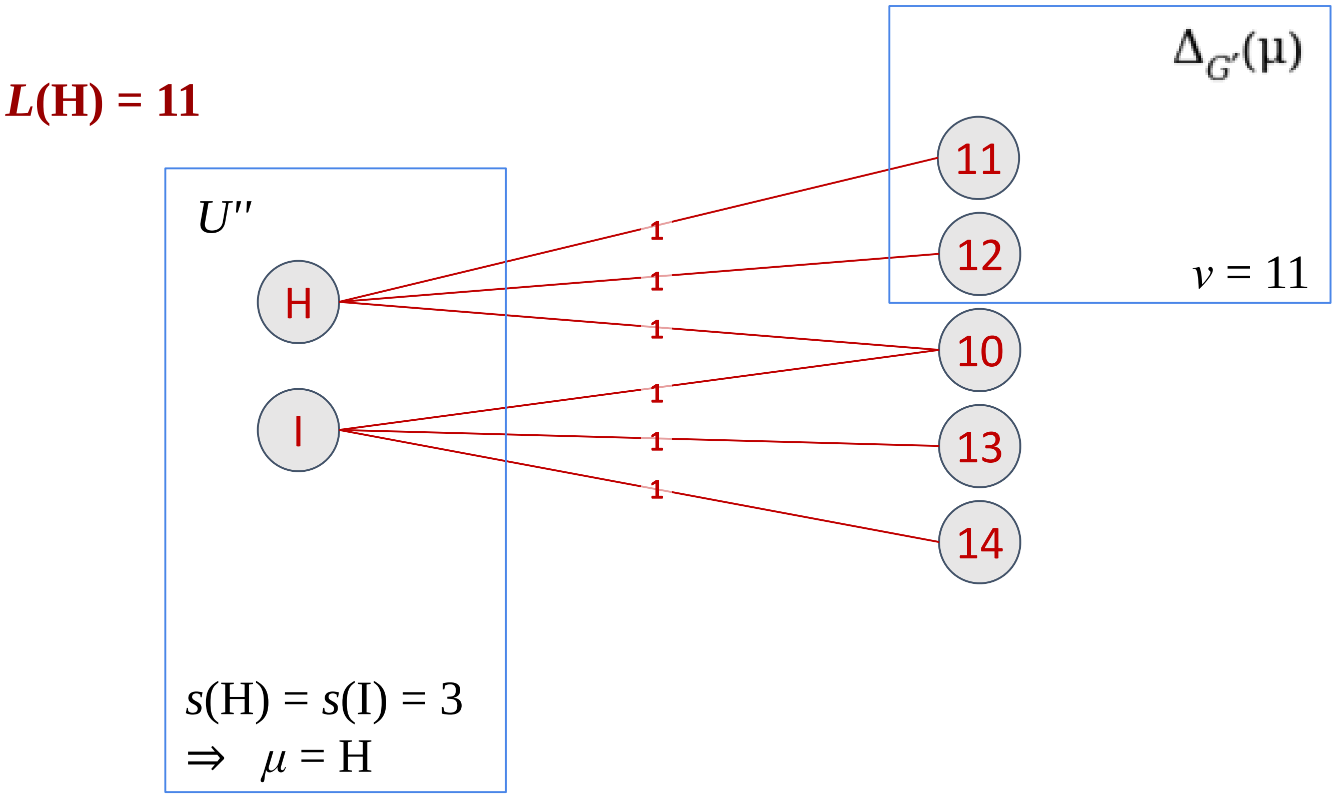


(c), (f), (h), (i) and (j)

Figure S15. (continued)

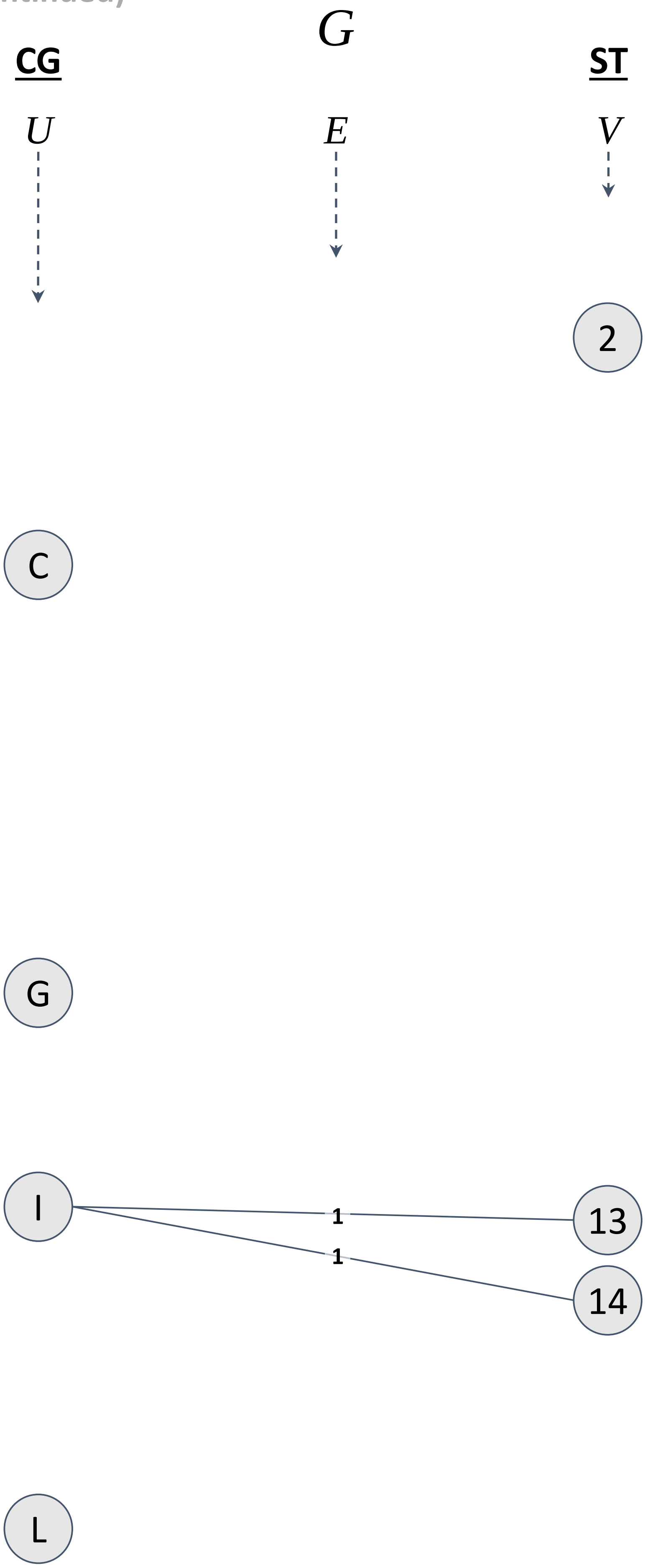


(i)

Figure S15. (continued)

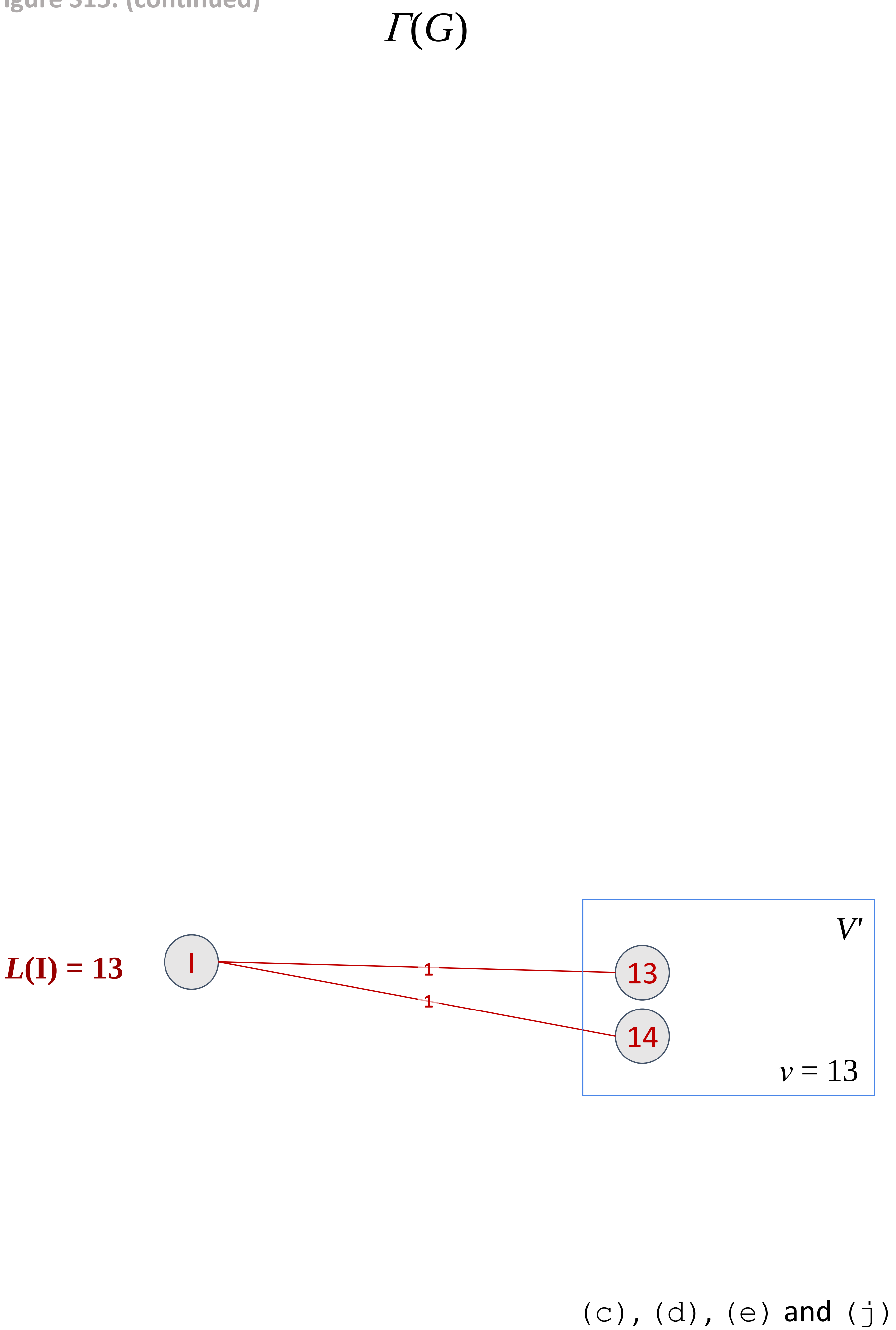

Figure S15. (continued)

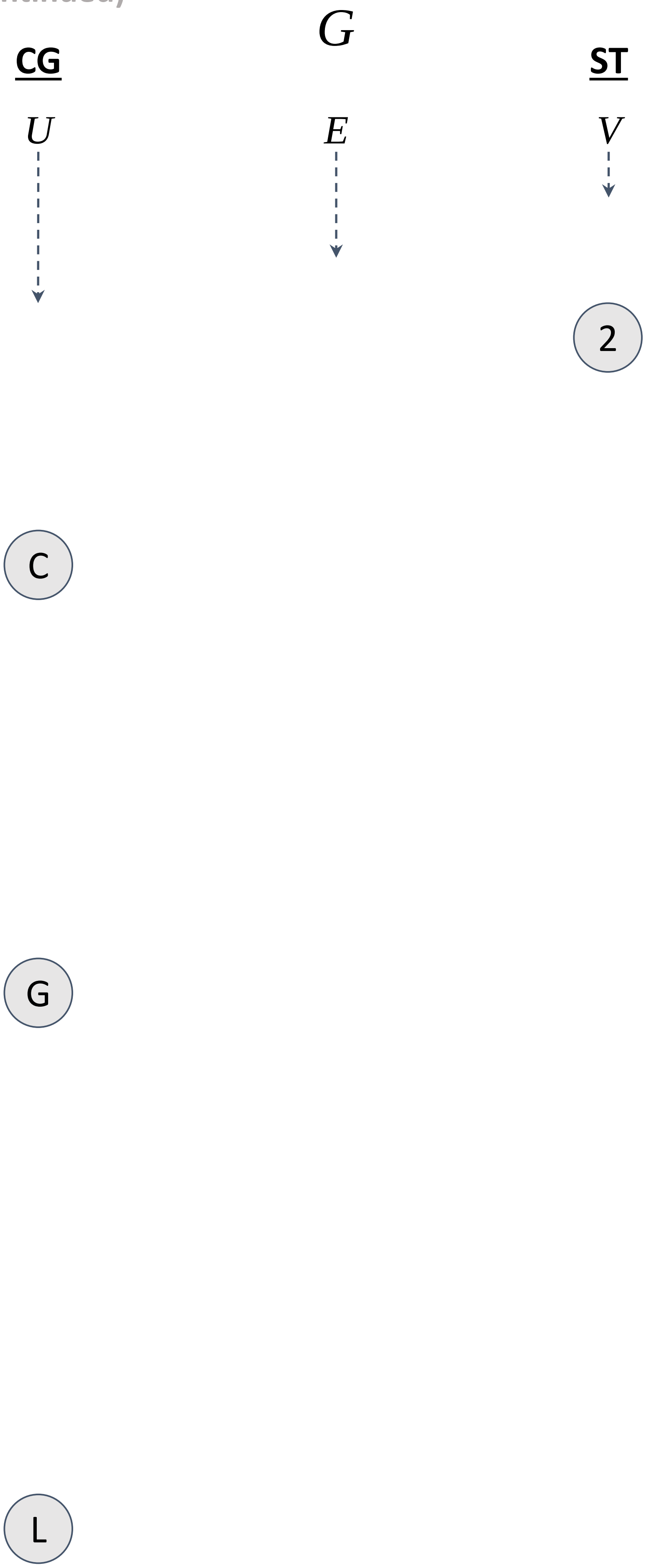


(i)

Figure S15. (continued)

$\lambda = 16$

$L(C) = 17$ C

$L(G) = 18$ G

$L(L) = 19$ L

(a), (l), (m) and (n)
